# Recovering signatures of archaic introgression using ancestral recombination graphs

**DOI:** 10.64898/2026.03.03.709416

**Authors:** Yulin Zhang, Arjun Biddanda, Sarah A. Johnson, Colm O’Dushlaine, Priya Moorjani

## Abstract

Neanderthal and Denisovan genomes have reshaped our understanding of archaic introgression. Yet, the limited number of archaic genomes sequenced and the reliance on unadmixed outgroups have left much of this history unresolved. We introduce TRACE, a method to identify archaic ancestry using features of ancestral recombination graphs inferred from contemporary genomes alone. Simulations show that TRACE reliably detects archaic introgression without requiring archaic genomes or unadmixed outgroups. Applied to 1000 Genomes data, TRACE recovers known Neanderthal and Denisovan introgression and reveals signals of ghost admixture from previously uncharacterized hominins in both Africans and non-Africans. Strikingly, ghost ancestry persists in Neanderthal and Denisovan ancestry deserts, challenging their interpretation as *Homo sapiens*-specific regions. In Oceanians, TRACE finds deep lineages enriched in Denisovan––and not Neanderthal––regions, supporting a model of super-archaic gene flow. TRACE provides a scalable framework for mapping the legacy of archaic introgression in the absence of archaic genome sequences.

## Introduction

The sequencing of the Neanderthal and Denisovan genomes has transformed our understanding of human evolution, revealing evidence for gene flow between modern humans and archaic hominins (*1–5*). Archaic ancestry has had a major impact on human adaptation and disease, contributing to a range of traits such as skin pigmentation, high altitude adaptation, and immune function (*6–12*). To date, however, only four high coverage archaic genomes have been published––three Neanderthals and one Denisovan––all from Eurasia. Thus, our knowledge of the evolutionary history and impact of archaic ancestry outside Eurasia and at deeper timescales remains incomplete (*1–3, 5*).

Several studies have posited that gene flow from other “ghost” lineages—unknown archaic hominins for whom no genomic sequences currently exist—may have occurred both within and outside Africa (*13–26*). Genetic analyses of present-day African populations have identified highly divergent haplotypes that cannot be explained by known demographic histories, hinting at introgression from unknown hominins (*13–25*). At deeper timescales, analysis of the Altai Denisovan genome suggests that they harbor ancestry from a deeply divergent population, referred to as “super-archaic”, that split from modern humans around 0.9–1.4 million years ago (Mya) (*2, 3, 27*). These histories, however, remain poorly understood as most available methods for studying archaic introgression rely on either sequenced archaic genomes (*28*) or an outgroup population (without archaic ancestry, (*14, 29–31*)) or both (*32, 33*). In both these cases, no archaic reference genomes exist––the oldest ancient genome from Africa is less than 20,000 years old (*22*) and recovering hominin DNA older than a million years is unlikely, particularly outside of permafrost (*34*).

With the availability of new methods to reconstruct ancestral recombination graphs (ARGs) (*35– 38*), it is now possible to infer the full evolutionary history of a set of sequences in a computationally tractable manner. An ARG provides a complete record of all the coalescence and recombination events and specifies a complete genealogy at each position in the genome (*39, 40*). This framework provides insights into changes in population size over time and enables the inference of historical relationships and gene flow among individuals (*41, 42*). Importantly, gene flow from deeply divergent lineages is expected to leave distinct signatures in the ARG––such as unusually deep coalescent events––that can be used to detect introgression, even in the absence of archaic reference genomes (*4*).

We introduce TRACE (*TR*acking *A*rchaic *C*ontributions via ARG *E*stimation), a novel method for identifying footprints of archaic ancestry in modern human genomes by leveraging features of ARGs constructed from contemporary genomes alone, requiring neither an archaic reference genome nor an unadmixed outgroup population. We perform extensive simulations to characterize the reliability of TRACE under a range of demographic scenarios, using true and inferred ARGs. We then apply TRACE to whole genome sequences of individuals from worldwide populations to reconstruct the evolutionary history and legacy of archaic gene flow in modern humans.

## Results

### TRACE: A novel method for identifying archaic gene flow using ancestral recombination graphs

TRACE infers archaic ancestry by leveraging genealogical information encoded in ARGs. The central idea is that archaic introgression leaves two characteristic signatures in the sequence of marginal trees within an ARG: long branches and long haplotypes. First, long branches arise because introgression from a deeply divergent population introduces lineages that coalesce much further back in time than non-introgressed lineages (*43*). In the ARG, this appears as branches with deep coalescence times that span the interval between the divergence time of the archaic and modern lineages (*T*_*archaic*_) and the time of the admixture event (*T*_*admix*_) (**Fig. 1A**, (*38*)). Second, because admixture is more recent than divergence, introgressed segments are expected to be longer than those arising from incomplete lineage sorting (ILS), as less time has elapsed since admixture for recombination to break down the ancestry tracts (*44*).

**Fig 1.**
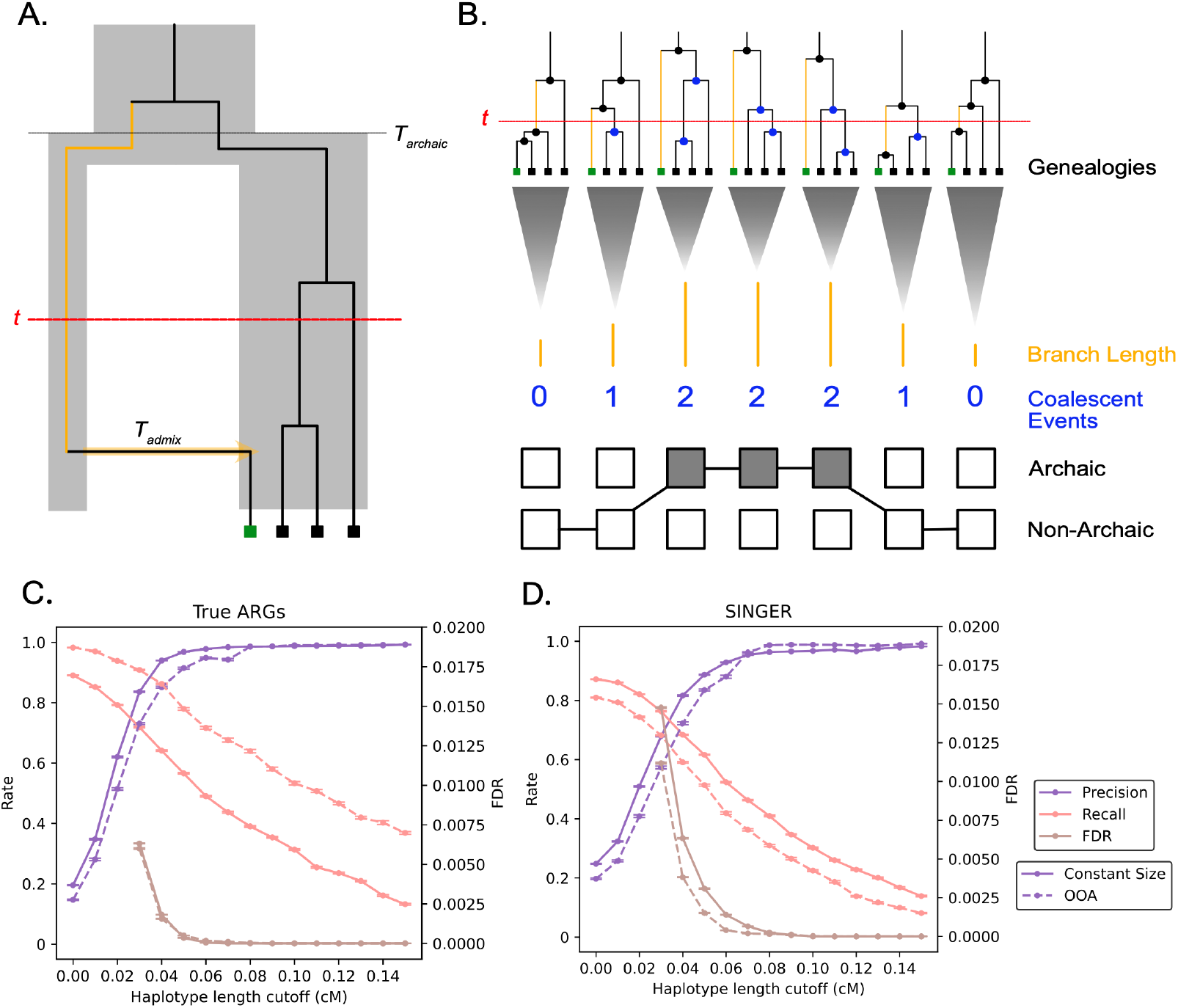
Conceptual overview and validation of TRACE. (A) Schematic of local genealogy relating four haplotypes under a demographic scenario of introgression, where introgression affecting the focal haplotype (green dot) creates long branches (orange branch) subtending that sample. Specifically, the long branch spans the timepoint of *t* generations, which is a user-defined timescale to identify archaic events inferred by TRACE, and is shown in the interval between the divergence time of the archaic and modern lineages (*T*_*archaic*_) and the time of the admixture event (*T*_*admix*_) in this figure. Individuals without archaic ancestry (black dots) coalesce at the standard rate within the ancestral population - generating a number of coalescent events during the timespan of the orange branch, which is modeled in the single-genealogy emission distribution. Note that time intervals are not drawn to scale. (B) Illustration of how the tree sequences are related to emission distributions of the focal branch-lengths and the number of non-focal coalescent events. Coalescent events that are incorporated into the emission distribution for each marginal tree are shown in blue (which occur during the span of the orange branch overlapping *t*). For simulations of a constant size model (solid line) and an Out-of-Africa (OOA) model (dashed line), we show the Precision (purple), Recall (pink) and False Discovery Rate (FDR, brown) when applying different haplotype length cutoffs (x-axis) in TRACE and using (C) ground-truth ARGs and (D) SINGER-inferred ARGs. Precision and recall fall between 0 and 1, shown as the “Rate” on the left y-axis; FDR falls between 0 and 0.02, shown on the right y-axis. See also Supplementary Section S1-3.

TRACE is implemented as a hidden Markov model with two states––archaic and modern human. As input, it uses ARGs inferred from phased whole genome sequences of contemporary individuals. For each marginal tree and focal haplotype, we calculate the emission probability by examining (1) the branch length subtending the focal sample at time *t* and (2) the number of *other* coalescent events on the tree that fall within the same time window (**Fig. 1B, fig. S1**). The transition probability depends on the time and magnitude of gene flow. The transition probabilities and the parameters of the emission distribution (modeled as Gamma distributions) are learned using the Baum–Welch algorithm (*45*). The parameter *t*—which defines the time cutoff for identifying “long” branches—is specified by the user and determines the timescale of introgression events targeted by the analysis. Finally, TRACE computes the posterior probability of archaic versus modern human ancestry at each genomic position using the forward-backward algorithm (*45*), integrating information across all trees to produce smoothed, position-specific state probabilities along the genome (**Supplementary Section S1**).

To validate TRACE, we simulated genome sequences for three populations–Africans, non-Africans and Neanderthals–under a demographic model incorporating an Out-of-Africa (OOA) bottleneck and ∼2% Neanderthal gene flow into non-Africans (*18, 46*). We applied TRACE to detect archaic ancestry in non-Africans using true ARGs for modern humans (**Supplementary Section S2, fig. S2**). To minimize the impact of population bottlenecks on the inference, we used joint ARGs for non-Africans (target, *n*= 100) and Africans (*n*= 100), since African populations do not share the OOA bottleneck with non-Africans. Applying *t* = 15,000 generations (close to the divergence time between Neanderthals and modern humans (*4*) and focusing on ancestry segments longer than 0.05 centi-Morgan (cM) to distinguish introgressed segments from ILS (*47*), we find TRACE has high (92%) accuracy and high (71%) recall. The false discovery rate is also low (FDR < 0.25%) as shown by applying TRACE to simulations with the same demographic history but no archaic introgression (**Fig. 1C, Supplementary Section S1.3**). Compared to published reference-free archaic ancestry inference methods such as *hmmix* (*29*) and *Sprime* (*30*), TRACE does not require an unadmixed outgroup for inference. In simulations, we observe that even small amounts of archaic gene flow into the outgroup can bias *hmmix* and *Sprime*, whereas TRACE maintains high sensitivity and specificity (**fig. S7**). TRACE shows robust performance across a range of additional demographic models and parameters, including constant population size models without the OOA bottleneck, varying proportions of Neanderthal ancestry (0.5–10%), varying times of divergence and gene flow, misspecification of user-specified parameter *t*, and target sample size (*n*= 10 − 200) (**fig. S3-S8, Supplementary Section S2.2-S2.3**).

Next, we applied TRACE to ARGs inferred using two recently published methods, SINGER (*36*) and *Relate* (*38*) (**fig. S9-S10**). Similar to the performance with true ARGs, TRACE achieves high precision (∼90%) and low FDR (<0.25%) using the inferred ARGs (**Fig. 1C-D, fig. S9**). However, the recall is substantially lower with inferred ARGs compared to true ARGs. For the OOA model (**fig. S2B**), TRACE recovers less than 10% of true segments using *Relate* and approximately 50% using SINGER, compared to ∼80% recall using true ARGs (**fig. S9**). Furthermore, recall decreases under more complex demographic models, though SINGER maintains higher recall (around 30%) than *Relate* under most demographic scenarios (**fig. S10**). Thus, we applied TRACE to SINGER-inferred ARGs for subsequent analysis of empirical data (**Supplementary Section S3, fig. S9-S14**).

### Evidence of ghost admixture in modern human populations before the Out-of-Africa migration

We analyzed 503 phased whole genome sequences from the 1000 Genomes Project (1000G), including 91 British (GBR) from Europe, 103 Han Chinese (CHB) from East Asia, 102 Indian Telugu (ITU) from South Asia, 108 Yoruba (YRI) from West Africa, and 99 Luhya (LWK) from East Africa (*48*). We applied SINGER (using only contemporary samples, no archaic genomes were included) using a mutation rate of 1.25 × 10^-8^ per base-pair (bp) per generation (*49*) and an effective population size of 20,000. We used 50 posterior tree sequences to account for uncertainty in ARG inference. To identify archaic introgression signals, we applied TRACE on SINGER-inferred ARGs with *t* = 15,000 generations (or 420,000 years, assuming a generation time of 28 years (*50*)), close to the estimated divergence time between modern humans and Neanderthals (*4*). To minimize the impact of ILS, we retained archaic segments longer than 50 kilo-basepairs (kbp) and 0.05 cM (which translates to FDR < 0.1% in simulations, **Fig. 1D**). We then compared variants on the inferred archaic segments with four high coverage sequenced archaic genomes (three Neanderthals (*2, 3, 5*) and one Denisovan (*1*)) to identify the source of archaic ancestry. For comparison, we also applied *IBDmix* (*28*) and *hmmix* to the same dataset and filtered with the same haplotype length filters. Unlike TRACE, both methods require either an archaic reference genome or an unadmixed outgroup: *IBDmix* was run using Neanderthal and Denisovan genomes, whereas *hmmix* used a panel of 426 sub-Saharan Africans as an outgroup. (**Supplementary Section S4**)

In non-African populations, TRACE detects an average of 0.99%, 0.97%, and 0.78% Neanderthal ancestry per individual in Europeans, East Asians, and South Asians, respectively (**Fig. 2A**). We recovered minimal Denisovan ancestry in Europeans (0.03%), with higher levels in East and South Asians (0.10% each), consistent with published results (*30, 51–54*). Overall, more than 90% of Neanderthal and 70% of Denisovan segments identified by TRACE overlap with segments inferred by *hmmix* or *IBDmix* (**table S4**). In both West and East Africans, TRACE detects less than 0.1% combined Neanderthal and Denisovan ancestry per individual. Around half of the Neanderthal segments identified by TRACE were also inferred by *IBDmix* (**table S4**, *hmmix* was not applied as there is no reliable outgroup for sub-Saharan Africans). We note that TRACE generally recovers less total Neanderthal and Denisovan ancestry than some previous estimates (*28, 53, 54*) (**Supplementary Section S4, fig. S15**), likely reflecting lower recall in inferred ARGs as seen in simulations (**Supplementary Section S5, table S7, fig. S10**). Nevertheless, TRACE robustly reconstructs the major features of Neanderthal and Denisovan introgression across global populations.

**Fig 2.**
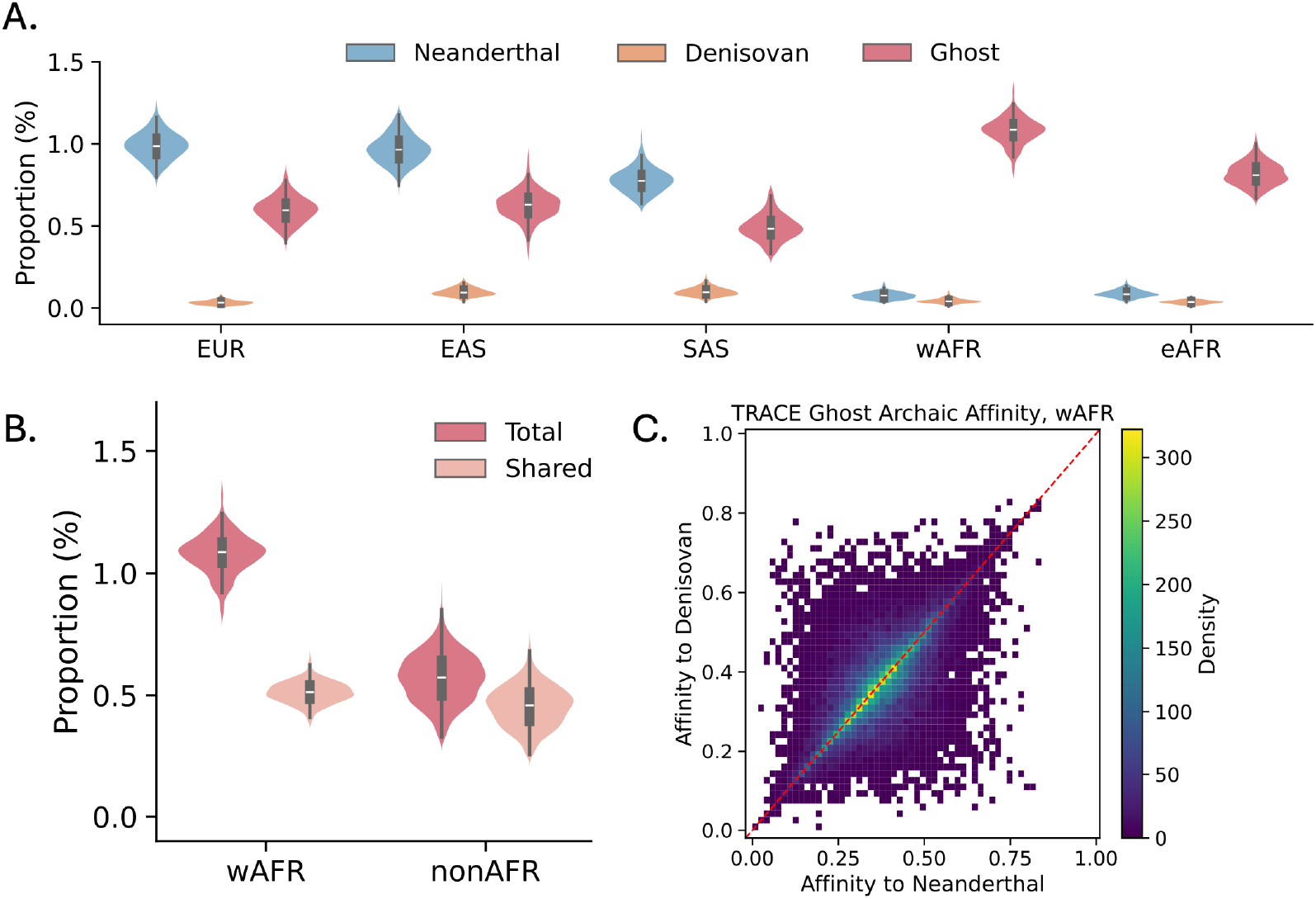
Archaic ancestry in global populations. **(A)** Proportion of archaic ancestry per genome recovered by TRACE using *t* = 15,000 generations that are > 50 kbp and > 0.05 cM. Archaic segments are classified as Neanderthal (blue), Denisovan (orange) and Ghost (red) based on allele sharing with sequenced Neanderthal and Denisovan genomes (Supplementary Section S4.2). **(B)** The amount of total (dark) and shared (light) ghost archaic ancestry per genome between West Africans and non-Africans. **(C)** Ghost segments affinity to Neanderthal (x-axis) and Denisovan (y-axis) genomes in West Africans, calculated as the proportion of derived mutations on the segment shared with Neanderthal and Denisovan sequences.

Beyond Neanderthal and Denisovan ancestry, TRACE revealed substantial “ghost” archaic ancestry––introgression from an archaic lineage more distantly related to Neanderthals and Denisovans––in all modern human populations studied. We detected 0.49–1.1% of ghost ancestry on average across populations (**Fig. 2A**). Most ghost ancestry segments found in non-Africans are shared with sub-Saharan Africans, while both East and West Africans harbor a greater diversity of unique ghost segments consistent with the reduction in genetic diversity in non-Africans caused by the OOA bottleneck (**Fig. 2B, fig. S22-S23**). Ghost segments exhibit deep divergence in marginal trees and show nearly identical genetic affinity to both sequenced Neanderthal and Denisovan genomes, indicating that they originated from an unsequenced lineage equally related to both archaic groups (**Fig. 2C, fig. S16-S18**). The average coalescence time between the ghost and modern human segments, inferred from introgressed segments in the inferred ARGs, is approximately 0.83 Mya (95% CI: 0.61–1.27 Mya, **table S6**).

Several additional lines of evidence support our results of ghost ancestry in modern humans. First, we find that genomic regions harboring ghost ancestry exhibit elevated heterozygosity levels, a pattern also observed for Neanderthal and Denisovan segments. The elevation in heterozygosity is consistent with a model of introgression from a deeply divergent lineage (*55*) and importantly, suggests that the ghost ancestry signal is not an artifact of ARG inference (**fig. S24**). Second, across all tested populations, ghost ancestry segments are shorter than Neanderthal and Denisovan segments, reflecting a more ancient introgression event predating Neanderthal gene flow (**fig. S25**). Additionally, we validated in simulations that in a model lacking ghost introgression, TRACE infers negligible levels of ghost ancestry (0.07%, 95% CI: 0–0.3%), significantly below the estimates from empirical data. In contrast, simulations including a ghost lineage produced results that closely matched empirical observations (**table S7-S8, fig. S27**). Furthermore, we computed the site frequency spectrum (SFS) and conditional SFS (cSFS) for variants present on archaic segments identified by TRACE (**fig. S19-S21, Supplementary Section S4**). While Neanderthal and Denisovan segments show the expected “U-shape” (*13*), ghost segments display distinct SFS and cSFS patterns that are consistent with a model of pre-OOA ghost introgression in simulations (**Supplementary Section S5, fig. S28-S29**).

Together, these findings suggest that an unknown archaic population, which diverged over 500,000 years ago, introgressed into the common ancestors of all modern humans prior to the OOA migration, resulting in similar patterns of ghost ancestry in non-Africans and Africans.

### The landscape of ghost ancestry in modern humans

To explore the legacy of ghost ancestry in modern humans, we examined the genome-wide distribution of ghost ancestry in global populations. Across all ghost haplotypes identified by TRACE in all populations (sub-Saharan Africans and non-Africans), we recovered 1548.75 Mbp or 71.54% of the accessible genome (**Fig. 3, Supplemental Section S6**). Broadly, the genome-wide patterns of ghost ancestry segments resemble those expected from hybridization between deeply divergent lineages (*56–58*). We find that ghost ancestry decreases in proximity to functional elements (lower *B* score (*59*); *ρ*_*Spearman*_ = 0.45, *p* < 2.2 × 10^−308^) and in regions of low recombination (*ρ*_*Spearman*_ = 0.34, *p* < 2.2 × 10^−308^), reflecting stronger effects of linked selection in these regions (*60, 61*). These patterns mimic those seen in Neanderthal and Denisovan introgression maps (*32, 52*) as well as patterns of hybridization across non-human species (*58*) (**fig. S30-S31, table S9, Supplementary Section S6.1**).

**Fig 3.**
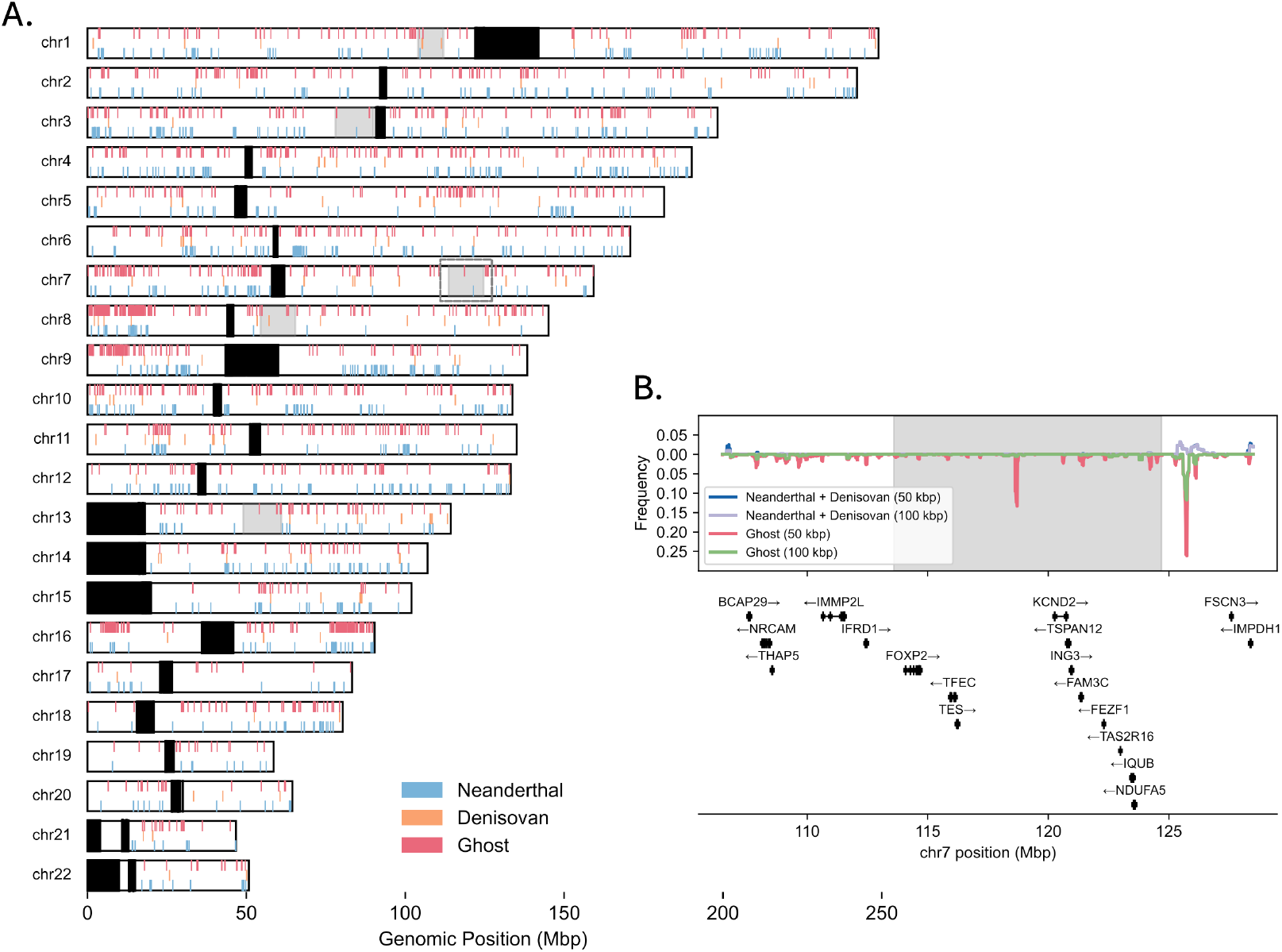
Characterizing the distribution of Archaic ancestry using TRACE. **(A)** Estimated peaks of archaic ancestry from TRACE—including ghost ancestry—aggregated across all populations. Black boxes indicate genomic regions masked as centromeric or genomic regions with no called variants (excluded from ARG inference). Previously estimated deserts of archaic (Neanderthal and Denisovan) ancestry are shown as gray highlights. **(B)** Frequency of Neanderthal, Denisovan and Ghost ancestry within the chromosome 7 archaic desert (shown as dashed grey box in panel A). The y-axis in the top panel reflects the frequency of Neanderthal and Denisovan ancestry, while the y-axis in the bottom panel reflects the frequency of Ghost ancestry within the region.

We also scanned for “peaks” of ghost ancestry, defined as regions with a significantly higher frequency of ghost ancestry compared to the genome-wide average estimate. Across all tested populations, we identified 1932 peaks of ghost ancestry (average length 61 kbp, SD 64 kbp) (**Supplementary Section S6.2, Fig. 3, table S12**). This result contrasts with the smaller number of 1155 and 160 peaks of Neanderthal and Denisovan ancestry, respectively (average length 84.97 and 76.54 kbp). We find that ghost ancestry peaks are over-represented in sub-Saharan African populations relative to non-African populations, which is consistent with the lower genetic diversity in non-Africans (**fig. S32-S33**). Several genes intersect with high-frequency peaks of ghost ancestry, such as *CSMD1* and *RBFOX1*, with an overall functional enrichment for immune and metabolic complexes, including the major histocompatibility and lipoprotein complexes (*p* < 10^−1^; Binomial Test; **fig. S34-S35**). We also identify 97 deserts of ghost ancestry (i.e., regions that are at least 10 Mbp long and have less than 0.1% frequency of ghost ancestry). Interestingly, all the deserts are only found in non-African populations, with nearly half (43.3%) shared between different non-African groups, likely formed during the OOA bottleneck (**Supplementary Section S6.3, fig. S36-S37**). We do not identify any ghost ancestry deserts in sub-Saharan Africans (though this may depend on the specific frequency and haplotype length thresholds applied). Overall, we find significant heterogeneity in the distribution of ghost ancestry tracts across the genome.

We also evaluated the presence of ghost ancestry in previously published archaic deserts, which are often considered as candidates of *Homo sapiens*-specific regions (*62, 63*). We examined five regions identified as shared Neanderthal and Denisovan deserts in two previous studies, three of which were suggested by both studies (**fig. S38-S44, table S10-S11, Supplementary Section S6.4**, (*52, 63*)). We replicate the absence of both ancestries in two deserts, but find non-negligible Neanderthal or Denisovan ancestry in other regions, as seen in recent studies (*53, 64*). Surprisingly, however, all five examined regions contain a substantial amount of ghost ancestry, even after restricting to longer and more confidently inferred segments (>100 kbp; **Fig. 3B, fig. S38-S39, table S10)**. Notably, the shared archaic desert on chromosome 7 containing the *FOXP2* gene has a peak of ghost ancestry with 13.3% frequency (**Fig. 3B**), and the Neanderthal desert on chromosome 13 contains a peak of ∼20% frequency overlapping *CSNK2A2IP* gene (**fig. S6.10**). Our findings suggest that ghost ancestry is often present in previously defined archaic deserts, providing additional context to putative *Homo sapiens*-specific selection.

### Detecting super-archaic ancestry in Oceanians

Previous studies examining patterns of allele sharing between Neanderthals, Denisovans, and modern humans have suggested Denisovans may harbor ancestry from a super-archaic population (*1, 2, 27*). Among modern humans, Oceanians and Southeast Asians derive the highest proportion of Denisovan ancestry (*51*). We therefore reasoned that a fraction of super-archaic ancestry may have been inherited through Denisovan gene flow and may persist within Denisovan introgressed segments in present-day Oceanians. To test this, we analyzed 92 high coverage whole genome sequences from Oceanian individuals including 25 individuals from Papuan New Guinea (*65*) and 67 individuals from Vanuatu and Santa Cruz Islands (*51*). We reconstructed the ARGs using SINGER along with 1000G YRI individuals and applied TRACE with *t* = 15,000 generations to detect archaic ancestry. For comparison, we also applied *IBDmix* and *hmmix*, using consistent parameters and filters as before (**Supplementary Section S7.1**).

We identified an average of 0.73% Neanderthal, 0.66% Denisovan and 0.33% ghost ancestry in Oceanian individuals. We note that the amount of archaic ancestry inferred by TRACE in Oceanian individuals is lower than previous studies (*51*), as well as other non-Africans in 1000G (**fig. S45**). In simulations, mimicking the demographic history of Oceanians including Neanderthal and Denisovan ancestry, we replicate similar underestimation of archaic ancestry (**Supplementary Section S8, table S14-S15**). As Neanderthals and Denisovans coalesce with each other (*t* = 13,600 − 16,900 generations, (*2*)) before they coalesce with modern humans, many of the long branches characteristic of the introgression are not detected, when using *t* = 15,000 generations as the time cutoff. The higher archaic introgression proportion further exacerbates this effect, as a larger fraction of coalescent events occur among archaic lineages (**Supplementary Section S7.2, table S14, fig. S46, S57**). Despite the lower recall, over 90% of the Neanderthal and Denisovan segments detected by TRACE overlap with segments detected by *hmmix* or *IBDmix* (**table S13**). Indeed, TRACE detected archaic segments showed archaic affinity patterns (**fig. S47**) and site frequency spectra (SFS/cSFS, **fig. S48**) that are consistent with those observed in 1000G data. Furthermore, ghost segments in Oceanians overlapped with those detected in 1000G populations (**fig. S49**), underscoring the shared nature of this ancestry.

To identify super-archaic introgression, we screened for “super-deep” coalescent events within Denisovan introgressed segments with *t* = 31,500 generations (similar to the inferred divergence time of the super-archaic lineage from modern humans (*27*)). To differentiate between introgressed segments and ILS, we focused on regions of at least 10 kbp and 0.01 cM and applied the same analysis to Neanderthal introgressed segments for comparison. Interestingly, we find a significantly higher proportion of these super-deep lineages in Denisovan introgressed segments than in Neanderthal segments (*p* < 2.2 × 10^−308^, Binomial Test; **Fig. 4, Supplementary Section S7.3**).

**Fig 4.**
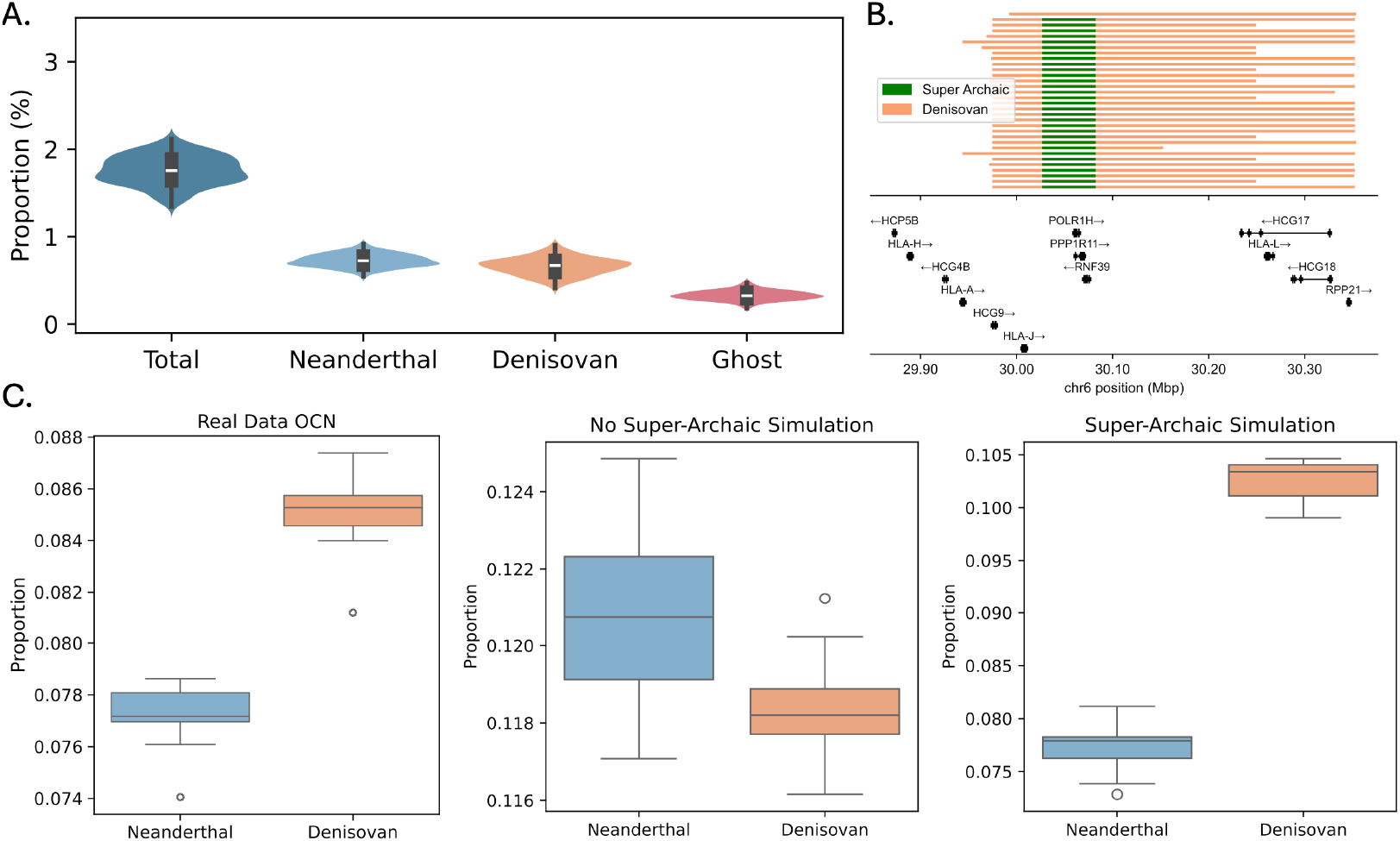
Super-archaic ancestry detected using TRACE within Oceanian (OCN) genomes. **(A)** Total proportion of Neanderthal, Denisovan, and Ghost ancestries across all sampled Oceanian individuals **(B)** Detected super-archaic ancestry in the MHC region lying on the background of Denisovan ancestry. **(C)** Denisovan segments have a significantly higher proportion of super deep lineages than Neanderthal segments, a signal reflective of super-archaic ancestry in simulations (*p* < 2.2 × 10^−308^, *p* = 1.00, *p* < 10^−36^ for real data OCN, No Super-Archaic simulation and Super-Archaic simulation respectively; Binomial Test).

We demonstrate in simulations that this pattern is not consistent with a model lacking super-archaic introgression, but is effectively recapitulated by a model including super-archaic introgression into Denisovans (**Fig. 4C, fig. S54-S56, table S14-S15**). Furthermore, the super-archaic fragments found within introgressed Denisovan segments exhibit distinct genetic features that differentiate them from the background Denisovan ancestry, including much deeper coalescence times with modern humans compared to other Denisovan introgressed segments, longer genomic lengths than mean ILS length inferred from deep lineages in Neanderthal segments, and low affinity to both Neanderthal and Denisovan reference genomes (assuming no super-archaic ancestry at the same locus in sequenced Denisovan, **Supplementary Section S8.4, fig. S58, table S16**). Using classification criteria developed from these features, we detected super-archaic segments in simulations with 70.2% accuracy (95% CI: 66.4–74.1%, **Supplementary Section S8.5**).

Applying this method to the Oceanian genomes, we infer that the super-archaic segments embedded within Denisovan ancestry tracts contributed almost 0.3% of the total detected Denisovan ancestry in Oceanians (**table S17**). This estimate constitutes a very conservative lower bound on the true fraction of super-archaic ancestry, as our analysis is restricted to Denisovan-introgressed regions in modern humans, requires segments longer than 20 kbp, and excludes loci where the sequenced Denisovan carries super-archaic ancestry (**fig. S58-S59, Supplementary Section S7.4**). Using the marginal trees in SINGER, we estimate the coalescence time between these super-archaic segments and modern human lineages to be approximately 1.77 Mya (95% CI: 1.13–3.98 Mya), consistent with earlier reports (*2, 27*). On average, super-archaic ancestry segments are 37.6 kbp, ranging between 20–83 kbp (**fig. S50**).

We evaluated the potential functional effects of super-archaic segments recovered from Oceanian genomes by characterizing common gene annotations and patterns of functional enrichment (**Fig. 4B**). Several genic regions harbor high proportions (>70%) of super-archaic tracts such as *RNF39, PPP1R11*, and *POLR1H* within the MHC (**Fig. 4B, fig. S51**) and *CYP24A1* that is a part of the cytochrome P450 family and a critical regulator of vitamin D degradation in humans (**fig. S52**, (*66, 67*)). Gene ontology analysis reveals significant enrichment for several pathways including MHC and Activity-related cytoskeleton (ARC), after multiple hypothesis testing (**fig. S53**). Overall, our results provide novel insights into the evolutionary history and potential functional legacy of super-archaic ancestry in modern humans.

## Discussion

In this study, we introduce TRACE, a method to identify archaic ancestry in modern human genomes using features of inferred ARGs. TRACE demonstrates robust performance across a range of demographic scenarios with both ground truth and inferred ARGs. Applied to data from the 1000 Genomes Project and Oceanian individuals, TRACE reveals two distinct and previously unresolved introgression events: ghost admixture into the ancestors of all modern humans prior to the Out-of-Africa expansion, and contribution from a super-archaic lineage that entered modern populations through Denisovan introgression. This work establishes a scalable, reference-free approach for mapping archaic ancestry, revealing new episodes of our complex evolutionary past.

TRACE also illuminates the enduring genomic legacy of archaic events, revealing lineage-specific patterns of enrichment and depletion of ancestry across the modern human genome. Notably, TRACE detects ghost ancestry within genomic regions previously identified as deserts of Neanderthal and Denisovan introgression. These deserts have often been interpreted as regions intolerant to archaic introgression due to positive selection for *Homo sapiens*–specific variation or strong purifying selection against archaic introgressed alleles (*56–58*). The persistence of deeply diverged ghost ancestry in these regions suggests there was lineage-specific selection acting against Neanderthal and Denisovan introgression, instead of positive selection for modern-human variation. This pattern is consistent with differential genetic load or epistatic incompatibilities associated with Neanderthal and Denisovan lineages, rather than with divergence time alone. Together, these findings indicate that introgression deserts do not represent universally archaic-free regions, but rather reflect selective filtering that depends on the source and evolutionary history of introgressed ancestry.

Recent work has reported evidence of unknown archaic ancestry, predominantly in West African individuals (*13–26*). In contrast to models suggesting unique archaic introgression in West Africans, our analysis points to at least one pre-OOA introgression event affecting all modern human populations (including West Africans, East Africans and non-Africans), a scenario most aligned with the model of Durvasula and Sankararaman (*13*). More recent research (*19, 23*) has proposed the existence of deep ancestral structure in the ancestors of modern humans, involving a deep population split (>1.5 Mya) followed by later rejoining to modern humans through a single introgression event around 300 kya (*23*) or through multiple “merger” events (*19*). Given that our inferred ghost lineage diverged at a similar time as the Neanderthal-Modern Human split (**table S6**), it appears to represent a more recent event than the proposed deep population structure events.

The two archaic introgression events we identify raise intriguing questions about their hominin sources. Archaeological records document numerous archaic forms with unclear amounts, if any, of direct contribution to modern humans (*26, 66, 67*). The ghost lineage, with a divergence time similar to Neanderthals, could plausibly correspond to Middle Pleistocene *Homo* groups (*70–72*) or African *Homo heidelbergensis* populations (*73–75*) that directly admixed with modern human ancestors before the OOA dispersal. For the super-archaic lineage, one potential candidate— compatible with the split time of approximately 1.8 Mya—is *Homo erectus*, as suggested by earlier studies (*2, 27, 76*).

Several open questions remain. First, the number, timing, and geographic locations of the inferred ghost introgression events are still uncertain: while our results support at least one pre–OOA introgression event affecting all modern human populations, additional episodes of gene flow among structured African populations may have occurred but remain difficult to resolve with present data. Second, the origin and geographic extent of the super-archaic lineage remain unresolved, including whether this ancestry entered modern humans exclusively via Denisovans or also through direct admixture with ancestors of modern humans. We anticipate that future studies, including new Denisovan reference genomes, may help to further pinpoint the timing and origin of these super-archaic sequences (*77*). Finally, while we detect signatures of selection influencing the distribution of ghost ancestry in the genome, the functional impacts of introgressed archaic regions, including their roles in immunity and metabolism, have yet to be fully characterized. Future studies that integrate additional ancient genomes at deep timescales, particularly from Africa and Asia, will be critical for refining the timing, sources, and evolutionary consequences of these introgression events.

## Acknowledgments

We thank Rasmus Nielsen, Nick Patterson, David Reich, Tianyi Wang, Iswar Hariharan and members of the Moorjani and McCoy labs for helpful discussions. We thank Yun Deng for technical support with SINGER, Laurits Skov for helpful discussions related to *hmmix* and *IBDmix*, and Joshua Bauman and Graydon Moorhead for testing of TRACE. We thank Nick Patterson, Montgomery Slatkin, Elise Kerdoncuff, Sophie Joseph, Michael Tassia, Rajiv McCoy and Etienne Patin for comments on the manuscript.

## Author contributions

Conceptualization: Y.Z., A.B., C.O.D., P.M.; Data curation: Y.Z., S.A.J., A.B., C.O.D., P.M.; Formal analysis: Y.Z., A.B.; Funding acquisition: P.M.; Methodology: Y.Z., A.B., P.M.; Software: Y.Z., A.B.; Supervision: C.O.D., P.M.; Validation: Y.Z., P.M.; Writing– original draft: Y.Z., A.B., P.M.; Writing– review & editing: Y.Z., A.B., S.A.J, C.O.D, and P.M

## Data and materials availability

All resources used in this study are listed and available in table S1-S2. TRACE is available on github: https://github.com/YulinZhang9806/trace

## Supplementary Materials

Supplementary Text

Figs. S1 to S59

Tables S1, S2, S4-S11, S13-S17

References(*78–96*)

## Other Supplementary Materials for this manuscript include the following

Tables S3 and S12

