## supplementary_text for "Recovering signatures of archaic introgression using ancestral recombination graphs"

**The PDF file includes:**

Supplementary Text
Figs. S1 to S59
Tables S1, S2, S4-S11, S13-S17
References (*78–96*)

**Other Supplementary Materials for this manuscript include the following:**

Tables S3 and S12

### Supplementary Text

### Section S1. TRACE: Method Details

#### S1.1 HMM Definitions

TRACE is implemented as a two-state hidden Markov model, with emission and transition densities that capture features of sequential genealogies along a chromosome. The emission distribution reflects the expectation that introgression at a focal haplotype from a sufficiently diverged population results in "long branches" in the marginal trees subtending the lineage (**Fig. 1A**). The transition probability distribution models the persistence of long branches across multiple adjacent trees, indicative of "long haplotypes" introduced through introgression (**Fig. 1B, fig. S1**).

For each marginal tree and focal haplotype of interest, we define the focal branch as the branch that subtends the focal sample at $t$ generations. We then model the product of 1) the branch length ($L^{(t)}$) of the focal branch, and 2) the number of coalescent events ($N^{(t)}$) occurring elsewhere on the tree during the time interval spanning the focal branch (**Fig. 1B, fig. S1**). Let $Z_i$ denotes the hidden state (0–modern human, 1–archaic), to capture these features jointly, we model the product of these two random variables — $X^{(t)} = L^{(t)} \times N^{(t)}$— as a Gamma distribution at the $i^{th}$ local tree

$$P(X_i^{(t)} | Z_i, \alpha_0, \beta_0, \alpha_1, \beta_1) = \begin{cases} Gamma(\alpha_1,\beta_1), Z_i=1 \\ Gamma(\alpha_0,\beta_0), Z_i=0 \end{cases}$$

where the parameters $(\alpha_0, \beta_0)$ describe the distribution of $X^{(t)}$ for haplotypes without introgression (modern human variation) and $\alpha_1, \beta_1$ represents haplotypes with introgression.
For parameter initialization, we consider the genome-wide distribution of $X^{(t)}$ across all marginal trees along the genome as an approximation to the null model of no introgression (i.e. modern human variation). To minimize bias due to any potential introgression signals, we exclude positive outliers of $X^{(t)}$ using the generalized Extreme Studentized Deviate (ESD) test (critical value $p = 0.05$). We use maximum likelihood to estimate the parameters $\widehat{\alpha_0}$ and $\widehat{\beta_0}$ of the gamma distribution of the null state. The outliers flagged by the ESD test are used to initialize the estimation of $\alpha_1, \beta_1$ for the alternate distribution, also via maximum likelihood estimation.

The transition model between the null and introgression state is:

$$R = \begin{bmatrix} 1-p & p \\ q & 1-q \end{bmatrix}$$

where $p$ is the transition probability from the modern human to the archaic state and $q$ denotes the reverse probability (archaic to modern human). To infer the parameters of the model $\hat{\Theta} = \{\widehat{\alpha_1}, \widehat{\beta_1}, \hat{p}, \hat{q}\}$, we used the Baum-Welch algorithm with a stopping criteria when the log-likelihood changes less than $10^{-2}$. In all cases, the parameter $t$ – which defines the time cutoff for identifying long branches – is set by the user and fixed throughout analysis.

Following parameter inference, we use the estimated maximum-likelihood parameters for calculating the posterior decoding for being in each state (archaic, modern human) across all marginal trees in the ARG using the forward-backward algorithm (*45*). We then extract all archaic ancestry segments with posterior probability > 0.9 and a length of at least 0.05 centimorgans (cM) and 50 kilo-basepairs (kbp) unless otherwise specified.

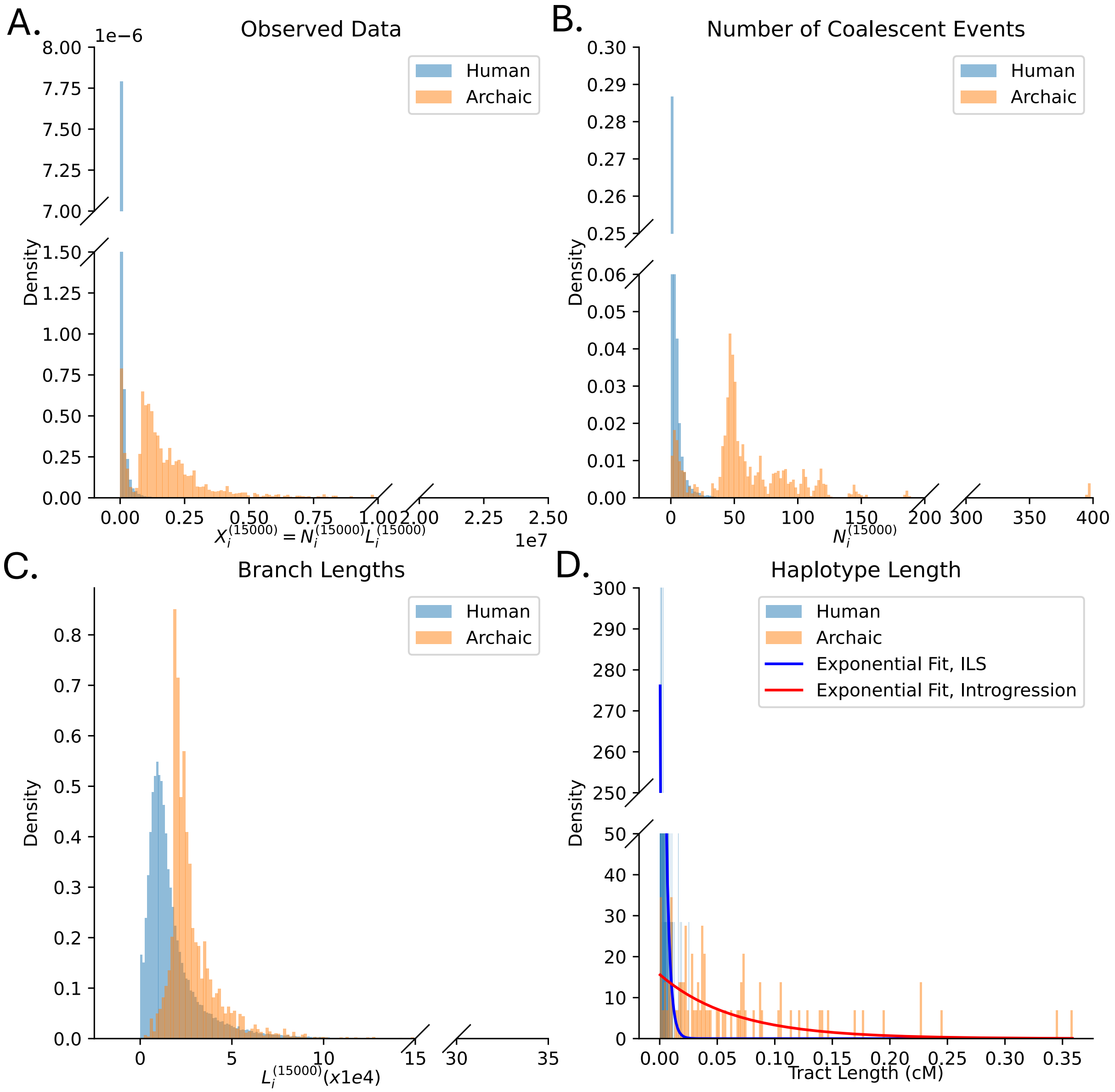

**Fig S1.** Summary statistics for TRACE observation data from a coalescent simulation under the Constant Size Model (**fig. S2A**) with 2% introgression proportion. All panels show results from a single simulation analyzing one target haplotype. For each marginal tree in the tree sequence, we extracted the branch subtending the target haplotype at $t = 15000$ generations. **(A)** Distributions of observation data ($X^{(t)} = L^{(t)} \times N^{(t)}$). **(B)** Distributions of the number of coalescent events ($N^{(t)}$) in the marginal tree during the branch's timespan. **(C)** Distributions of branch lengths ($L^{(t)}$), colored by true ancestry (human vs. archaic). **(D)** Length distribution of genomic segments with long branches, classified by their origin (Incomplete Lineage Sorting [ILS] vs. Introgression).

#### S1.2 Incorporating genomic accessibility masks

TRACE allows users to specify genomic accessibility masks to exclude low-confidence genomic regions. In any marginal tree with less than 99% overlap with the accessible genome, the emission probabilities are set to 0 for both "Archaic" and "Modern Human" states, effectively making the model blind to the specific tree topology and branch lengths in that region. However, the Markov chain remains continuous, so the posterior probabilities for a masked region are interpolated based on the neighboring trees via the transition probability distribution.

#### S1.3 Estimating the performance of TRACE

To evaluate the reliability of TRACE, we performed simulations where the true ancestry for a set of sequences is known. We apply TRACE to identify archaic ancestry segments and compare these to true archaic segments, and infer the precision and recall based on segment overlaps.

Let $L_{overlap}$ be the total length (in basepairs, bp) of the overlap between inferred and true segments, $L_{inferred}$ be the total length of inferred archaic segments, and $L_{truth}$ be the total length of true archaic segments. Precision and recall are calculated as:

$$Precision = \frac{L_{overlap}}{L_{inferred}}$$
$$Recall = \frac{L_{overlap}}{L_{truth}}$$

To estimate the False Discovery Rate (FDR), we applied TRACE to null simulations (same demographic model but no archaic introgression). The FDR is given by:

$$FDR = \frac{L_{FD}}{L_{genome}}$$

where $L_{FD}$ is the total length of segments falsely inferred as archaic in the null simulations, and $L_{genome}$ is the total length of the simulated genome.

### Section S2. Assessing Power and Robustness of TRACE in simulations with ground-truth ARGs

#### S2.1 Simulation and evaluation settings

The objective of TRACE is to identify archaic introgression—specifically, deep introgression events from highly divergent lineages. Although the framework can be applied to introgression detection at different timescales by adjusting the user-defined parameter *t*, our analyses focus on older events. To evaluate performance, we conducted simulations using *msprime* (*35*) mimicking the demographic history of Neanderthal introgression into non-Africans. We used a genome-wide recombination rate of $10^{-8}$ per bp per generation and a mutation rate of 1.2 x $10^{-8}$ per bp per generation (*78*). For each demographic model, we generated 10 replicate simulations of 50 Mbp. From each simulation under the Constant Size Model (**fig. S2A**), we sampled 100 diploid individuals (200 haplotypes) from the target population A. For simulations under the other extended models (**fig. S2B-D**), we sampled 100 diploid individuals from population A and 100 diploid individuals from population C (200 individuals, 400 haplotypes in total).

We considered a range of demographic models and parameters:

Constant Size (**fig. S2A**): We simulated data for three populations–Africans (Population C), non-Africans (Population A) and Neanderthals (Population B) with the following parameters):

- Populations A, B, and C had a constant effective population size ($Ne$) of 10,000 throughout their history.
- The archaic lineage B diverged from the modern human lineage C at 19,275 generations before present ($T_{archaic} = 19{,}275$).
- The modern human populations A and C diverged at 2,093 generations before present.
- A single instantaneous pulse of introgression from archaic population B into human population A occurred 1,724 generations before present ($T_{admix} = 1{,}724$) contributing 2% ancestry to population A.

We extended the Constant Size Model to incorporate more complex demographic histories, including an Out-of-Africa (OOA) model and two derivative scenarios where the archaic introgression precedes the OOA split and another with post-OOA gene flows between Africans and non-Africans (population C and A):

OOA (**fig. S2B**): Following (*18*), we modeled the OOA demographic scenario with the following parameters. The modern human population C started with $Ne = 20{,}000$. Around 20,345 generations ago, its population size expanded to $Ne = 25{,}900$. The archaic population B diverged from C around 19,275 generations ago with a constant $N_e$ = 3,600. Population A experienced a bottleneck immediately after splitting from population C at 2,093 generations ago, and its $N_e$ was 880 during the bottleneck that lasted for 852 generations. This was followed by an exponential growth phase starting 1,241 generations ago with a $Ne = 2{,}300$ and continuing to the present and reaching $Ne = 10{,}855$.

Archaic Introgression before OOA (**fig. S2C**): All demographic parameters are identical to the OOA model, except the introgression from population B to C occurred earlier at 2,500 generations before present.

Gene Flow between Modern Humans after OOA (**fig. S2D**): All demographic parameters are identical to the OOA model, except that populations C and A experienced continuous gene flow since the split of A from C: first at a rate of 0.000522 for a duration of 852 generations during the OOA bottleneck, then a rate of $2.48 \times 10^{-5}$ until present after the OOA bottleneck (*18*).

We applied TRACE with $t = 15{,}000$ generations to the true ARGs for population A (for the Constant Size Model, **Section S2.2-S2.3, S2.5**) or the joint ARGs of populations A and C (for the extended models, **Section S2.4**) and retained all segments which had a posterior probability threshold greater than 0.9. To evaluate performance, we measured precision, recall, and FDR (**Section S1.3**) using haplotype length cutoffs ranging from 0.01cM to 0.15cM. For each simulation, we measured performance statistics for 10 haplotypes (5 diploid individuals) from population A, resulting in 100 data points in total per tested scenario (10 replicates x 10 haplotypes). We assessed uncertainty by performing 30 bootstrap replicates, resampling 90% of the data without replacement each time.

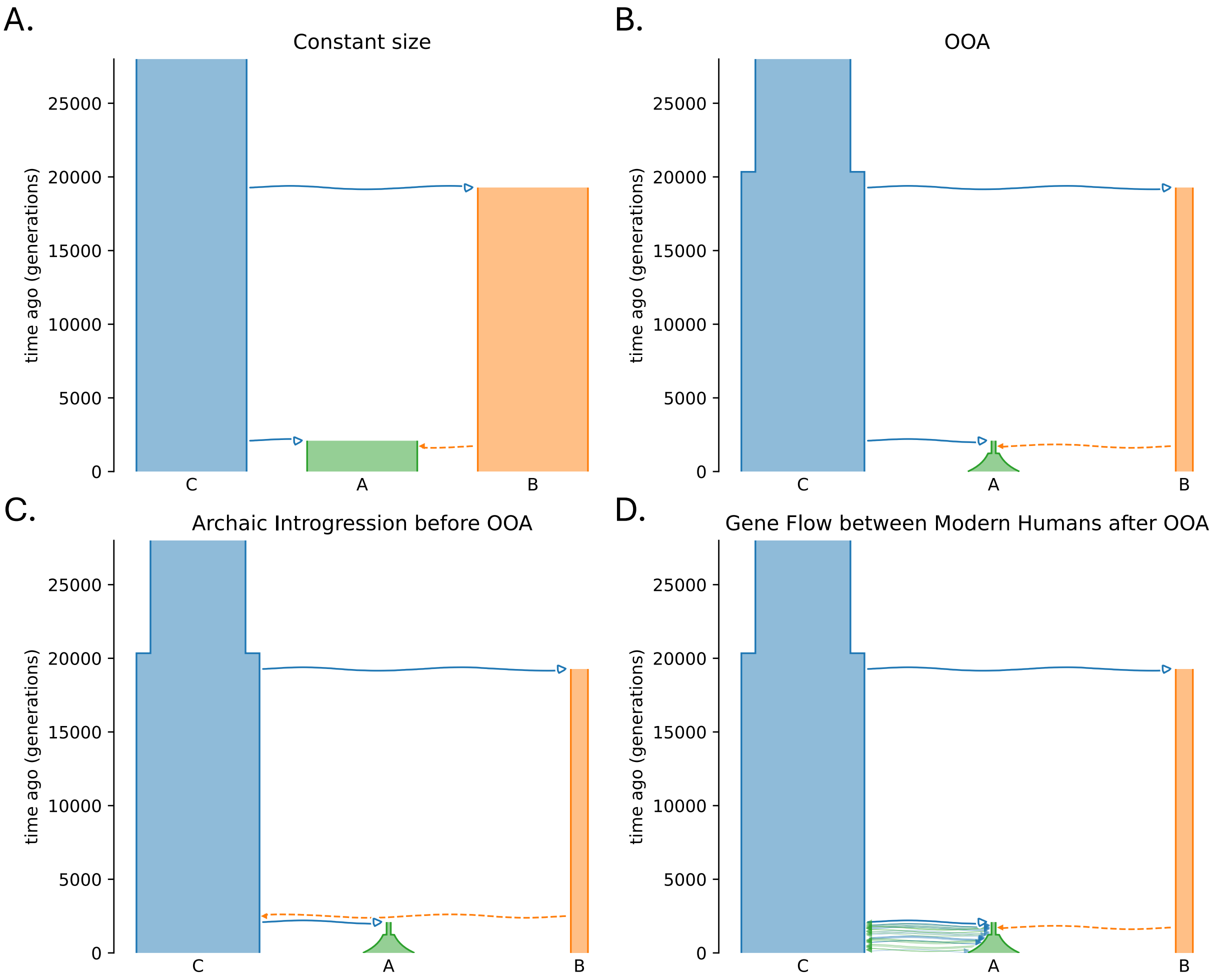

**Fig S2**. Demographic models for TRACE evaluation. **(A)** Constant Size: All populations (A: target modern human, B: archaic, C: outgroup modern human) maintain a constant size ($N_e = 10,000$). **(B)** Out-of-Africa (OOA): A simplified model based on Ragsdale et al. (2019) (*18*) with discrete divergence and a single pulse of archaic introgression. **(C)** Introgression before OOA: An OOA model incorporating an older pulse of archaic introgression into the ancestors of both modern human groups A and C prior to the bottleneck. **(D)** Gene Flow after OOA: The original (non-simplified) OOA model from (*18*)---with continuous gene flow between the target (A) and outgroup (C) populations after their divergence.

#### S2.2 Impact of input parameters

We tested the impact of key input parameters: (i) the user-defined time threshold $t$, (ii) posterior probability thresholds, (iii) the number of target samples included in the ARG, and (iv) proportion of introgression. We used the Constant Size Model (**fig. S2A**) with $t = 15000$ generations for all tests, except for those specifically analyzing the parameter $t$ itself.

Using the true ARGs, we find TRACE performs reliably to identify archaic introgression (**fig. S3-S4**). TRACE requires a key input parameter $t$, which is used to identify long branches in the marginal trees of the ARG. One should think of this parameter as "the time defining an archaic event": optimal performance is achieved when $t$ is close to the true divergence time $T_{archaic}$; if $t$ is set much older than $T_{archaic}$, TRACE would fail to capture a proportion of true signals where the introgressed archaic lineage coalesces with modern human lineages before $t$. Alternatively, if $t$ is set too young, archaic inference results may be confounded by other recent demographic events. In simulations ($T_{archaic} = 19{,}275$ generations), we find the precision of TRACE remains robust regardless of the input $t$, with precision > 80% for a range of values between $t = 10{,}000 - 25{,}000$ generations. (**fig. S3**). However, the recall is highest when $t < T_{archaic}$ (e.g., $t = 10{,}000$ or $15{,}000$ generations), because archaic segments that coalesce with modern human lineages before time $t$ are missed when $t > T_{archaic}$ (e.g., $t = 20{,}000$ or $25{,}000$ generations).

We find the posterior probability thresholds had a minimal impact on the performance of TRACE– applying posterior thresholds of 0.8, 0.9 and 0.95 yield almost identical results. In terms of introgressed haplotype length, the longer haplotypes are more likely to be true introgression signals, as expected (since shorter haplotypes can be due to incomplete lineage sorting (ILS) and introgression). In this simulation, the theoretical mean introgressed haplotype length is approximately 0.06cM, assuming for a single instantaneous 2% introgression event that occurred 1724 generations ago (*79*). Thus, using a threshold above 0.06cM effectively differentiates introgressed segments from ILS, and is exemplified by the precision of 98% and FDR < 0.05% (**fig. S4A**).

Considering the sample size (e.g. the number of diploid individuals included in the input ARG for TRACE, $n = 10 - 200$) of the target population A, we find TRACE delivers robust performance across a range of sample sizes as low as 10 diploid individuals (**fig. S4B**). Moreover, TRACE can reliably detect introgression signals at ancestry proportions as low as 0.5%, with precision improving at higher proportions. However, recall decreases substantially with increasing introgression proportion. This trade-off arises because increased introgression leads to more frequent coalescence events *between* archaic lineages prior to time $t$, thereby breaking long branches that serve as the primary signal of archaic introgression in TRACE (**fig. S4C**).

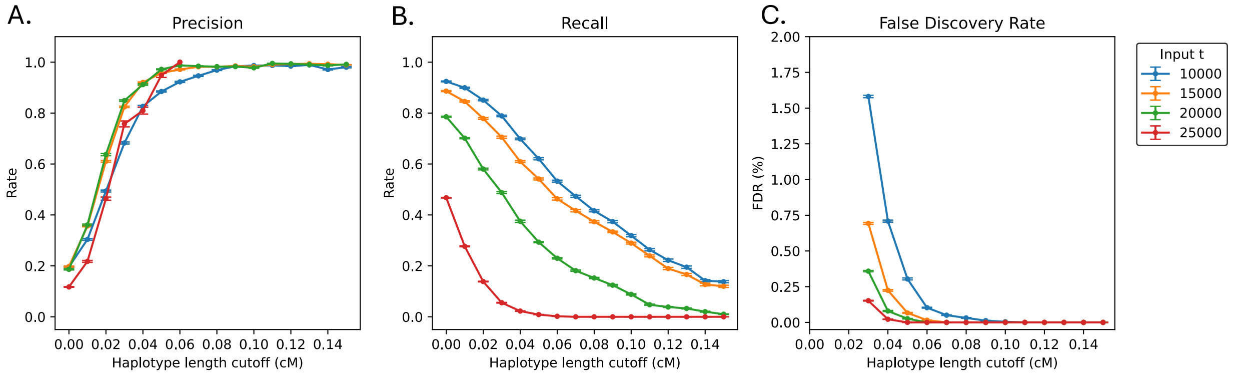

**Fig S3**. TRACE performance applying different user-defined $t$. Evaluations are done based on true ARGs (Constant Size Model, 2% introgression proportion, $T_{archaic} = 19{,}275$). **(A)** Precision, **(B)** Recall, and **(C)** False Discovery Rate (FDR) for segments (posterior probability > 0.9) across different haplotype length cutoffs. Colors indicate different input time parameters t. Error bars represent standard error from 30 bootstraps (90% of 100 total data points resampled).

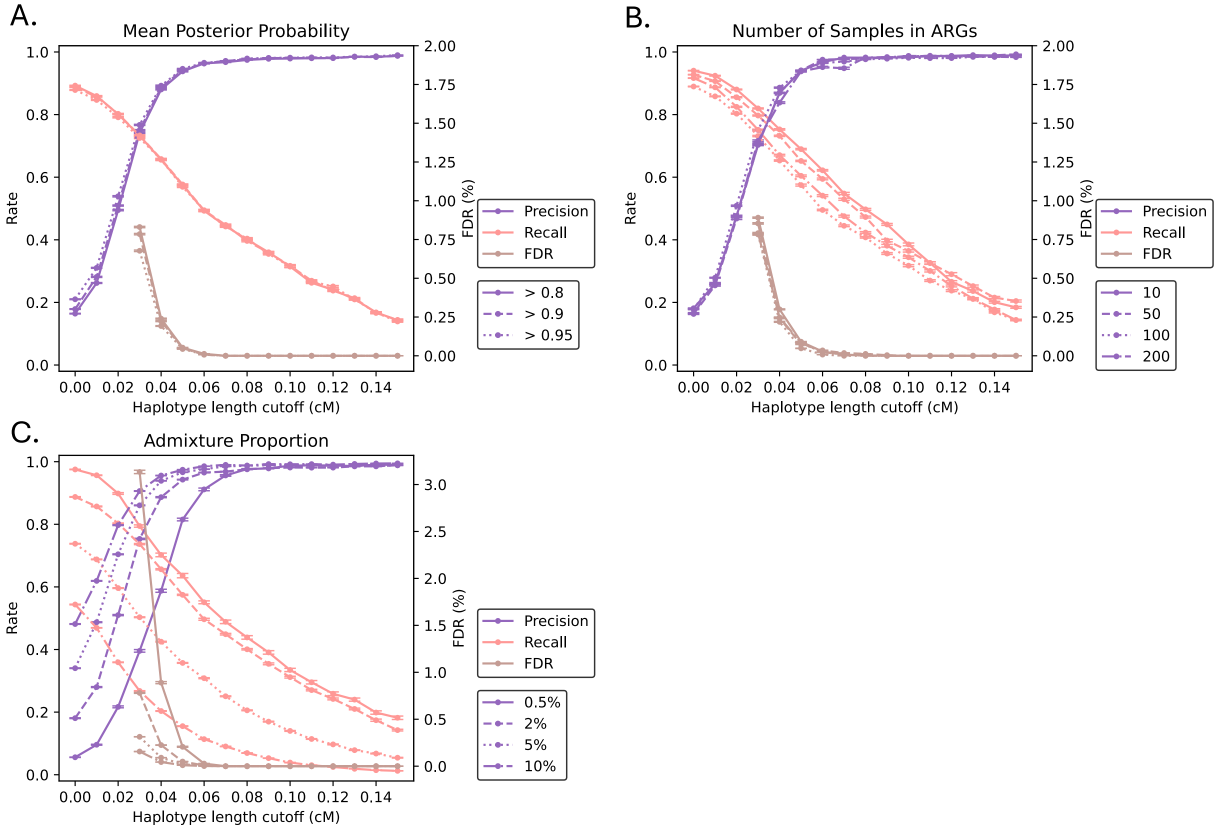

**Fig S4.** Evaluation of TRACE performance under the Constant Size Model, assessing sensitivity to: **(A)** Mean posterior probability threshold for segment calls, **(B)** Number of diploid samples included in ancestral recombination graphs (ARGs), and **(C)** Simulated introgression proportion. Performance metrics include precision (purple, left), recall (pink, left), and false discovery rate (brown, right), with error bars representing standard errors across 30 bootstrap replicates (90% of data each time; 100 data points total: 10 haplotypes per simulation × 10 replicates).

#### S2.3 Robustness of TRACE under extreme demographic scenarios

We evaluated the robustness of TRACE under various demographic scenarios.

First, we modified the Constant Size Model by varying the divergence time between populations B and C ($T_{archaic} = 9275, 14275, 19275, 21275, 24275$ generations) and the introgression time from B to A ($T_{admix} = 724, 1224, 1724, 3724, 5724$ generations). This allowed us to test scenarios where the introgressed lineages in marginal trees are deeper or shallower compared to the modern Human-Neanderthal population history. We find TRACE is robust to changes in $T_{archaic}$ (**fig. S5B**), with precision only being affected when $T_{archaic}$ is as recent as 9,275 generations. However, TRACE's recall decreases for older $T_{admix}$ if we use a fixed haplotype length cutoff (for example, > 0.05cM or 50kbp) (**fig. S5A**). As introgression events become older, recombination breaks down introgressed segments into progressively shorter tracts, making it increasingly difficult to differentiate them from ILS tracts (*79*).

Next, we tested the impact of population size changes in the target population A. We simulated two types of scenarios:

- Bottlenecks: Population A experienced a severe bottleneck of varying duration (850, 1050, 1250 generations) and intensity (during bottleneck $Ne = 480, 680, 880$) after its split from C, before introgression from B.
- Expansions: Population A undergoes a post-introgression expansion to 5x or 10x its original population size.

While population expansions have little effect on the inference of TRACE (**fig. S6C**), we find bottlenecks in the target population A can significantly reduce precision (**fig. S6AB**). This is because bottlenecks create long branches in ARGs due to reduced population diversity, making it difficult to distinguish true archaic introgression from the background variation. To mitigate the confounding effect of bottlenecks, we ran TRACE on the joint ARG of populations A and C (**Section S2.1**). Because population C has not experienced a severe bottleneck, its lineages provide a comparative background in which long branches arising from the bottleneck in population A are less pronounced, thereby improving the discrimination of archaic signals from modern human variation. It is important to note that population C is not used as an "outgroup" without archaic ancestry in TRACE as in other reference-free archaic inference methods like *hmmix* (*29*) and *Sprime* (*30*). This joint analysis effectively removes the impact of a simulated OOA bottleneck on the performance of TRACE, yielding comparable performance metrics under the Constant Size and OOA models (**fig. S7**).

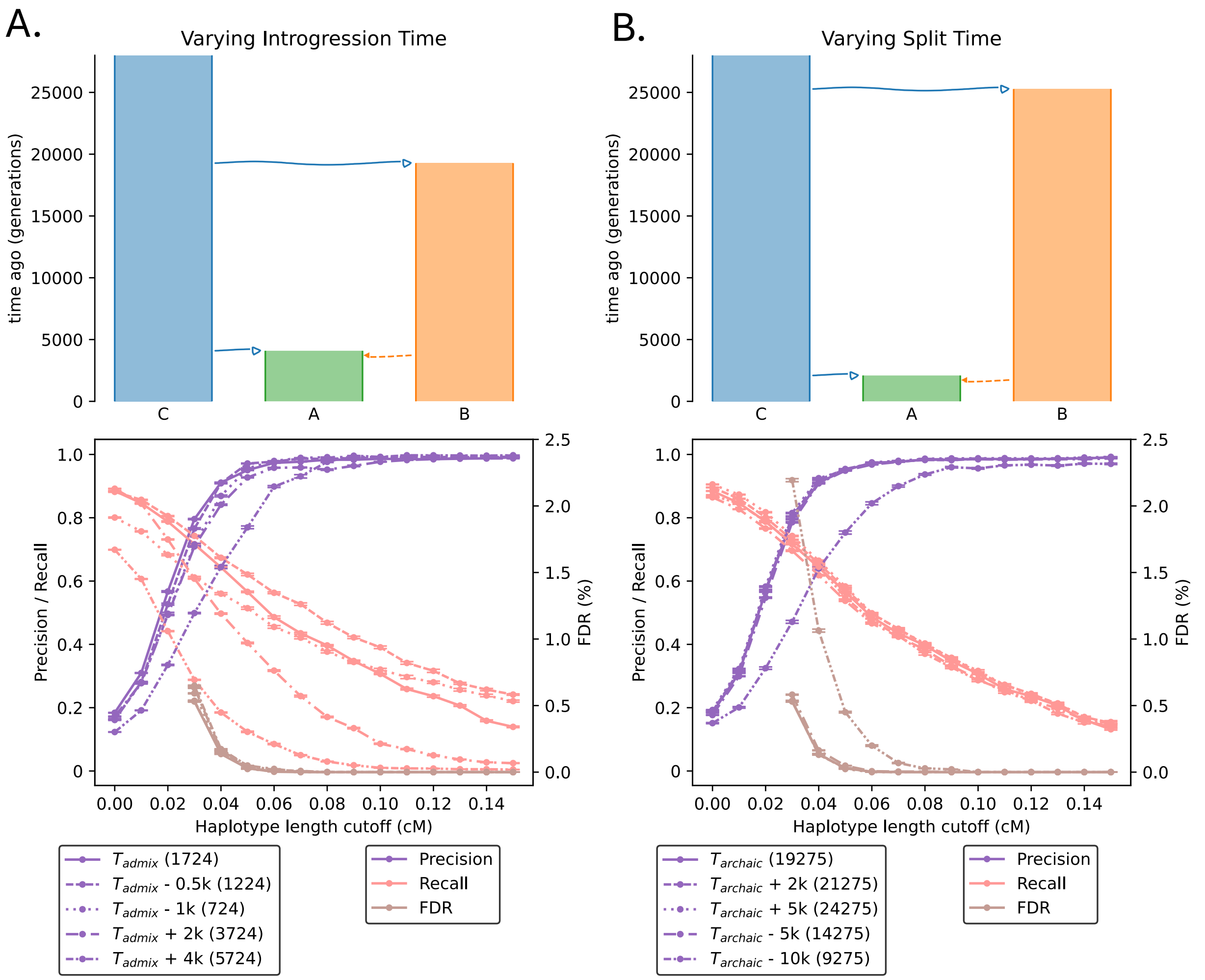

**Fig S5**. Sensitivity of TRACE to divergence and introgression times. Performance was evaluated under the Constant Size Model with input $t = 15000$ (except $t = 9275$ for the $T_{archaic} = 9275$ scenario in (B). **(A)** Varying introgression time from B to A. **(B)** Varying split time between B and C. Metrics show precision (purple, left), recall (pink, left), and FDR (brown, right). Error bars represent standard error from 30 bootstraps of 90% of 100 data points.

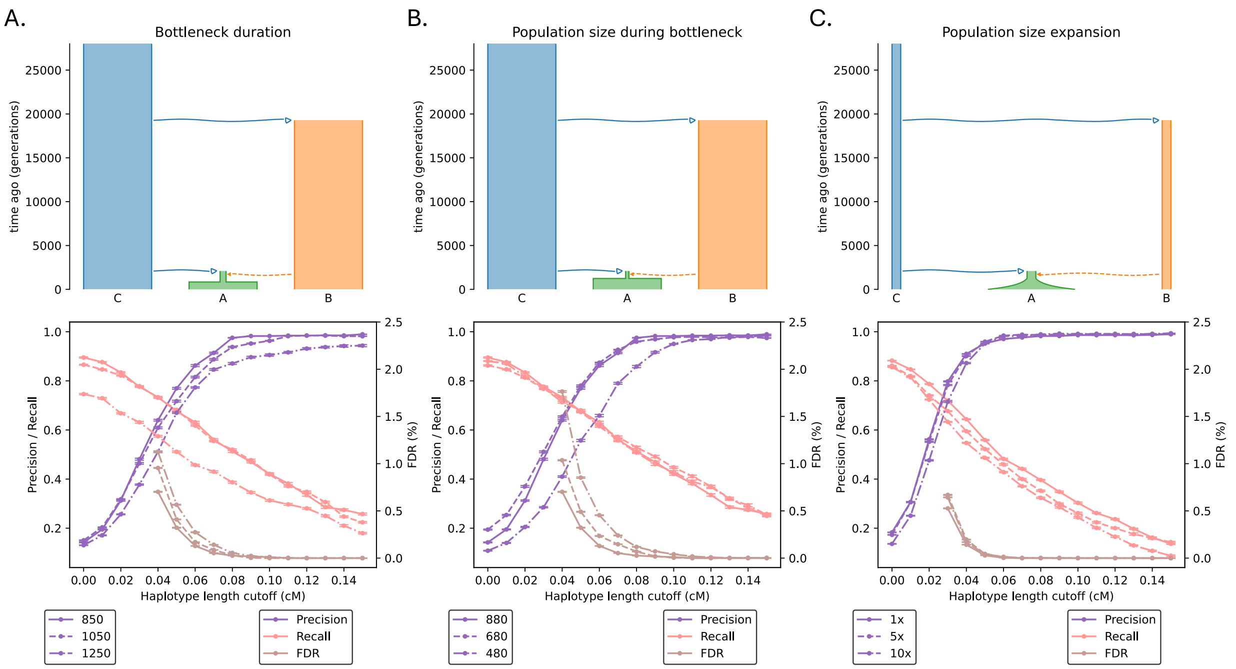

**Fig S6**. Sensitivity of TRACE to demographic changes in the target population. Performance was evaluated under the Constant Size Model for: **(A)** Bottleneck duration, **(B)** Population size during bottleneck, and **(C)** Population size expansion. Precision (purple, left), recall (pink, left), and FDR (brown, right) are shown, with error bars representing standard error across 30 bootstraps of 90% of the data.

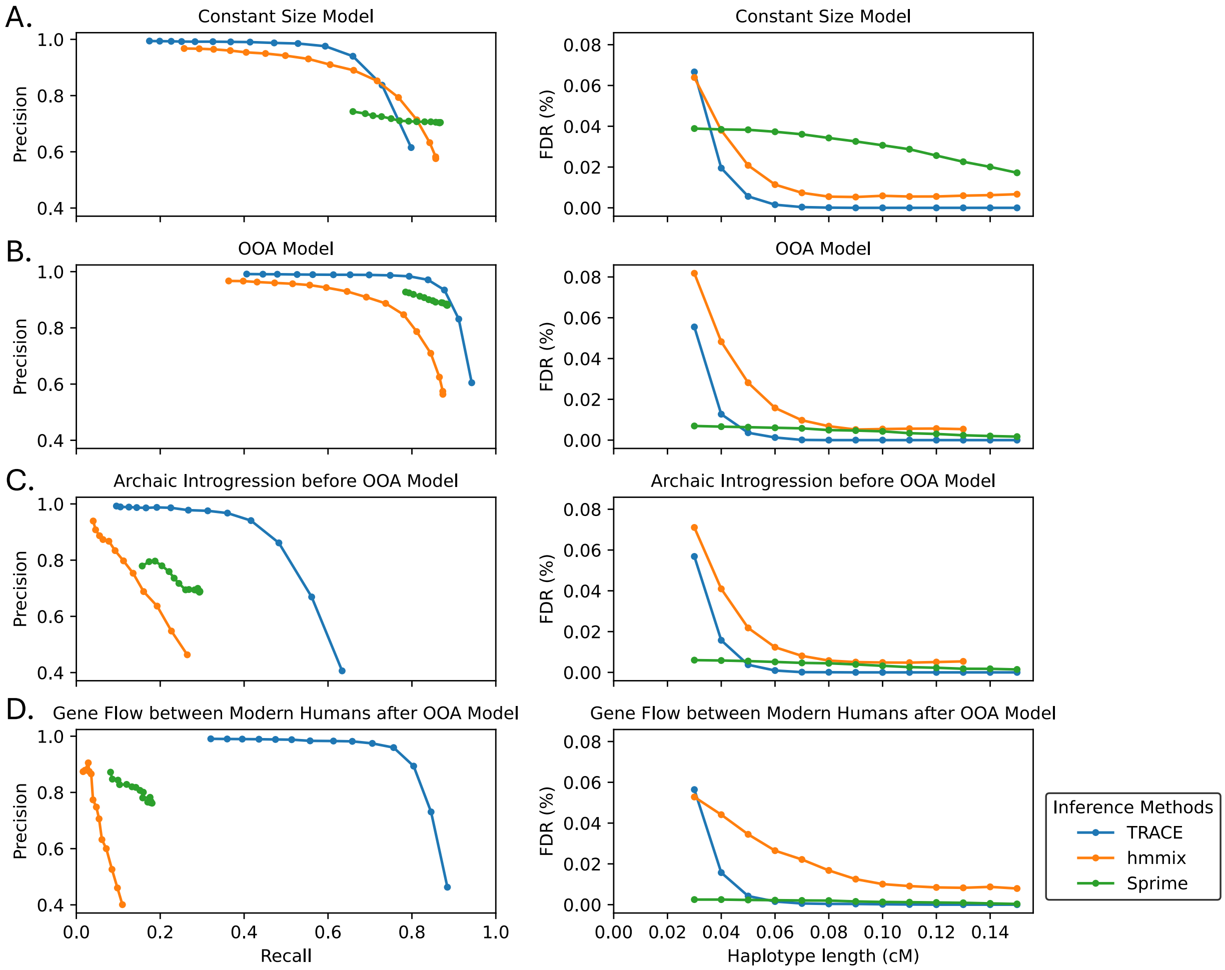

**Fig S7.** Performance of TRACE, *hmmix* and *Sprime*. Precision-recall curves (left) and false discovery rate, FDR (right), for TRACE (blue), hmmix (orange), and Sprime (green) across demographic models with 2% archaic introgression: **(A)** Constant Size, **(B)** Out-of-Africa (OOA), **(C)** Archaic Introgression before OOA, and **(D)** Gene Flow between Modern Humans after OOA.

#### S2.4 Comparing performance of TRACE with existing methods

To compare performance of TRACE with published reference-free archaic inference methods (**fig. S7**), we performed simulations under four demographic models (**fig. S2, Section S2.1**). A key methodological difference between TRACE and other reference-free archaic inference methods like *hmmix* and *Sprime* is that TRACE does not require an outgroup without archaic introgression.

To ensure robust performance evaluation with *hmmix*, which can be biased when trained on smaller genomic datasets, we generated 10 simulation replicates of 250 Mbp genomes for each demographic scenario. We sampled 100 individuals each from populations A (non-Africans) and C (Africans) (200 diploid genomes in total). TRACE was applied to the joint genealogies of sampled individuals from populations A and C and used $t = 15{,}000$ generations and a haplotype length threshold of 0.05cM. For *hmmix* and *Sprime,* we used 100 individuals from population C as the outgroup (without archaic introgression). Following user documentations, we retained *hmmix* results with a posterior probability > 0.8 and filtered *Sprime* results using a score threshold of 5 x $10^4$. Fragments from both methods were then filtered to haplotypes larger than 0.05cM.

We evaluated the performance of these three methods based on the metrics described in Section S2.1, with one adaptation for *Sprime*: As *Sprime* infers archaic segments at the population level rather than for individual level, we calculated population-level precision and recall for *Sprime*. This was done by merging the ground-truth segments from the 10 individuals that we used to evaluate method performance and comparing this merged set of true introgressed segments to inferred segments by *Sprime* for the same group of individuals. The FDR for *Sprime* was calculated by dividing the total length of false discoveries across 10 individuals by the sample size (=10) to infer a per-individual rate.

For demographic scenarios (Constant Population Size & OOA, **fig. S2AB**) where population C serves as an unadmixed outgroup, TRACE, *hmmix* and *Sprime* perform reliably with high precision and recall, and low FDR (**fig. S7AB**). In contrast, in demographic scenarios where population C does not function as an unadmixed outgroup—such as the Archaic Introgression before OOA Model where populations A and C carry equal amounts of archaic ancestry (**fig. S2C**), and the Gene Flow between Modern Humans after OOA Model, where post-OOA gene flow introduces introgressed segments into the outgroup (**fig. S2D**)—*hmmix* and *Sprime* can be biased by even small amounts of archaic gene flow into the outgroup while TRACE remains reliable. In these cases, *hmmix* and *Sprime* recover less than 20% ground-truth introgression tracts at around 80% precision, while TRACE remains robust – recovering over 50% introgression signals and high (90%) precision and low FDR (**fig. S7CD**). In practical applications using inferred ARGs, performance of TRACE may be lower depending on the accuracy of the underlying genealogical inference (**fig. S10**).

#### S2.5 Estimating Divergence time and Admixture Time

The identified archaic segments also contain information about the timing of admixture ($T_{admix}$) and the split time ($T_{archaic}$) of the introgressing population from other extant lineages within the ARG. We can estimate bounds for these times from the long branches in the ARGs. Under an instantaneous admixture model, the more recent endpoint (lower-end) of a long branch provides a lower-bound for $T_{admix}$, since most introgressed segments coalesce with each other at or shortly after the introgression event. Conversely, the older endpoint (upper-end) of the long branch provides an upper-bound estimate for $T_{archaic}$, as the coalescence between the introgressing archaic lineage and the non-introgressed modern human lineage must have occurred before population divergence. We note, however, that substantial stochastic variation in the coalescent process leads to considerable uncertainty in $T_{archaic}$ estimates.

Using the upper-end and lower-end times of introgression branches in marginal trees, we provide an approximate estimate for $T_{admix}$ and $T_{archaic}$. We report mean values of upper-end and lower-end times for the introgression branch from marginal trees embedded in TRACE-inferred archaic segments from simulations under the Constant Size Model (**fig. S8**). We show that using confident archaic segments (> 0.05cM), this approach gives a robust lower-bound estimation for $T_{admix}$ and upper-bound estimation for $T_{archaic}$.

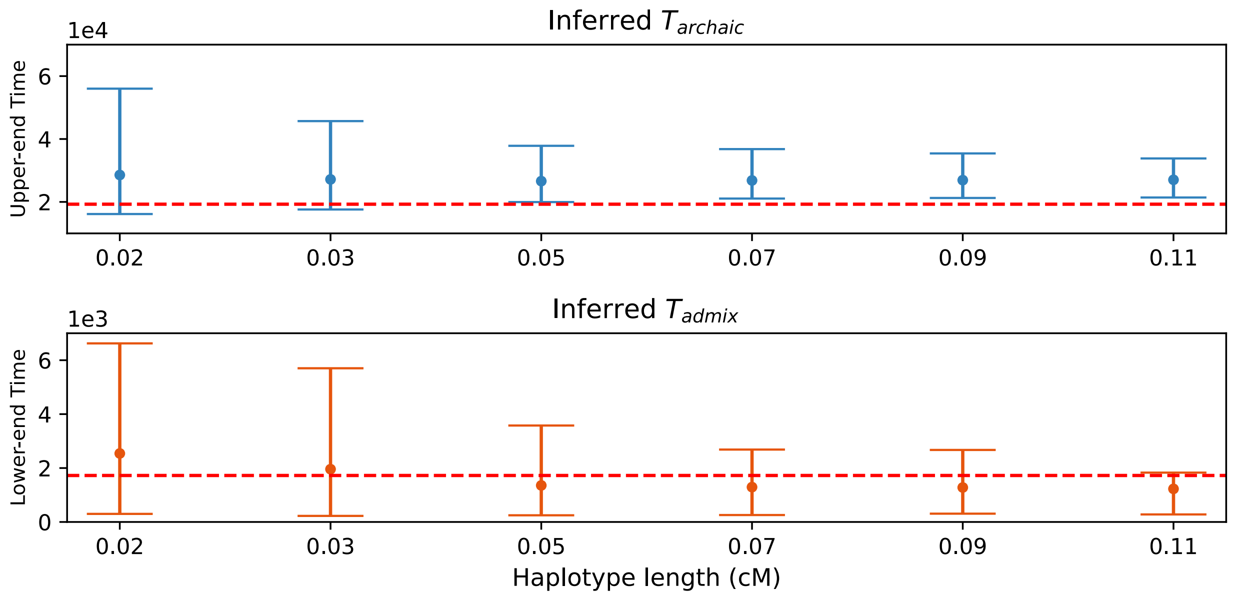

**Fig S8**. Estimating divergence and admixture times from TRACE-inferred segments. We applied TRACE to ground-truth ARGs simulated under the Constant Size Model (fig. S2A) with 2% archaic introgression. For each detected segment, we extracted the upper- and lower-end times from all introgression branches that the segment embedded. Divergence time ($T_{archaic}$) and admixture time ($T_{admix}$) were estimated by taking the average across all segments. Error bars reflect the 95% confidence intervals of the estimated times across all evaluated segments.

### Section S3. Benchmarking performance of TRACE with inferred ARGs for archaic introgression detection

#### S3.1 Simulation and performance evaluation

We used the same four demographic models as in fig. S2. For each model, we generated 10 replicate simulations of 50 Mbp genomes and sampled 100 individuals each from populations A and C. The VCF output containing all genotype information for these 200 sampled individuals per simulation served as the input for inferring ARGs using *Relate* (*38*) and *SINGER* (*36*).

For all simulations, we provided *Relate* with a constant recombination rate of `10-8` per bp per generation. We ran *Relate* (v1.1.5_x86_64_static) with the parameters `-m 1.2e-8 -N 1e4`. The resulting ARGs were then processed with the EstimatePopulationSize.sh script (using `-m 1.2e-8`) to re-estimate branch lengths. Finally, we converted the output to *tskit* (*79*, *80*) format using *Relate*'s Convert utility.

We ran *SINGER* with the parameters `-m 1.2e-8 -n 200 -thin 100 -Ne 2e4 -polar 0.99` for all simulations. For downstream analyses, we used the 150th to 199th posterior tree sequences from the *SINGER* output to ensure proper mixing of the Markov chain Monte Carlo (MCMC) sampler. For each of these posterior tree sequences, we extracted the observation data required for TRACE from every marginal tree. We then summarized this data into 1000 bp windows by calculating a weighted mean (weighted by the genomic span of each tree) of the observations across all marginal trees falling within each window, resulting in one data point per window. Finally, we averaged the observation data across all 50 posterior tree sequences per window to generate the final input for TRACE.

We applied TRACE with $t = 10000, 15000$ generations to the true ARGs, the *Relate*-inferred ARGs, and the *SINGER*-inferred ARGs. For the Constant Size Model (**fig. S2A**), we used ARGs including individuals from only population A. For the OOA model and its extensions (**fig. S2BCD**), we used the joint ARG for individuals from populations A and C to mitigate the impact of population bottleneck in population A. Results were filtered using a posterior probability threshold of 0.9.

We evaluated the performance of TRACE by measuring precision, recall, and FDR (**Section S1.3**). We applied haplotype length cutoffs ranging from 0.01cM to 0.15cM and measured performance statistics for 10 haplotypes (5 individuals) from population A in each simulation, yielding 100 total data points (10 haplotypes x 10 simulations). Standard errors were measured using 30 bootstrap resampling iterations, where each iteration involved resampling 90% of the data.

#### S3.2 Performance of TRACE using inferred ARGs

As shown in fig. S9 and S10, TRACE maintains high precision (~90%) and a low false discovery rate (FDR < 0.25%) on ARGs inferred by both *Relate* and *SINGER* at a haplotype length threshold of 0.05cM. However, recall is substantially lower on inferred ARGs compared to true ARGs.

Under the OOA model (**fig. S2B**), TRACE recovers less than 10% of true segments using *Relate* and approximately 50% using *SINGER*, compared to ~80% recall on true ARGs (**fig. S9**). Furthermore, recall decreases under more complex demographic models, with performance declining from the Constant Size Model to the OOA model and its extensions (**fig. S10**). Despite these challenges, TRACE successfully recovered over 30% of archaic segments from *SINGER*-inferred ARGs even under complex demographies, while maintaining high precision (~90%) and low FDR (< 0.25%).

We estimated both the divergence time ($T_{archaic}$) and introgression time ($T_{admix}$) using TRACE on inferred ARGs (**Section S1.4, fig. S11**). Across all demographic models, TRACE provided a robust upper bound for $T_{archaic}$ using inferred ARGs, as seen with true ARGs. For $T_{admix}$, unlike with true ARGs, we observed a systematic upward bias with inferred ARGs—particularly those from *SINGER*—due to biases in coalescent time estimation (*41*).

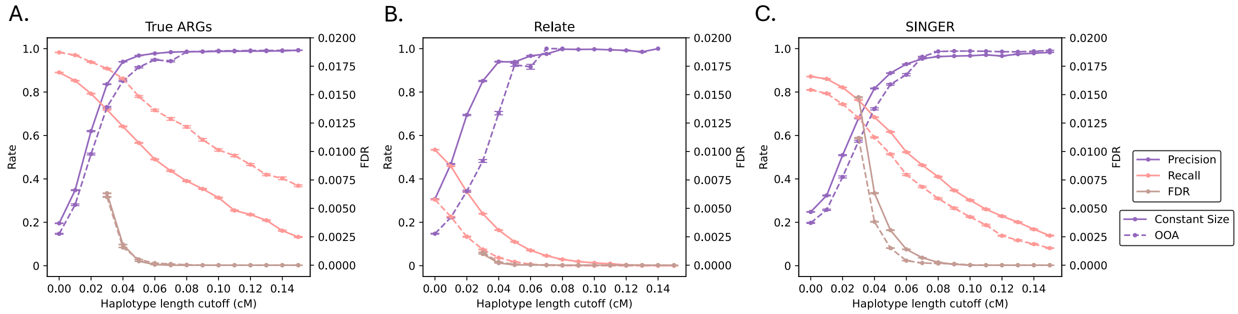

**Fig S9**. Evaluation of TRACE with inferred ARGs. Performance metrics (precision: purple, recall: pink, FDR: brown) versus haplotype length cutoff for (A) ground-truth, (B) *Relate*-inferred, and (C) *SINGER*-inferred ARGs. Solid and dashed lines represent Constant Size and OOA models, respectively. Error bars show standard error across 30 bootstrap replicates.

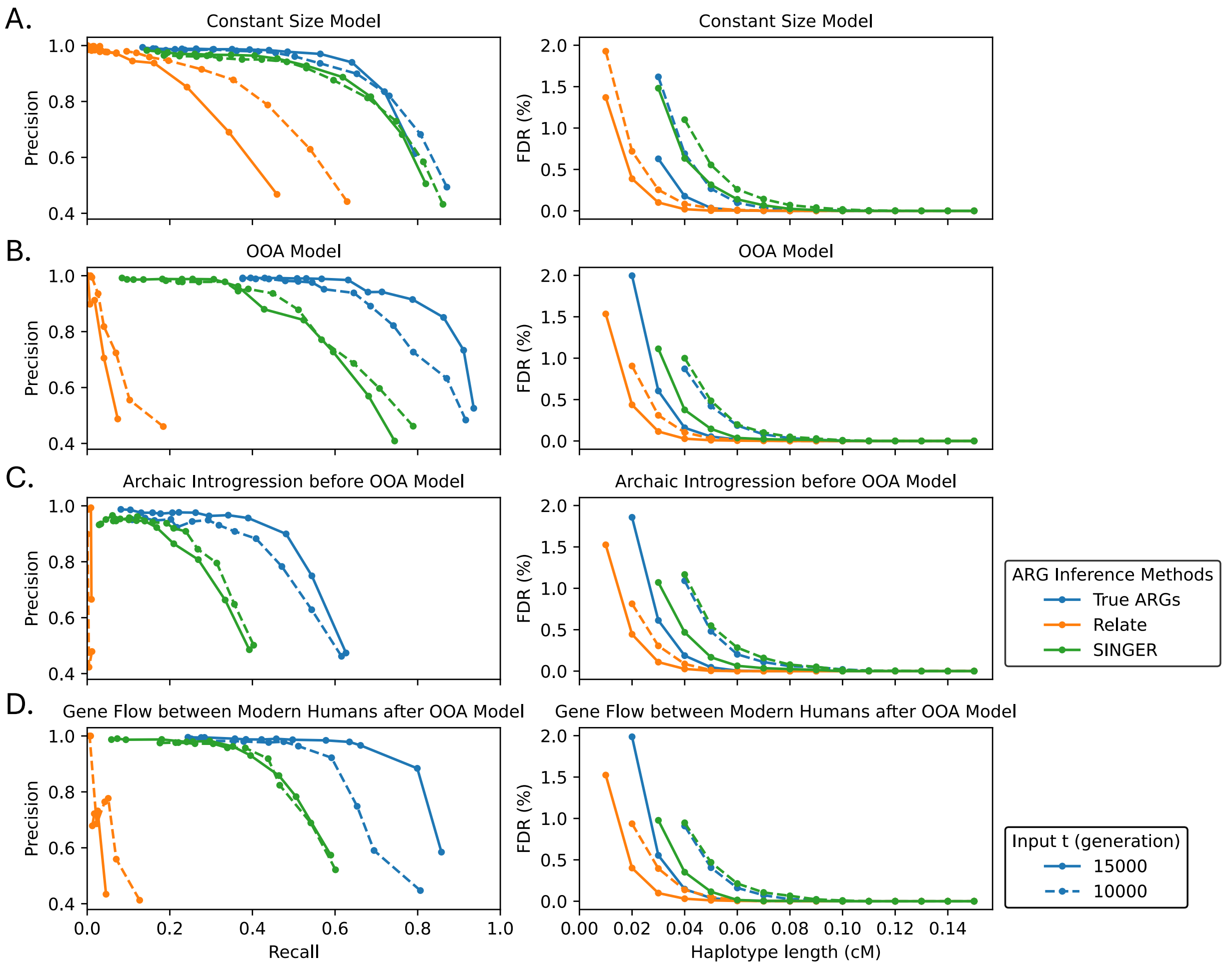

**Fig S10.** Performance of TRACE using inferred ARGs. Precision-recall curves (left) and false discovery rate (right) for TRACE applied to ground-truth (blue), *Relate*-inferred (orange), and *SINGER*-inferred (green) ARGs. Results are shown across four demographic models with 2% archaic introgression: **(A)** Constant Size, **(B)** OOA, **(C)** Archaic Introgression before OOA, and **(D)** Gene Flow between Modern Humans after OOA.

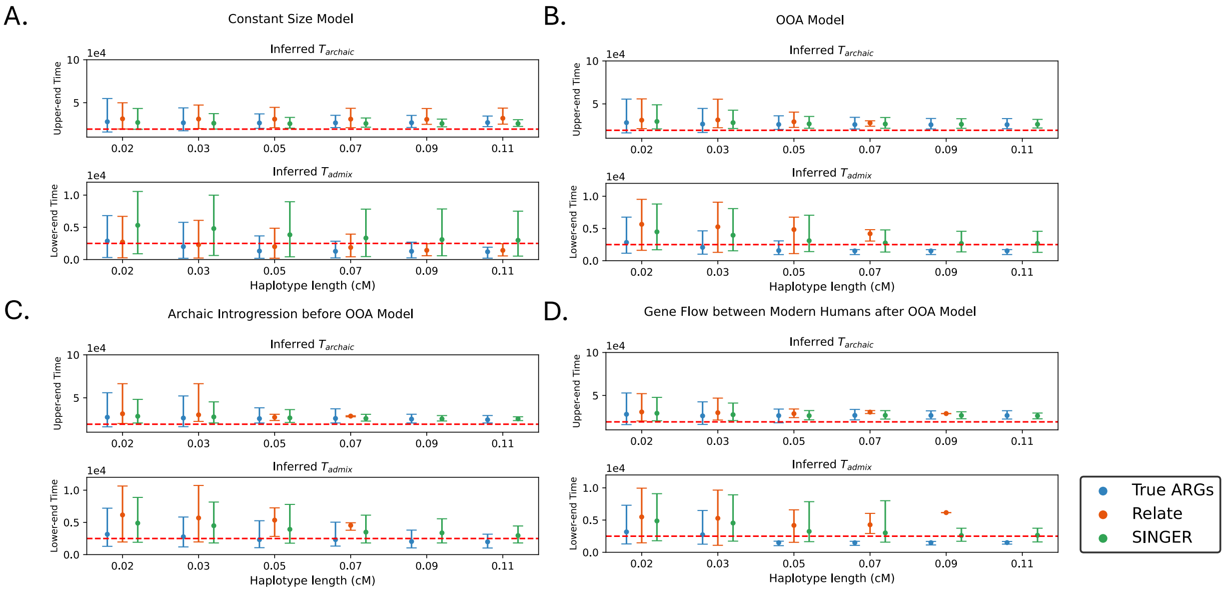

**Fig S11**. Estimating divergence and admixture times using TRACE. Results are shown for ground-truth (blue), *Relate*-inferred (orange), and *SINGER*-inferred (green) ARGs across four demographic models with 2% archaic introgression: **(A)** Constant Size, **(B)** OOA, **(C)** Archaic Introgression before OOA, and **(D)** Gene Flow between Modern Humans after OOA. Missing *Relate* data indicate scenarios where no recovered segments passed the haplotype length cutoff when applying TRACE to Relate-inferred ARGs. Error bars reflect the 95% confidence intervals of the estimated times across all evaluated segments.

#### S3.3 Benchmarking the accuracy of ARGs reconstruction for features related to archaic introgression

To better understand why TRACE performance differs between inferred and true ARGs, we benchmarked the accuracy of ARG inference for features critical to detecting archaic introgression. We focused on two key summaries: (1) branch lengths, specifically the "long branch" signal characteristic of archaic introgression, and (2) tree topologies that reflect the presence of introgressed lineages.

Using simulations under the Constant Size Model, we first compared the distributions of TRACE's observation data across true ARGs and inferred ARGs using *SINGER* and *Relate*. For each individual in population A, we extracted branches spanning $t = 15000$ generations from marginal trees and labeled them as "Modern Human" or "Archaic" based on the true ancestry (known in simulations). We then visualized the distributions of three key features: branch length ($L^{(t)}$), number of coalescence events ($N^{(t)}$), and the composite observation data ($X^{(t)} = L^{(t)} \times N^{(t)}$) used by TRACE. As shown in fig. S12, *SINGER* recovers branch lengths and tree topologies associated with the introgression signal more accurately than *Relate*, which explains TRACE's superior performance on *SINGER*-inferred ARGs. To quantify the separation between "Modern Human" and "Archaic" distributions, we calculated the Kolmogorov–Smirnov (KS) distance between emissions related to the two state distributions using `scipy.stats.ks_2samp`. *SINGER*-inferred ARGs nearly replicate the KS distances observed in true ARGs for all features (**fig. S12**), whereas *Relate*-inferred ARGs have trouble distinguishing emission distributions between two states due to topology errors.

We also assessed the accuracy of coalescent time estimation in inferred ARGs by analyzing pairwise time to the most recent common ancestor (TMRCA) between a randomly selected haplotype and all other haplotypes in the ARG across the genome (**fig. S13**). We separated TMRCAs originating from true "Modern Human" versus "Archaic" regions. For each distribution, the probability mass between two time points $T_{admix}$ and $T_{archaic}$ was calculated as the proportion of TMRCAs falling within that interval. Figure S13 further supports that *SINGER* more accurately recovers the characteristic "long branch" signal of introgression in marginal trees, demonstrated by a more pronounced reduction in probability mass between the $T_{admix}$ and $T_{archaic}$ as well as a higher correlation (Pearson's $r$) with ground-truth TMRCAs.

Focusing specifically on the "introgression branch" within marginal trees from admixed segments, we compared the lower-end and upper-end coalescence times between the true ARGs and the inferred ARGs (**fig. S14**). We found that both *SINGER* and *Relate* overestimated the lower-end times ($T_{admix}$) and underestimated the upper-end times ($T_{archaic}$), though the magnitude of this bias differed between the methods—*SINGER* recovers the upper-end branch time with much higher accuracy than *Relate* while both methods exhibited systematic bias in estimating the lower-end time. We quantified the correlation between the true and estimated times using Pearson's $r$. These results demonstrate that the previously described "branch shrinking" bias in ARG reconstruction methods (*41*, *81*) occurs at different timescales and can significantly impact the accuracy of archaic introgression detection.

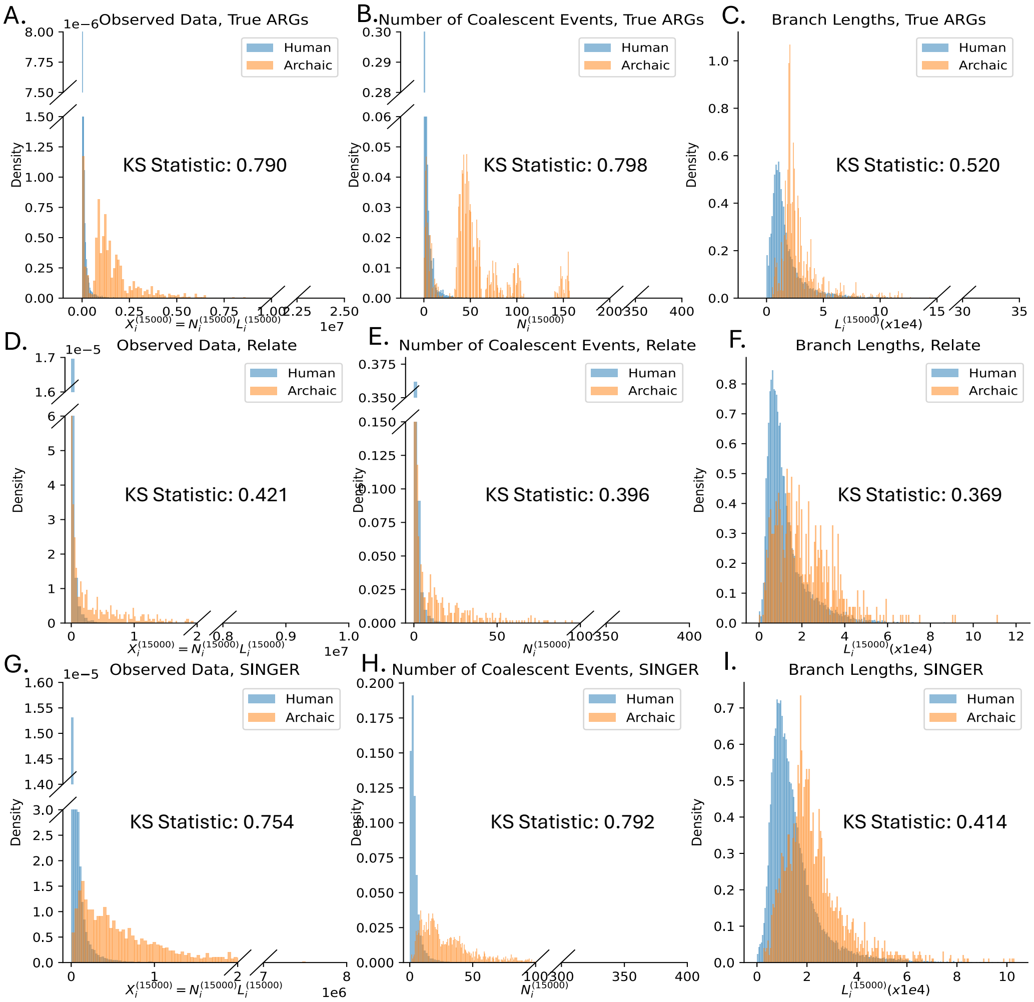

**Fig S12.** Summary statistics of TRACE features from simulated data (Constant Size Model, 2% introgression proportion). Analysis of a single target haplotype, showing the branch subtending the target at $t = 15000$ generations. Distributions are shown for: (**A, D, G**) branch length ($L^{(t)}$), (**B, E, H**) number of external coalescent events ($N^{(t)}$), and (**C, F, I**) the compound statistic $X^{(t)} = L^{(t)} \times N^{(t)}$. Rows represent ground-truth, *Relate*-inferred, and *SINGER*-inferred ARGs, respectively. Colors indicate true ancestry (modern human / archaic).

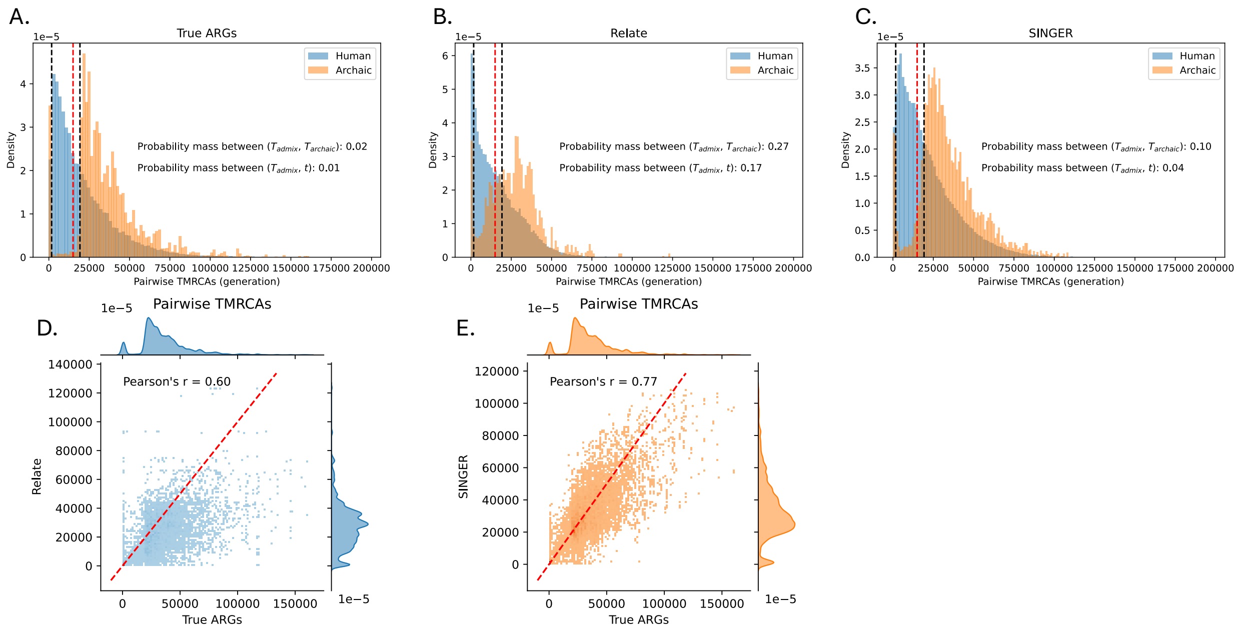

**Fig S13.** Pairwise TMRCA distributions and accuracy in inferred ARGs. Analysis of a single haplotype from a single simulation (Constant Size Model, 2% introgression proportion). (**A-C**) Genome-wide TMRCA distributions between the target haplotype and all other samples in the ARG, colored by true archaic ancestry. Vertical lines indicate key time points: $T_{admix}$ and $T_{archaic}$ (black), and the input parameter $t = 15000$ generations (red). (**D-E**) Scatterplots comparing TMRCA estimates for the same sample pairs and loci between ground-truth (x-axis) and inferred (y-axis) ARGs, with the y=x line in red.

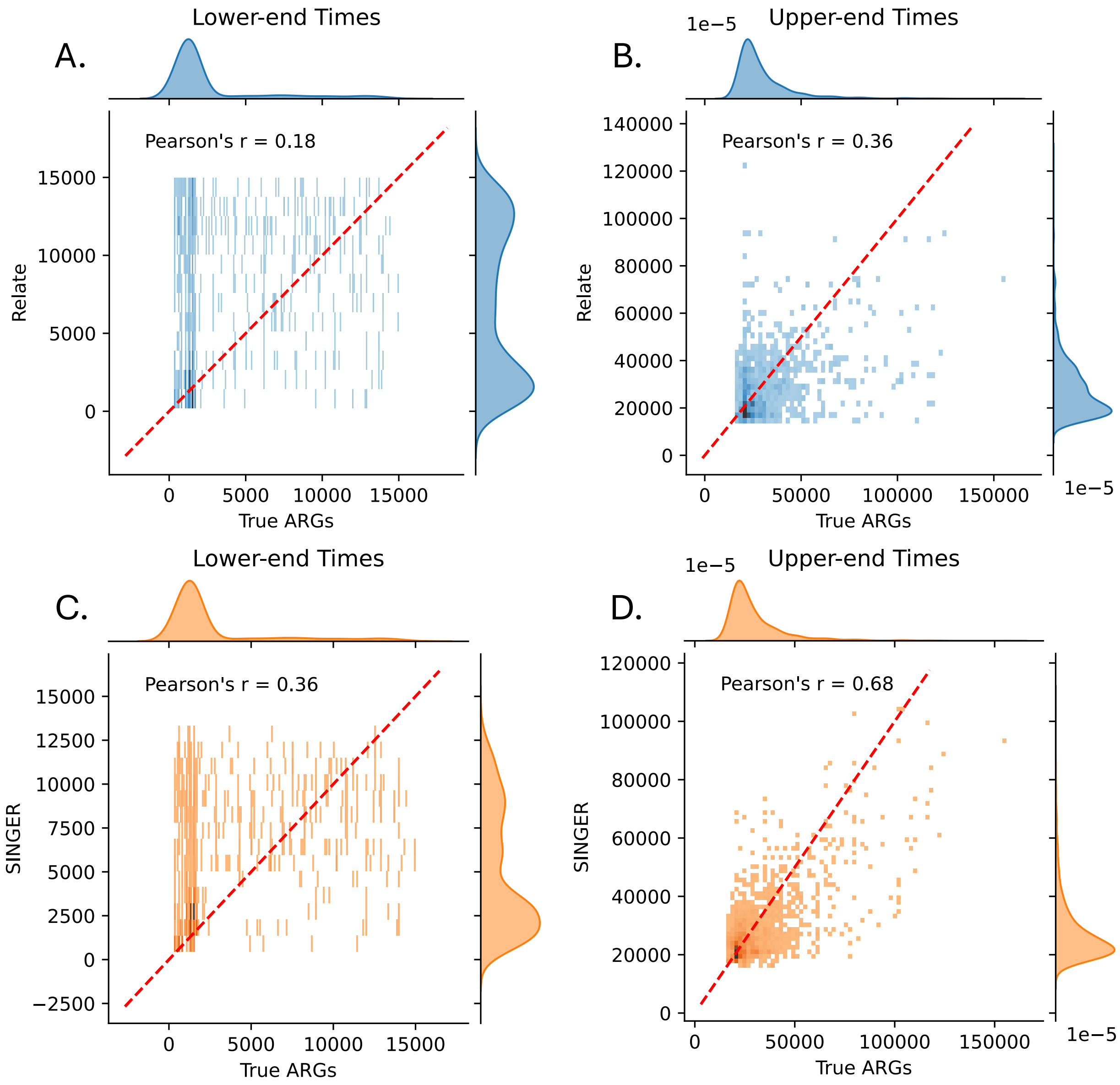

**Fig S14.** Estimated introgression branch times in inferred ARGs. Comparison of upper- and lower-end branch times between true ARGs (X-axis) and inferred ARGs (Y-axis) in one simulation under a Constant Size Model with 2% introgression proportion. Results are shown for: (A-B) *Relate*-inferred and (C-D) *SINGER*-inferred ARGs. All analyses are based on branches subtending a single target haplotype at $t = 15000$ generations.

### Section S4. Inferring archaic introgression signal on 1000 Genomes data

#### S4.1 ARGs reconstruction and archaic inference methods settings

We used high-coverage whole genome sequences from the 1000 Genomes Project (*48*) and selected 108 Yoruba in Ibadan, Nigeria (YRI), 99 Luhya in Webuye, Kenya (LWK), 91 British From England and Scotland (GBR), 103 Han Chinese in Beijing, China (CHB) and 102 Indian Telugu in the UK (ITU) individuals for analysis. To minimize the impact of the Out-of-Africa (OOA) bottleneck on archaic inference in non-African populations, we ran *SINGER* separately for each non-African population (GBR, CHB, and ITU) together with YRI samples; for African populations, we ran *SINGER* with just individuals from that population only (5 independent *SINGER* runs in total, **table S3**). We used *bcftools* (*83*) to retain only biallelic SNPs and polarized alleles to ancestral versus derived (instead of reference versus alternative) using the human ancestral genome (**table S1**). We then ran *SINGER* with `-m 1.25e-8 -n 300 -thin 100 -Ne 2e4 -polar 0.99` and used the $250^{th}$ to $299^{th}$ posterior tree sequences for all downstream analyses.

We extracted observation data from TRACE (number of coalescent events and branch lengths) from the 50 posterior tree sequences. We then applied the 1000 Genomes strict mask and retained only marginal trees where over 99% of their span overlapped with accessible regions in the strict mask (*pybedtools* (*84*), **Section S1.2**). We summarized the observation data into 1000 bp windows by calculating a weighted mean (weighted by the span of each tree) of the observations across all marginal trees within the same genomic window. This yielded one data point per window for each posterior tree sequence. The final input for TRACE was generated by averaging this summarized data across all 50 posterior tree sequences.

We ran TRACE with $t = 15{,}000$ generations to detect archaic introgression. The inferred archaic ancestry tracts were filtered to retain those with posterior probability > 0.9, haplotype length > 50kbp and > 0.05cM based on genetic distances in HapMap genetic map (*85*) (**table S1**). For each filtered archaic ancestry tract, we examined the number of derived mutations (within the 1000 Genomes strict mask) that were also sequenced in high-coverage archaic genomes using three Neanderthals (Altai, Vindija, Chagyrskaya) and Altai Denisovan (Denisova 3). We retained only tracts containing more than 30 derived mutations and for which at least 10 SNPs were also sequenced in the archaic genomes (**table S1**).

For comparison, we applied *hmmix* to 1000 Genomes data using 426 individuals including Yoruba in Ibadan, Nigeria (YRI), Esan in Nigeria (ESN) and 64 Africans from HGDP who have less than 1% West Eurasian ancestry, including Bantu South Africa, Biaka Pygmy, Mbuti Pygmy, San and Yoruba as outgroups. We applied 1000 Genomes strict mask to estimate the number of callable sites in the outgroup, the SNP density (as a proxy for per-window mutation rate) and the number of private variants in the test population (GBR, CHB or ITU) in 1kb windows across the genome. We filtered the *hmmix* output to retain segments with posterior probability > 0.8, haplotype length > 50kbp and > 0.05cM. To identify the source of archaic ancestry as "Neanderthal" or "Denisovan", we used the four high-coverage archaic genomes—Altai, Vindija, Chagyrskaya, and

Denisova (**table S1**). Following documentation, a segment was labeled "Neanderthal" if it shared more derived alleles with the sequenced Neanderthals (Altai, Vindija, Chagyrskaya) than with the sequenced Denisovan; conversely, it was labeled "Denisovan" if it shared more with the Denisovan genome. Segments with an equal number of shared alleles or no shared alleles were labeled as "Ambiguous" (*53*).

We ran *IBDmix* using the Altai Neanderthal and Altai Denisovan genomes as the archaic reference panels. We obtained the accessible genome mask from (*28*) and lifted it over from hg19 to hg38 using *CrossMap* (*86*). The *IBDmix* output was filtered for segments with `slod` > 4, haplotype length > 50kbp and > 0.05cM. Segments passing these filters that were identified using the Altai Neanderthal reference genome were labeled as "Neanderthal", while those identified using the Altai Denisovan reference genome were labeled as "Denisovan".

#### S4.2 Classification of archaic ancestry segments detected by TRACE

We classified archaic segments detected by TRACE as "Neanderthal", "Denisovan", or "Ghost" ancestry by examining the sharing patterns of mutations that mapped onto the introgressed branches and evaluated the performance of resulting classification rules in simulations.

To map mutations to the introgression branch, we implemented the following procedure:

- Estimate mutation ages from posterior tree sequences: For each mutation in a *SINGER* posterior tree sequence, we extracted the time of the node on which the mutation occurred and the time of its parent node. The mutation age was estimated as the average of these two node times.
- Average mutation ages across posteriors: We then computed the mean mutation age across all 50 *SINGER* posterior tree sequences.
- Record introgression branch coordinates: While extracting observation data for TRACE, we recorded the lower-end and upper-end of the coalescence times for the branch spanning the input time $t$ that subtends the target haplotype in each marginal tree. These branch times were summarized into 1000 bp windows and averaged across the 50 posterior tree sequences (**Section S4.1**).
- Map mutations to the introgression branch: For each mutation within a detected archaic segment, we compared its average age to the summarized branch times of the 1000 bp window in which it is located. A mutation was classified as:
    - "On" the branch if its age falls between the upper-end and lower-end times.
    - "Above" the branch if its age was older than the upper-end time.
    - "Below" the branch if its age was younger than the lower-end time.

We categorized mutations within the archaic ancestry segments that were mapped to ("On") the introgressed branches, polarized by ancestral (=0) or derived (=1) alleles in Neanderthals (*N*) and Denisovans (*D*) as follows:

- ND10: Derived in at least one sequenced Neanderthal genome (Altai, Vindija, Chagyrskaya) and ancestral in the sequenced Denisovan genome.
- ND01: Ancestral in all sequenced Neanderthals and derived in the sequenced Denisovan genome.
- ND00: Ancestral in all sequenced Neanderthals and the sequenced Denisovan genome.

- ND11: Derived in at least one sequenced Neanderthal genome and the sequenced Denisovan genome.
- sYRI: Derived in at least one YRI individuals in the ARG.

Let $N_t, t \in \{ND10, ND01, ND11, ND00\}$ be the count of mutations of type $t$ on the introgression branch. We then defined the following statistics:

$$P_{ND01} = \frac{N_{ND01}}{N_{ND01} + N_{ND10} + N_{ND00}}$$

$$P_{ND10} = \frac{N_{ND10}}{N_{ND01} + N_{ND10} + N_{ND00}}$$

$$P_{ND00} = \frac{N_{ND00}}{N_{ND01} + N_{ND10} + N_{ND00} + N_{ND11}}$$

$$P_{sYRI} = \frac{N_{sYRI}}{N_{ND01} + N_{ND10} + N_{ND00} + N_{ND11}}$$

We developed two classification rule groups (**table S5**):

- *Rule Group 1:* A segment is classified as
    - "Neanderthal" if $P_{ND10} > 0.2$ & $N_{ND10} > N_{ND01}$
    - "Denisovan" if $P_{ND01} > 0.2$ & $N_{ND10} < N_{ND01}$
    - "Ghost" if $P_{ND00} > 0.8$
- *Rule Group 2:* A segment is classified as
    - "Ghost" if $P_{ND00} > 0.8$ & $P_{sYRI} < 0.1$
    - "Neanderthal" if ($P_{ND00} < 0.8$ | $P_{sYRI} > 0.1$) & $N_{ND10} > N_{ND01}$
    - "Denisovan" if ($P_{ND00} < 0.8$ | $P_{sYRI} > 0.1$) & $N_{ND10} < N_{ND01}$

We applied *Rule Group 1* as our primary classification scheme to identify the source of archaic ancestry segments. We applied *Rule Group 2* to identify super-archaic segments embedded in Denisovan ancestry tracks (**Section S6.3**).

We evaluated the performance of these classification schemes using simulations (see **Section S7.1** for simulation models). Let $N_d^s$ be the total number of detected tracts classified as archaic source $s$ ($where\ s \in \{NEA, DEN, Ghost\}$), $N_s$ be the number of detected archaic tracts that overlap with a ground-truth segment from source $s$ , and $N_{tp}^s$ be the number of tracts correctly classified as source $s$. We then calculated:

$$precision_s = \frac{N_{tp}^s}{N_d^s}$$

$$recall_s = \frac{N_{tp}^s}{N_s}$$

As shown in table S5, both classification rule groups achieve high overall accuracy with minimal misclassification across all archaic ancestry types. In general, *Rule Group 1* has higher precision. However, when the goal is to detect super-archaic segments embedded *within* Denisovan ancestry, *Rule Group 2* is preferable as it helps retain segments with high super-archaic ancestry as

"Denisovan" in classification, thereby increasing the overall recall of the super-archaic detection pipeline.

#### S4.3 Overlap in archaic ancestry segments between TRACE and other inference methods

We compared TRACE inference results with inference from two established archaic ancestry inference methods*: hmmix,* which is reference-free, and *IBDmix*, which utilizes an archaic reference genome. *Sprime* is not applied here as it does not provide haplotype-level ancestry painting, making it impossible to do one-to-one direct comparison with TRACE outputs. For each archaic segment detected by TRACE, we used *pybedtools* to check if it overlaps with *hmmix*- or *IBDmix*-detected segments on the same haplotype (*hmmix*) or individual (*IBDmix*). In table S4, we report the proportion of TRACE segments that overlap for at least 1 bp with those from the other methods, defined as:

$$proportion\ overlap\ = \frac{number\ of\ segments\ overlap}{total\ number\ of\ segments}$$

As shown in table S4, over 80% of Neanderthal and over 70% of Denisovan segments that TRACE detected overlap with those found by by *hmmix* or *IBDmix* (**table S4**). In contrast, less than 5% TRACE detected ghost segments are identified as Neanderthal or Denisovan segments by either of the two methods. We note that without an outgroup or ghost reference genome, ghost segments are unlikely to be reliably detected by *hmmix* or *IBDmix*.

We note that TRACE generally recovers less total Neanderthal and Denisovan ancestry than *hmmix* and *IBDmix*, as well as some previous estimates (*28*, *53*, *54*) (**fig. S15**). This likely reflect lower recall of TRACE using inferred ARGs as seen in simulations (**fig. S10**).

We evaluated the affinity (the proportion of derived mutations that are shared with the sequenced archaic genomes) of detected segments to Neanderthal or Denisovan origins (**fig. S16**). The vast majority of segments shared between TRACE and the other methods exhibit high Neanderthal or Denisovan affinity (**fig. S16AB**). Conversely, segments uniquely detected by TRACE predominantly show identical affinity to both sequenced Neanderthal and Denisovan genomes, consistent with their classification as originating from an unknown archaic source distantly related to Neanderthals and Denisovans.

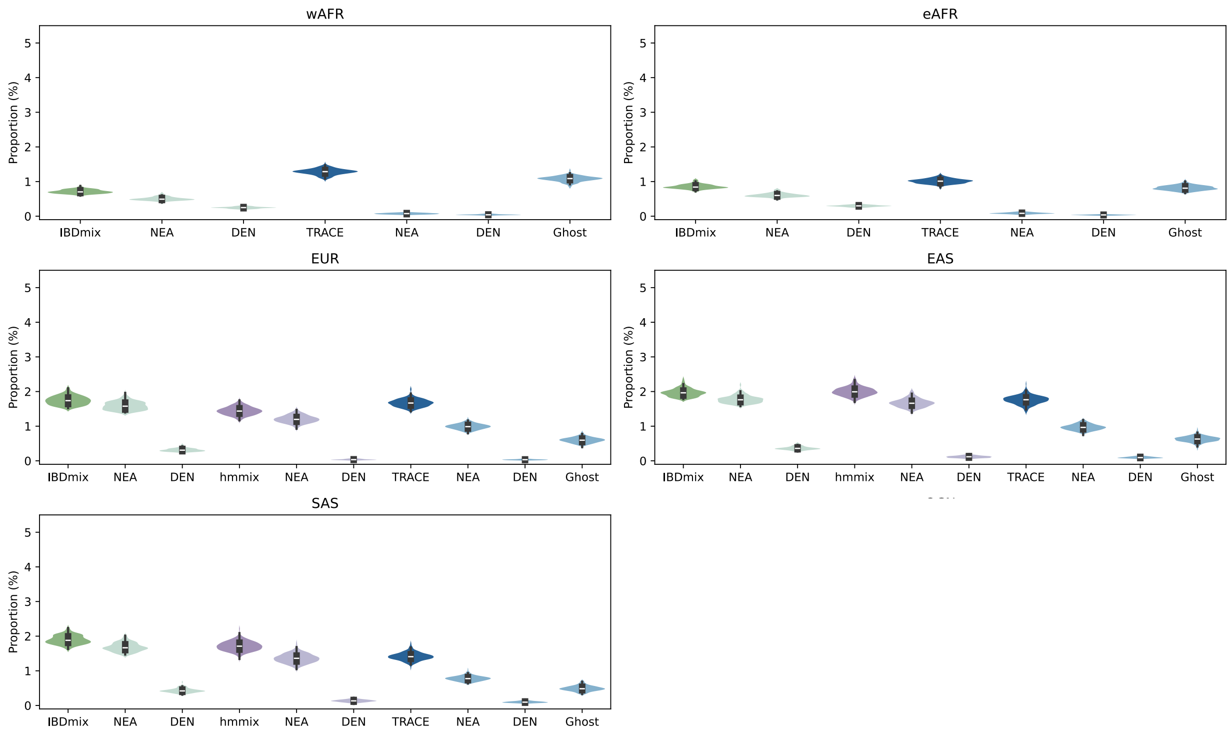

**Fig S15**. Per-individual archaic ancestry in 1000 Genomes populations. Total archaic ancestry (dark colors) and its composition (lighter colors: Neanderthal-NEA, Denisovan-DEN, Ghost) estimated by TRACE (blue), IBDmix (green), and hmmix (purple).. All methods used a consistent length filter (> 50kbp, > 0.05cM). TRACE was run with $t = 15000$ and posterior probability > 0.9; *hmmix* with posterior > 0.8; and *IBDmix* with `slod` > 4. wAFR = West Africans (YRI), eAFR = East Africans (LWK), EUR = Europeans (GBR), EAS = East Asians (CHB) and SAS = South Asians (ITU) in 1000 Genomes data.

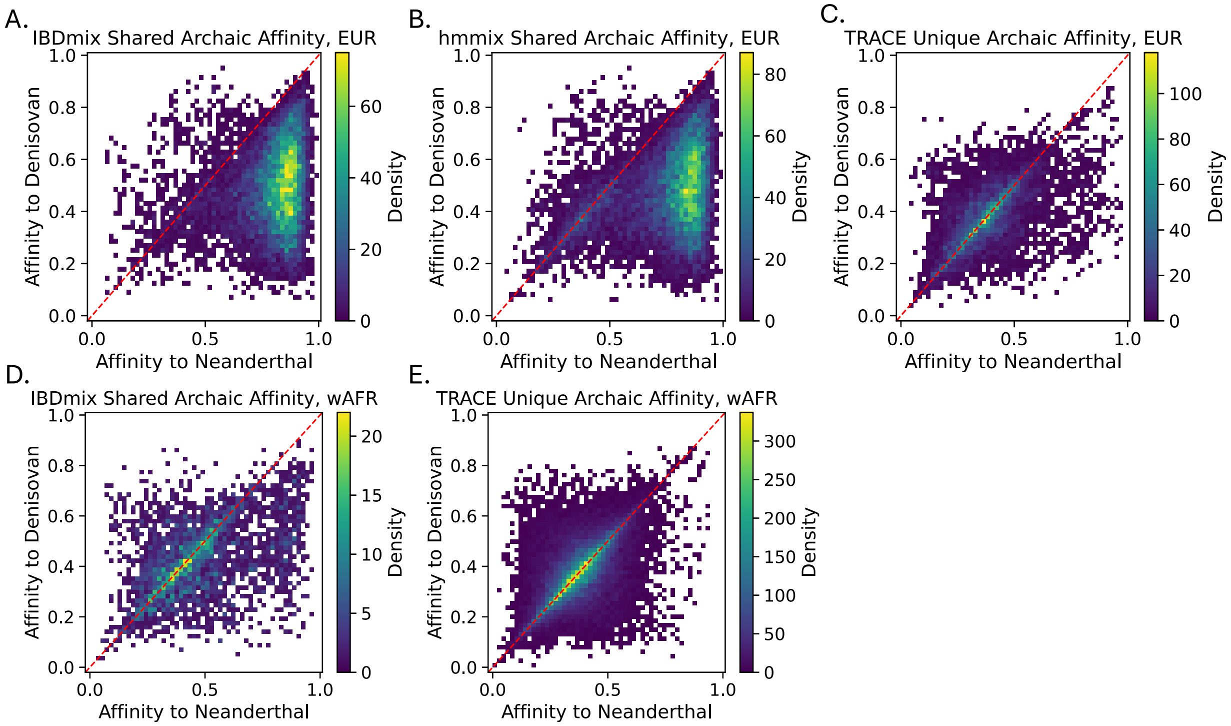

**Fig S16**. Archaic affinity of introgressed segments. Neanderthal versus Denisovan genetic affinity for TRACE-detected segments, categorized by their concordance with other methods and population origin: **(A)** *hmmix*-overlapping (EUR), **(B)** *IBDmix*-overlapping (EUR), **(C)** TRACE-specific (EUR), **(D)** *IBDmix*-overlapping (AFR), **(E)** TRACE-specific (AFR). We do not apply *hmmix* to AFR due to a lack of outgroup population.

#### S4.4 Assessing archaic affinity, site frequency spectrum (SFS) and conditional SFS (cSFS) of segments

For segments detected by TRACE, we quantified their Neanderthal and Denisovan affinity by calculating the proportion of derived mutations that are shared with the sequenced archaic genomes. This analysis was restricted to mutations within the 1000 Genomes strict mask (note that unlike in Section S4.3, here we are using all mutations on the segments, not just mutations on the introgression branch). Let $N_t$, where $t \in \{ND10, ND01, ND00, ND11\}$, be the count of mutations of type $t$ on a segment. We then defined:

$$Neanderthal\ affinity = \frac{N_{ND10} + N_{ND11}}{N_{ND01} + N_{ND10} + N_{ND00} + N_{ND11}}$$

$$Denisovan\ affinity = \frac{N_{ND01} + N_{ND11}}{N_{ND01} + N_{ND10} + N_{ND00} + N_{ND11}}$$

We report the Neanderthal and Denisovan affinity for all detected segments (**fig. S17**). As expected, segments classified as "Ghost" display equal affinity to both Neanderthal and Denisovan genomes, consistent with their origin in an unsampled archaic lineage. This finding is robust to the number of Neanderthal reference genomes used in the classification, as shown by the nearly identical results obtained with one (Vindija only, **fig. S18**) versus three Neanderthal genomes (**fig. S17**) used in the analysis.

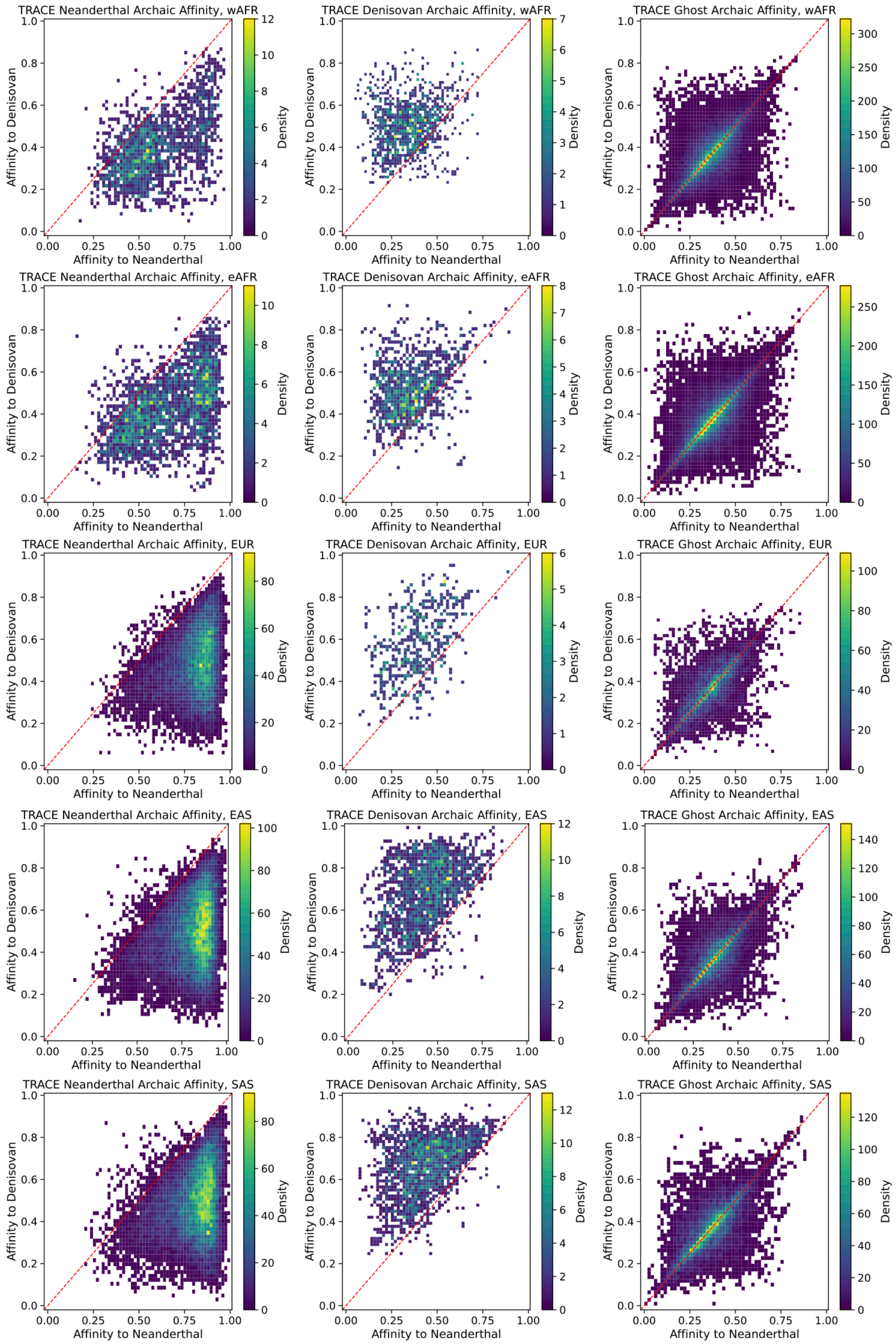

**Fig S17**. Neanderthal versus Denisovan genetic affinity for archaic segments. TRACE-detected segments are classified as Neanderthal (left column), Denisovan (middle), and Ghost (right) ancestry. Rows represent populations: West Africans (row 1), East Africans (row 2), Europeans (row 3), East Asians (row 4), and South Asians (row 5).

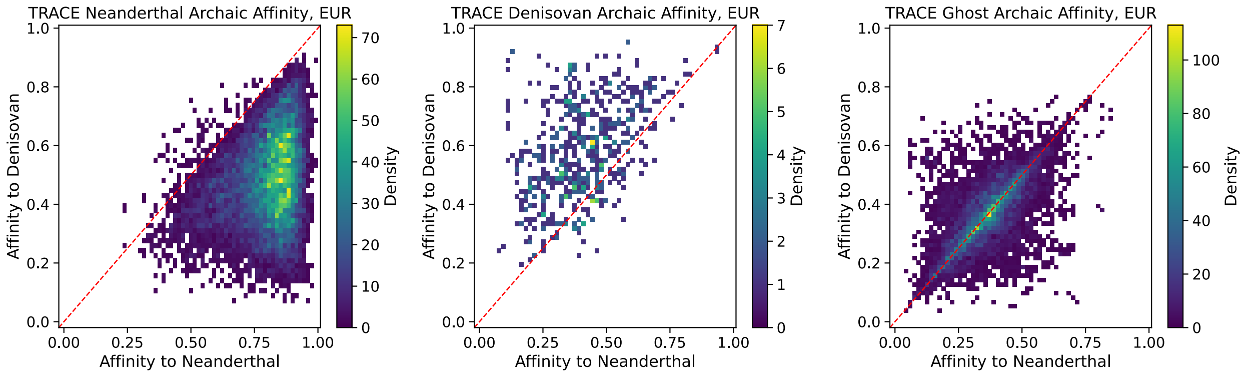

**Fig S18.** Neanderthal versus Denisovan genetic affinity for archaic segments in Europeans using Vindija Neanderthal only. TRACE-detected segments are classified as Neanderthal (left column), Denisovan (middle), and Ghost (right) ancestry.

We next analyzed the Site Frequency Spectrum (SFS) and Conditional SFS (cSFS) of the detected archaic segments. For a given archaic source $s \in \{NEA, DEN, Ghost\}$, the SFS represents the derived allele frequency distribution of all mutations in the segments assigned to the source $s$. The cSFS, different from SFS, represents the derived allele frequency distribution for mutations shared with a specific archaic source: mutations shared with Neanderthals (ND10 and ND11) when conditioning on Neanderthals (**fig. S19**), and mutations shared with Denisovans (ND01 and ND11) when conditioning on Denisovans (**fig. S20**). All histograms use a bin width of 0.02.

The SFS and cSFS for Neanderthal and Denisovan segments recapitulate the characteristic "U-shape" reported previously (*13*), with a pronounced elevation in low-frequency bins (**fig. S19-S20**). In contrast, ghost segments exhibit distinct patterns: their SFS shows a sharp elevation in low-frequency bins in Africans, which becomes more dispersed in non-Africans, while their cSFS displays a consistent, sharp elevation exclusively in high-frequency bins across all populations. These patterns are robust to conditioning on Neanderthal or Denisovan (**fig. S19 vs. S20**) and the number of Neanderthal genomes used in the analysis (**fig. S21**).

We interpret these findings as consistent with a model of ghost introgression prior to the OOA dispersal. This event introduced two classes of variants:

1. New mutations from the ghost lineage, which were absent from the modern human gene pool, and subsequently result in elevated low-frequency bins in the SFS. These mutations do not appear in the cSFS, as they are not shared with sequenced archaics (Neanderthals or Denisovans). The OOA bottleneck subsequently alters the distribution of these variants in non-Africans, causing loss or frequency shifts and resulting in a more dispersed low-frequency signal in GBR, CHB and ITU (but not in YRI or LWK).
2. Ancestral polymorphisms, which were shared among modern human and archaic lineages prior to the ghost lineage's divergence, elevating high-frequency bins in both the SFS and cSFS.

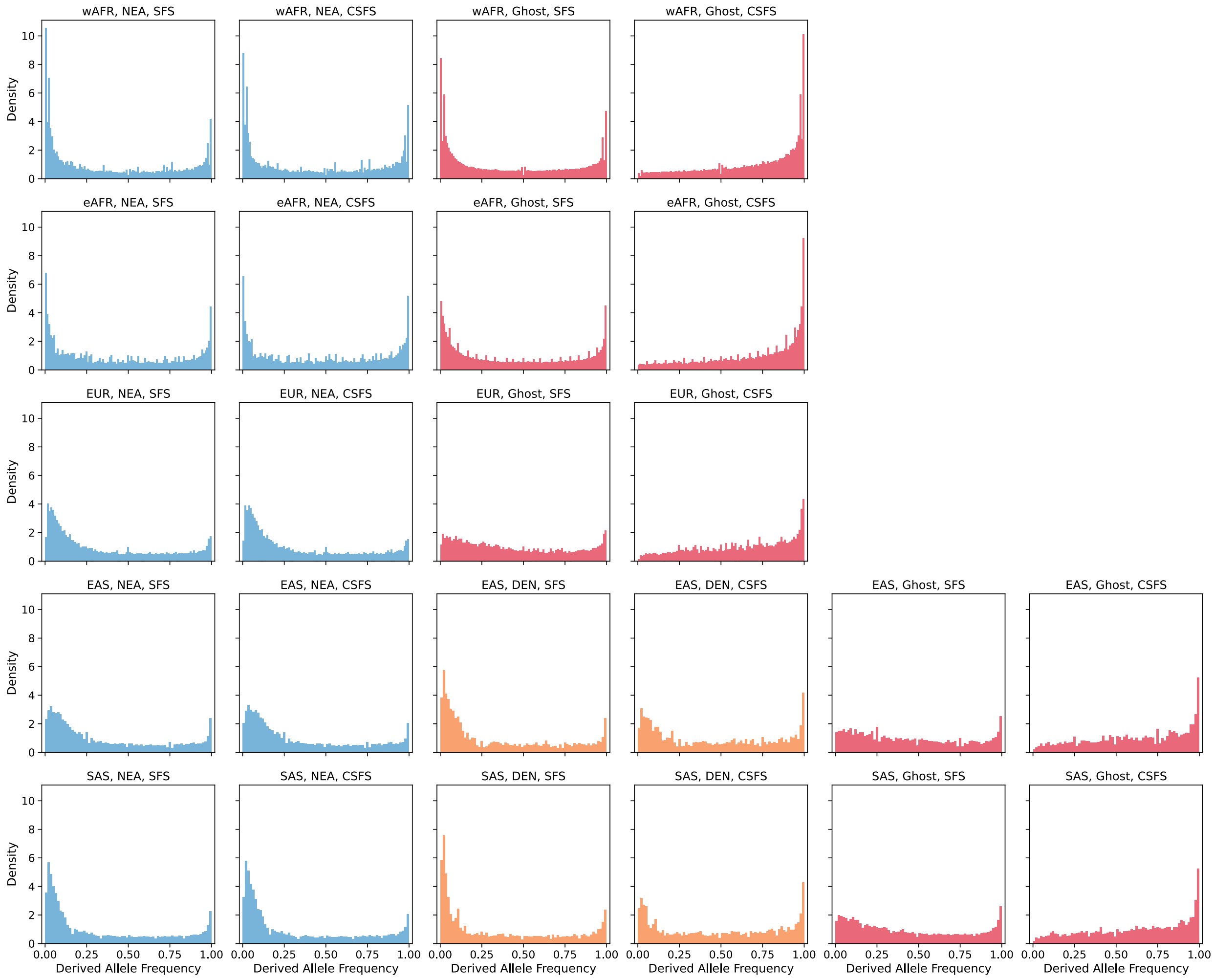

**Fig S19**. (conditional on Neanderthal) Site frequency spectra of archaic segments. Rows represent populations: West Africans, East Africans, Europeans, East Asians, and South Asians. For each population, the site frequency spectrum (SFS, left column) and conditional SFS (cSFS, right column) are shown for alleles on TRACE-detected segments. Colors indicate the assigned segments archaic ancestry: Neanderthal (blue), Denisovan (orange), and Ghost (red). The cSFS is conditioned on the derived allele being present in at least one sequenced Neanderthal genome.

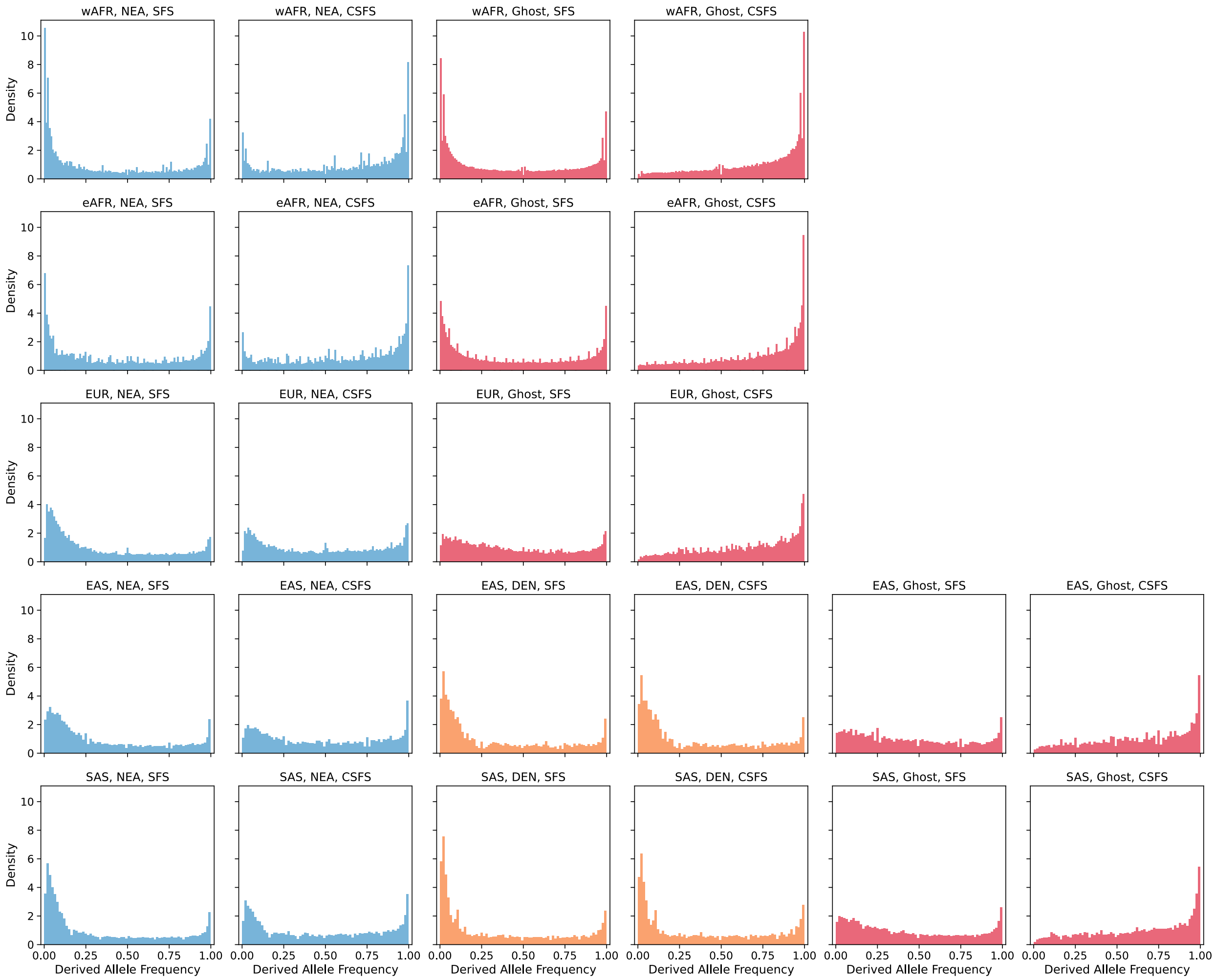

**Fig S20.** (conditional on Denisovan) Site frequency spectra of archaic segments. Rows represent populations: West Africans, East Africans, Europeans, East Asians, and South Asians. For each population, the site frequency spectrum (SFS, left column) and conditional SFS (cSFS, right column) are shown for alleles on TRACE-detected segments. Colors indicate the assigned segments ancestry: Neanderthal (blue), Denisovan (orange), and Ghost (red). The cSFS is conditioned on the derived allele being present in the Altai Denisovan genome.

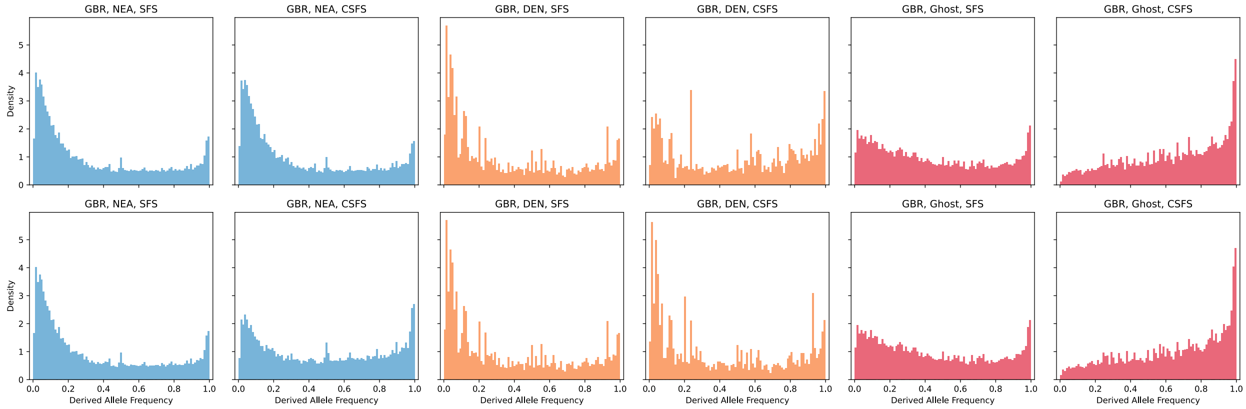

**Fig S21.** Derived allele frequency spectra of archaic segments in GBR, using only Vindija Neanderthal in TRACE inference and allele sharing pattern determinations. (Left) SFS and (Right) cSFS conditioned on the derived allele being present in Vindija Neanderthal (top) or Altai Denisovan (bottom). Colors indicate TRACE-assigned ancestry: Neanderthal (blue), Denisovan (orange), Ghost (red).

#### S4.5 Distribution, sharing and heterozygosity of ghost segments among modern humans

To assess the sharing of ghost segments across populations, we first defined, for each population $Y$ , a set $G_Y$ representing all genomic positions covered by ghost segments in at least one individual from $Y$. For each individual $i$ in population $X$, we calculated their total ghost ancestry in base pairs ($T_{X(i)}$) and the amount overlapping with $G_Y$ ($S_Y^{X(i)}$). The proportion of sharing between individual $i$ in population $X$ with population $Y$ was then calculated as $P_Y^{X(i)} = \frac{S_Y^{X(i)}}{T_{X(i)}}$.

Figure 2B and S22 show the distributions of $T_{X(i)}$ and $S_Y^{X(i)}$. We note $G_{nonAFR}$ was defined as the union of segments from all non-African populations (GBR, CHB, ITU).

We then estimated a matrix of the mean $P_Y^{X(i)}$ across all individual $i$ in population $X$ for all population pairs $(X, Y)$, with the same $G_{nonAFR}$ definitions for Africans (wAFR, eAFR) applied (**fig. S23A**) while for each non-African populations, $G_{nonAFR}$ was defined as the union of segments from the other two non-African populations. For comparison, the same analysis was performed for Neanderthal ancestry segments (**fig. S23B**).

We observed that non-Africans share a great proportion of their ghost ancestry with Africans, while Africans retain higher diversity of unique ghost segments (**fig. S23**). Ghost ancestry is also more shared between West and East African populations and between pairs of non-African populations than between non-Africans and Africans. These patterns hold across comparisons of individual African and non-African populations and are not driven by any single population. Together, these results are consistent with expectations under an Out-of-Africa dispersal model, in which a ghost introgression event predates the dispersal, leading Africans to retain higher diversity of ghost ancestry while non-Africans carry a subset (largely shared among non-African populations) shaped by the Out-of-Africa bottleneck.

We calculated heterozygosity in every 50kbp window genome-wide. For each population, the "Background" heterozygosity was the mean rate across all individuals per window. For each archaic ancestry type ($s \in \{NEA, DEN, Ghost\}$), we calculated the mean heterozygosity for the subset of individuals carrying that archaic ancestry in windows overlapping archaic haplotypes. As shown in fig. S24, ghost segments show a similar elevation in heterozygosity compared to Neanderthal or Denisovan segments across all populations, unsurprising given the expectation of "long branches" underlying both introgression signals (**fig. S24,** (*55*)).

Finally, we compared the length of the ghost ancestry segments and Neanderthal ancestry segments. Across all populations, ghost ancestry segments exhibit systematically shorter haplotype lengths than Neanderthal ancestry segments (**fig. S25**). This pattern is evident in both African (wAFR, eAFR) and non-African (EUR, EAS, SAS) populations, with ghost segments showing lower mean lengths and a stronger skew toward shorter tracts. Because recombination progressively breaks down introgressed haplotypes over time, shorter segment lengths are expected for older introgression events. The consistently reduced length of ghost segments therefore supports the inference that ghost introgression predates Neanderthal introgression, which has been inferred to have occurred ~47,000 years ago (*64*).

Together, these results are consistent with a ghost introgression event predating the Out-of-Africa dispersal, followed by reduced diversity in non-Africans due to the OOA bottleneck.

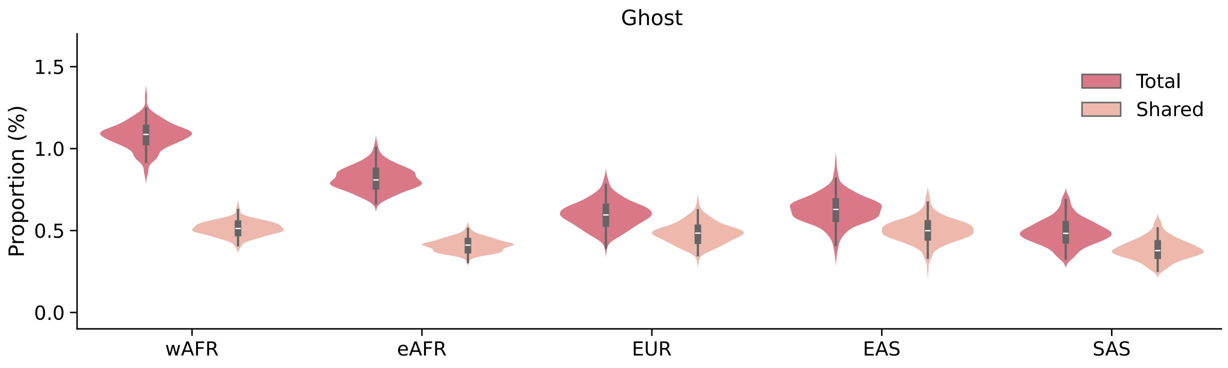

**Fig S22**. Distribution and sharing of ghost segments across populations. The proportion of ghost segments per individual is shown as the total (dark) and the subset shared with other populations (light). For African (wAFR and eAFR) individuals, sharing is calculated with any non-African population. For non-Africans (EUR, EAS, SAS), sharing is calculated with West Africans (wAFR).

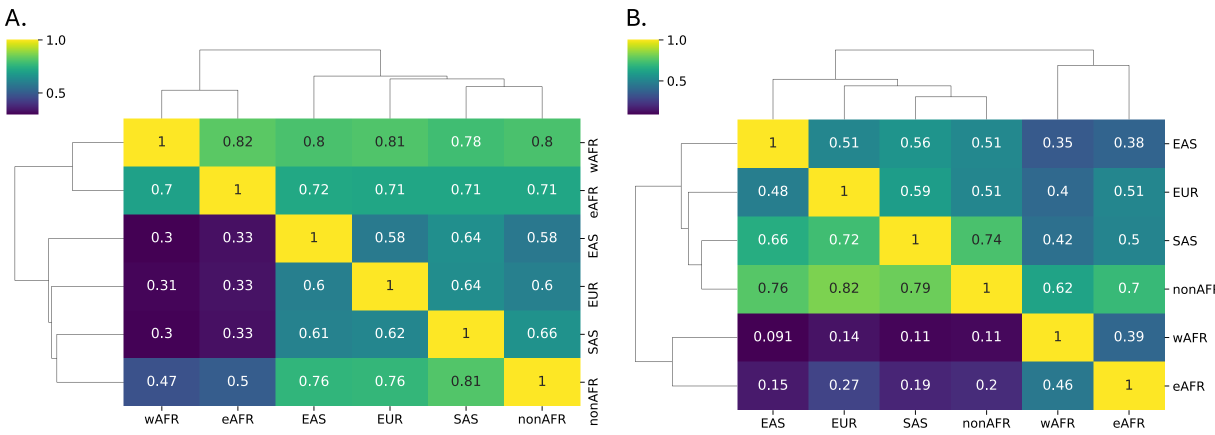

A.

| | wAFR | eAFR | EAS | EUR | SAS | nonAFR |
|---|---|---|---|---|---|---|
| wAFR | 1 | 0.82 | 0.8 | 0.81 | 0.78 | 0.8 |
| eAFR | 0.7 | 1 | 0.72 | 0.71 | 0.71 | 0.71 |
| EAS | 0.3 | 0.33 | 1 | 0.58 | 0.64 | 0.58 |
| EUR | 0.31 | 0.33 | 0.6 | 1 | 0.64 | 0.6 |
| SAS | 0.3 | 0.33 | 0.61 | 0.62 | 1 | 0.66 |
| nonAFR | 0.47 | 0.5 | 0.76 | 0.76 | 0.81 | 1 |

B.

| | EAS | EUR | SAS | nonAFR | wAFR | eAFR |
|---|---|---|---|---|---|---|
| EAS | 1 | 0.51 | 0.56 | 0.51 | 0.35 | 0.38 |
| EUR | 0.48 | 1 | 0.59 | 0.51 | 0.4 | 0.51 |
| SAS | 0.66 | 0.72 | 1 | 0.74 | 0.42 | 0.5 |
| nonAFR | 0.76 | 0.82 | 0.79 | 1 | 0.62 | 0.7 |
| wAFR | 0.091 | 0.14 | 0.11 | 0.11 | 1 | 0.39 |
| eAFR | 0.15 | 0.27 | 0.19 | 0.2 | 0.46 | 1 |

**Fig S23**. Population sharing of archaic ancestry. Heatmap shows the mean proportion of ghost ancestry **(A)** and Neanderthal ancestry **(B)** in each population (x-axis) that is shared with every other population (y-axis). Values represent the average across all individuals within a population.

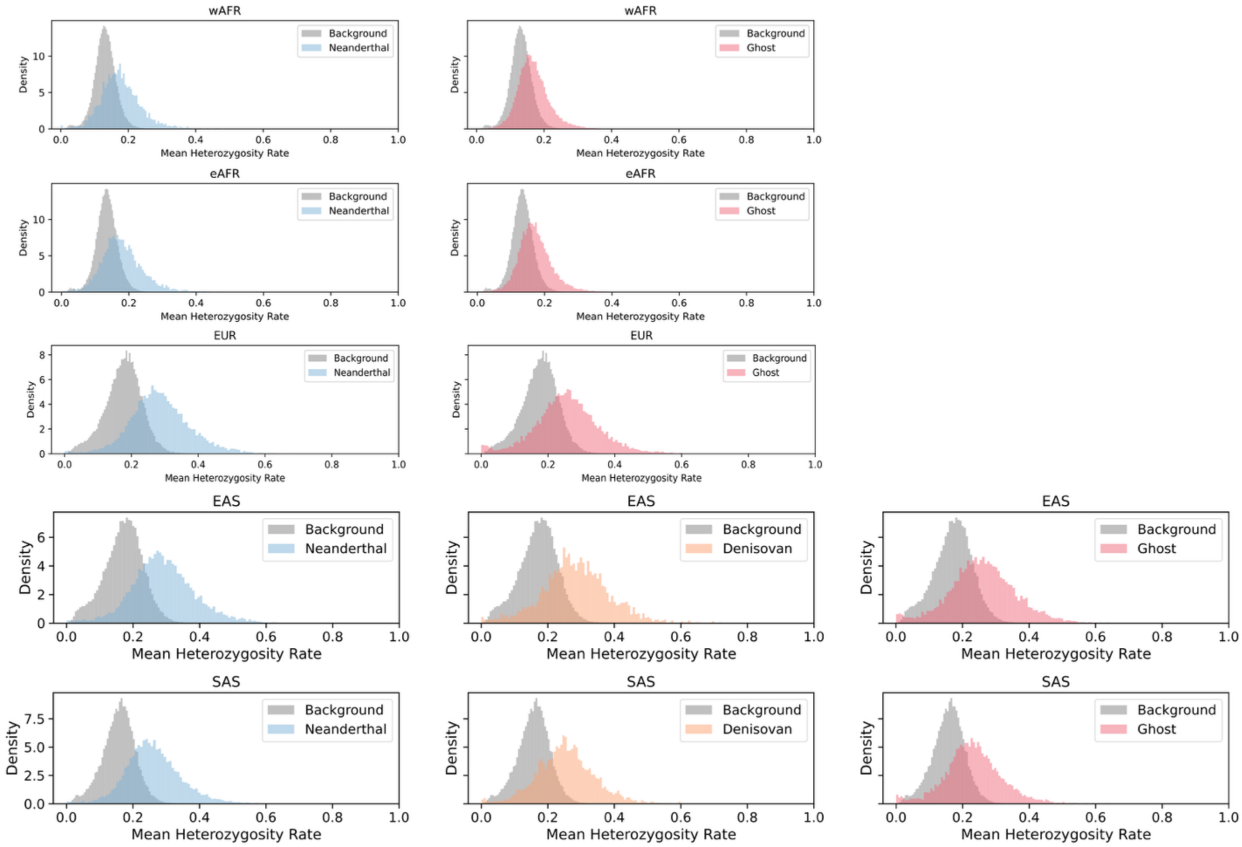

**Fig S24**. Heterozygosity in archaic segments across populations. Each row represents a population (wAFR, eAFR, EUR, EAS, SAS). The mean heterozygosity rate is shown for genomic regions containing Neanderthal (blue), Denisovan (orange), or Ghost (red) segments, compared to the genome-wide background (grey).

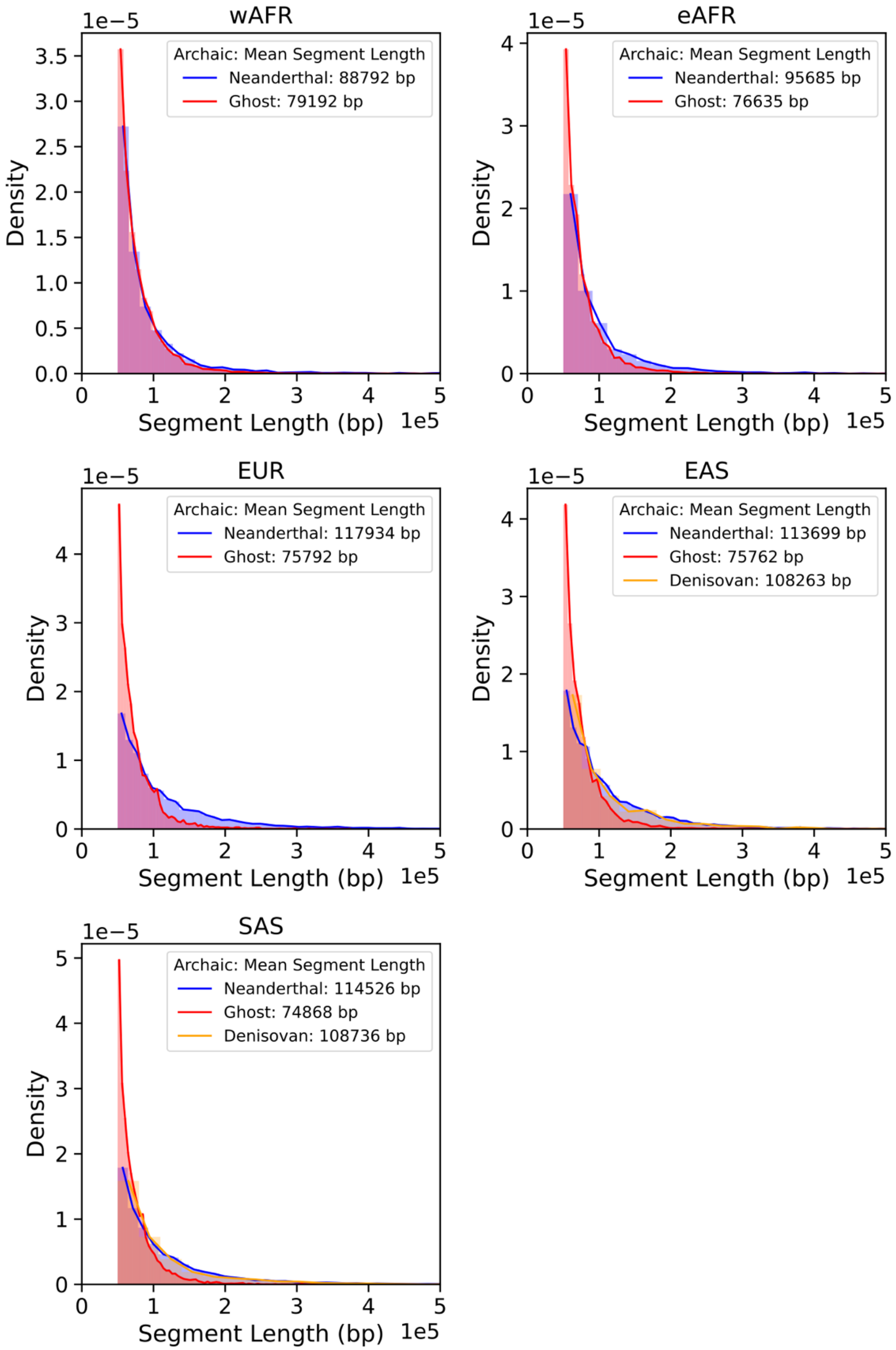

**Fig S25**. Haplotype length distributions of archaic segments. Segment length distributions (in base pairs) for Neanderthal (blue), Denisovan (yellow), and Ghost (red) ancestry across populations.

### Section S5. Validating ghost admixture using simulations

#### S5.1 Simulation models

To validate the signal of ghost admixture in simulations, we constructed an Out-of-Africa (OOA) model based on current understanding of the demographic history of non-African (Europeans) and African populations (OOA Model**, fig. S26A,** (*53*, *54*)). The model parameters are as follows:

- The ancestral modern human population originated with an effective size $Ne = 20{,}000$. Approximately 575,000 years before present, $Ne$ increased to 25,000 and remained constant until the present, leading to the present-day African population.
- The Neanderthal-Denisovan lineage diverged from modern humans 575,000 years ago, with an initial $Ne = 5{,}000$. Neanderthals and Denisovans split from each other around 420,000 years ago, each maintaining a constant $Ne = 2{,}000$ thereafter. The sequenced Neanderthal lineage (Altai) diverged from the introgressing Neanderthal source 105,000 years ago, while the sequenced Denisovan lineage diverged from the introgressing Denisovan source 300,000 years ago.
- The non-African population split from Africans 60,000 years ago and underwent an instantaneous bottleneck with $Ne = 1{,}000$, which lasted for 14,000 years. This was followed by an exponential growth to a final $Ne = 50{,}000$ by the present.
- A single pulse of 2% Neanderthal introgression into the non-African population occurred 50,000 years ago.
- A single pulse of 0.1% Denisovan introgression into the non-African population occurred 45,000 years ago.
- A single pulse of 5% back-migration from the non-African population into Africans occurred 40,000 years ago.

We next constructed an "OOA + Ghost" model (**fig. S26B**) by incorporating a ghost archaic lineage into the baseline OOA model, with parameters informed by previous empirical results (*13*):

- The ghost population diverged from the modern human lineage 783,000 years ago.
- A single pulse of 9% ghost introgression into the ancestors of all modern humans occurred 87,000 years ago (prior to the OOA split).

We generated 10 simulation replicates of 50 Mbp genomes under both the OOA and OOA + Ghost demographic models. From each simulation, we sampled 100 non-African and 100 African modern human individuals. We also included one diploid individual each from the sequenced Neanderthal and Denisovan populations (two archaic genomes in total). The sequenced Neanderthal genome is sampled 46,400 years before present in simulations, while the sequenced Denisovan genome is sampled 43,500 years before present. The VCF output containing all genotype information for the 200 modern humans from each simulation served as the input for *SINGER*.

We ran *SINGER* with the parameters `-m 1.2e-8 -n 300 -thin 100 -Ne 2e4 -polar 0.99` on all simulations and used the 250$^{th}$ to 299$^{th}$ posterior tree sequences for all downstream analyses. We extracted the observation data for TRACE from each marginal tree and summarized it into 1000 bp windows (as described in **Section S3.1**). The final input for TRACE was generated by averaging this summarized data across all 50 posterior tree sequences. We applied TRACE with

$t = 15000$ generations to the *SINGER*-inferred trees. The output was filtered to retain segments with posterior probability > 0.9 and length > 0.05cM, 50kbp. Finally, we classified the filtered segments into Neanderthal, Denisovan, and Ghost categories using *Rule Group 1* (**Section S4.2**).

We ran *hmmix* using all 100 sampled African individuals as the outgroup. To ensure robust performance evaluation and mitigate potential bias from training on small genomic datasets, we adopted the following procedure:

- Training: For each individual, we merged genotypes across 10 simulation replicates (creating a combined 500 Mbp genome) and trained *hmmix* on this merged VCF.
- Decoding: We then used the trained parameters on each individual to decode that same individual in each simulated replicate separately.

The *hmmix* output was filtered to retain segments with posterior probability > 0.8 and length > 0.05cM, 50kbp. To classify these segments, we used the sampled archaic genomes from simulations. A segment was labeled "Neanderthal" if it shared more derived alleles with the sequenced Neanderthal genome than with the sequenced Denisovan genome, and "Denisovan" in the converse case. Segments with an equal number of shared derived alleles were labeled as "Ambiguous", and segments with no shared derived alleles with either archaic genomes are removed (*53*).

We ran *IBDmix* using the sampled Neanderthal and Denisovan genomes as the archaic reference panels. The *IBDmix* output was filtered for `slod` > 4 and length > 0.05cM, 50kbp. Segments passing these filters that were identified using the Neanderthal reference were labeled "Neanderthal," while those identified using the Denisovan reference were labeled "Denisovan."

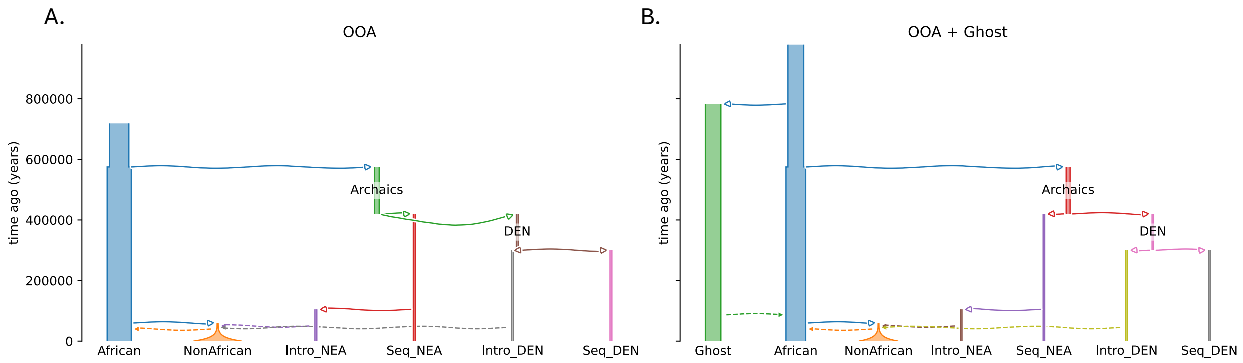

**Fig S26**. Demographic models for ghost admixture validation. **(A)** OOA model: a model recapitulating known demographic history for 1000 Genoms YRI (African) and GBR (NonAfrican). **(B)** OOA + Ghost model: OOA model incorporating a ghost lineage, parameters following (*13*).

#### S5.2 Performance of inference methods in simulations

We assessed the precision and recall of each method using the approach detailed in Section S1.3. The results demonstrate that the OOA model cannot explain the substantial number of non-Neanderthal segments we detected in 1000 Genomes GBR individuals (**table S7**). Specifically, in the absence of ghost introgression (OOA model), TRACE identifies less than 0.1% of the genome per individual as "Ghost" and recovers less total archaic ancestry than *hmmix*—a pattern inconsistent with real data. In contrast, with additional ghost ancestry TRACE recovered in the OOA + Ghost model, both the total archaic and Neanderthal segment quantities better match empirical observations (**fig. S15**).

We further evaluated method concordance by calculating the proportion of TRACE segments overlapping those from *hmmix* and *IBDmix* (**Section S4.2**). As shown in table S8, the OOA model fails to recapitulate the key pattern observed in real data: low overall overlap of archaic segments between TRACE and *hmmix* (or *IBDmix*) but high overlap for Neanderthal segments specifically. The OOA + Ghost model, however, reproduces this pattern accurately, providing further support that this model captures patterns observed in real data (**table S4**).

#### *S5.3 Archaic affinity and site frequency spectra*

We assessed the Neanderthal and Denisovan affinity of segments detected by TRACE using the sampled archaic genomes from simulations. Ghost segments inferred from simulations under the OOA + Ghost model exhibit the expected pattern of nearly identical affinity to both sequenced Neanderthal and Denisovan genomes (**fig. S27**), mirroring the signature observed in the real data (**fig. S18**).

We also reconstructed the Site Frequency Spectrum (SFS) and conditional SFS (cSFS) for the detected segments in both African and non-African populations (**Section S4.4**). The results (**fig. S28-S29**) show broadly consistent patterns between simulated and real data. One notable discrepancy is that the cSFS for Neanderthal segments in the real data is more skewed toward low-frequency alleles than in simulations. However, the relative difference in Neanderthal cSFS between African and non-African populations is consistent, e.g. non-Africans have more dispersed low-frequency-bin elevation patterns not found in Africans. We hypothesize the discrepancy may stem from a less severe Out-of-Africa bottleneck in real data than modeled in our simulations, or it may reflect the impact of negative selection against Neanderthal ancestry (*52*, *57*, *58*) that was not incorporated in these simulations.

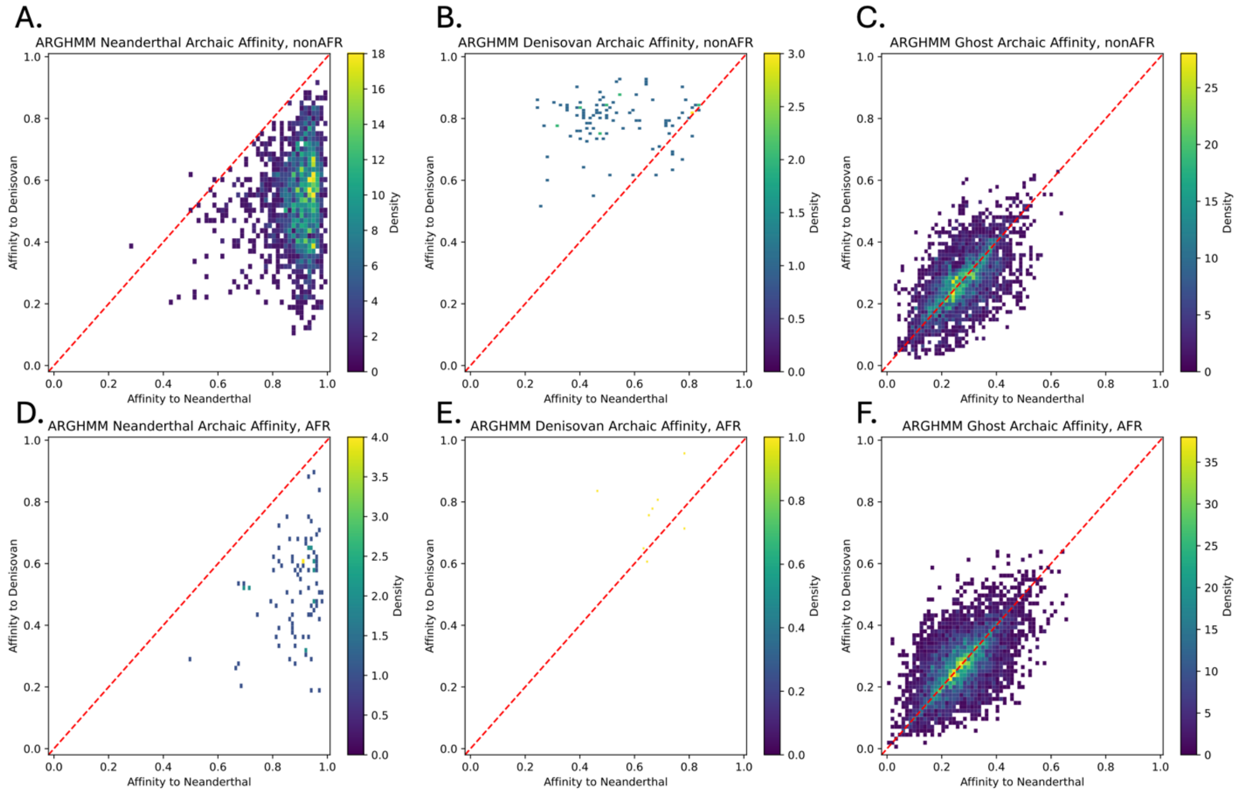

**Fig S27**. Neanderthal versus Denisovan genetic affinity for TRACE inferred archaic segments under OOA + Ghost model. TRACE-detected segments are classified as Neanderthal (left column), Denisovan (middle), and Ghost (right) ancestry. Rows represent populations: NonAfricans (row 1), Africans (row 2).

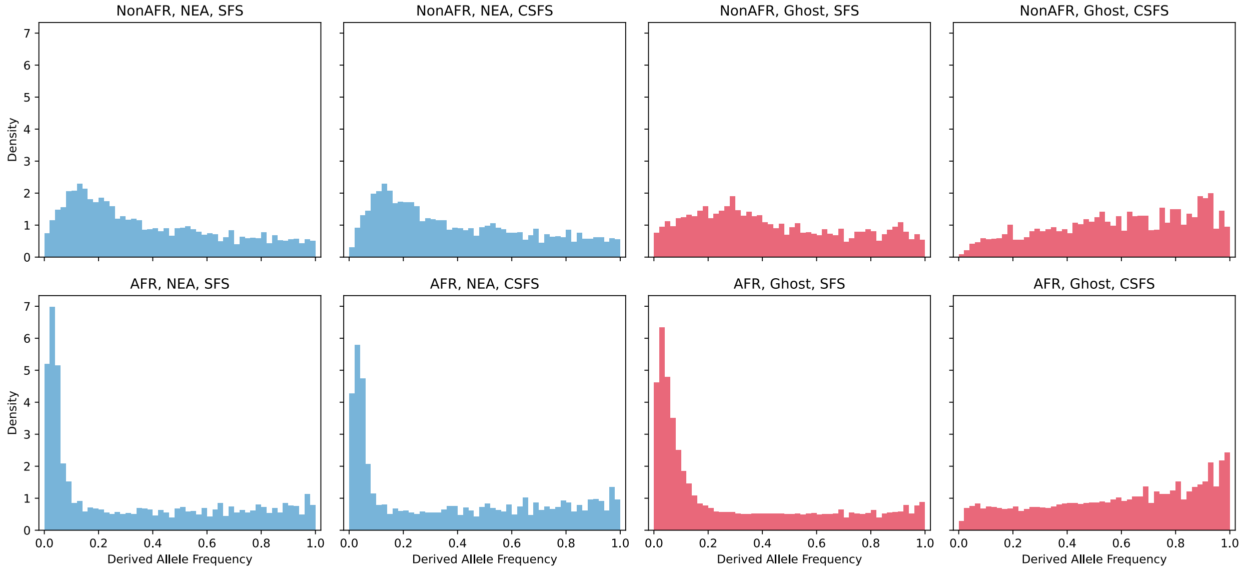

**Fig S28**. (conditional on Neanderthal) Site frequency spectra of TRACE inferred archaic segments under OOA + Ghost model. Rows represent populations: NonAfricans, Africans. For each population, the site frequency spectrum (SFS, left column) and conditional SFS (cSFS, right column) are shown for alleles on TRACE-detected segments. Colors indicate the assigned segments archaic ancestry: Neanderthal (blue) and Ghost (red). The cSFS is conditioned on the derived allele being present in the sampled Neanderthal genome.

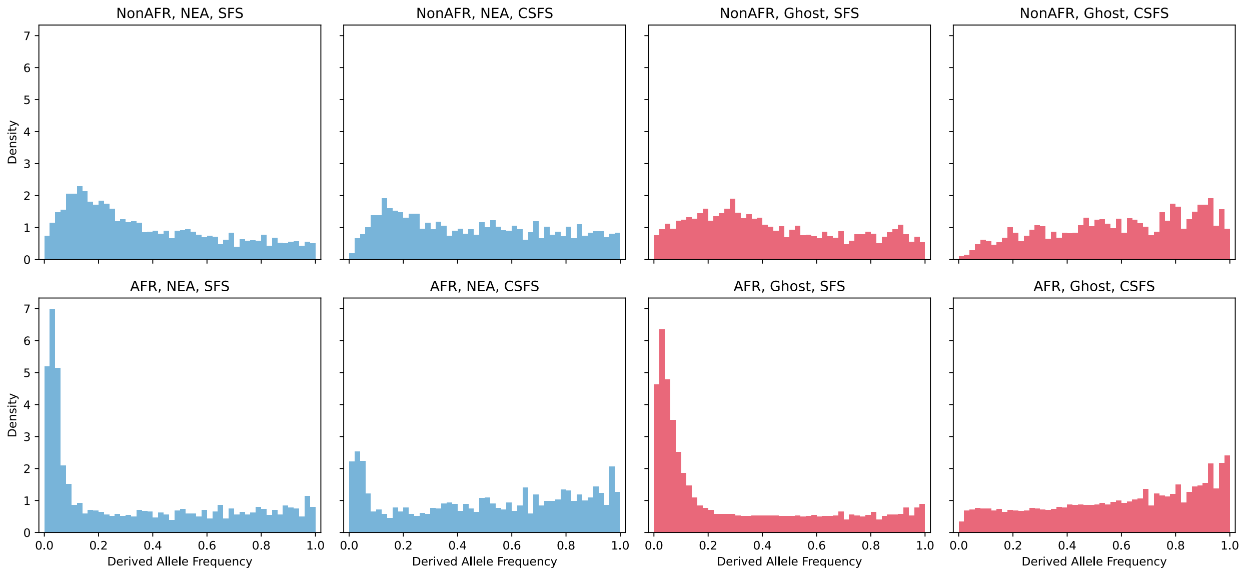

**Fig S29**. (conditional on Denisovan) Site frequency spectra of TRACE inferred archaic segments under OOA + Ghost model. Rows represent populations: NonAfricans, Africans. For each population, the site frequency spectrum (SFS, left column) and conditional SFS (cSFS, right column) are shown for alleles on TRACE-detected segments. Colors indicate the assigned segments archaic ancestry: Neanderthal (blue) and Ghost (red). The cSFS is conditioned on the derived allele being present in the sampled Denisovan genome.

### Section S6. Genomic landscape of ghost admixture

#### S6.1 Correlation between ghost ancestry, background selection, and recombination rate

We investigated the relationship between archaic ancestry and regions under linked selection (using the B statistic (*59*)) as well as local recombination rates (HapMap). For each 10kbp window genome-wide (290,000 windows in total), we calculated the mean B-score and recombination rate, along with the population frequency of TRACE-inferred archaic segments (Neanderthal, Denisovan, Ghost).

Consistent with previous findings (*32*, *52*), both Neanderthal and Denisovan ancestries are significantly depleted ($p < 2.2 \times 10^{-308}$) in regions of low B-score and low recombination across all populations (**table S9**). Ghost ancestry displays a nearly identical pattern, showing significant positive correlations with both B-score and recombination rate, indicating similar depletion in conserved, low-recombination regions.

To visualize these relationships, we binned the genome by recombination rate and B-score into six quantile-based intervals each using `pandas qcut` function:

- (-0.001, 2.72e-08], (2.72e-08, 0.00342], (0.00342, 0.0719], (0.0719, 0.545], (0.545, 4.344], (4.344, +∞) for recombination rate bin 0-5.

- (-0.001, 669.0], (669.0, 795.0], (795.0, 863.0], (863.0, 907.0], (907.0, 943.0], (943.0, +∞) for B-score bin 0-5.

Figures S30-S31 show a clear positive correlation between these binned values and mean archaic ancestry frequency. Error bars, derived from block jackknife resampling ($n = 290$) with block size of 10 Mbp, confirm the robustness of this trend. This result holds whether archaic ancestry segments are analyzed across all populations jointly (**fig. S30**) or examined separately within each population (**fig. S31**).

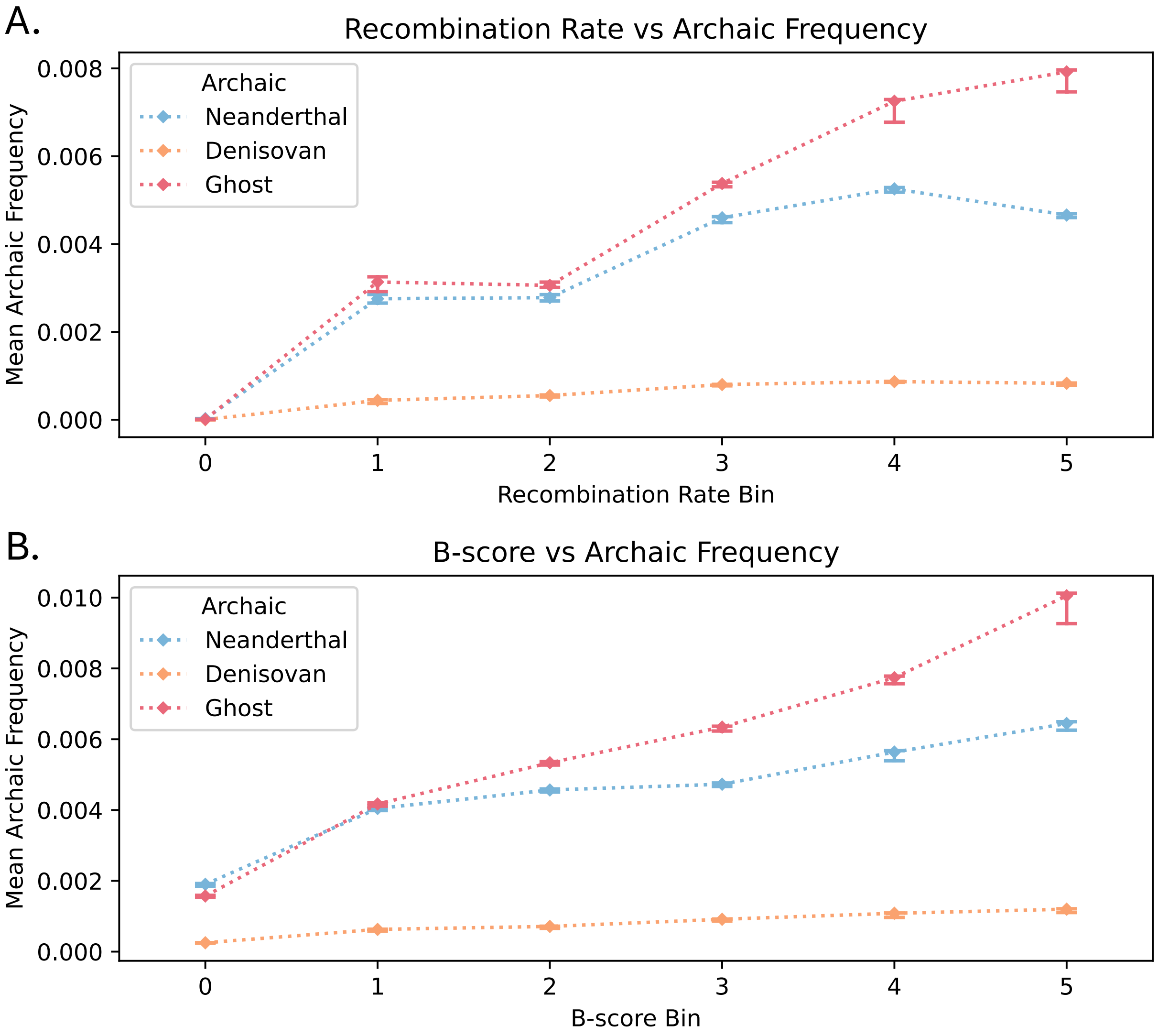

**Fig S30.** Correlation of archaic ancestry frequency with genomic features. The mean frequency of total archaic ancestry (across wAFR, eAFR, EUR, EAS and SAS) is plotted against binned values of **(A)** recombination rate and **(B)** B-score in 10kbp windows. Error bars show standard errors across 290 jackknife resampling replicates. See Supplementary Section S6.1 for bin definition.

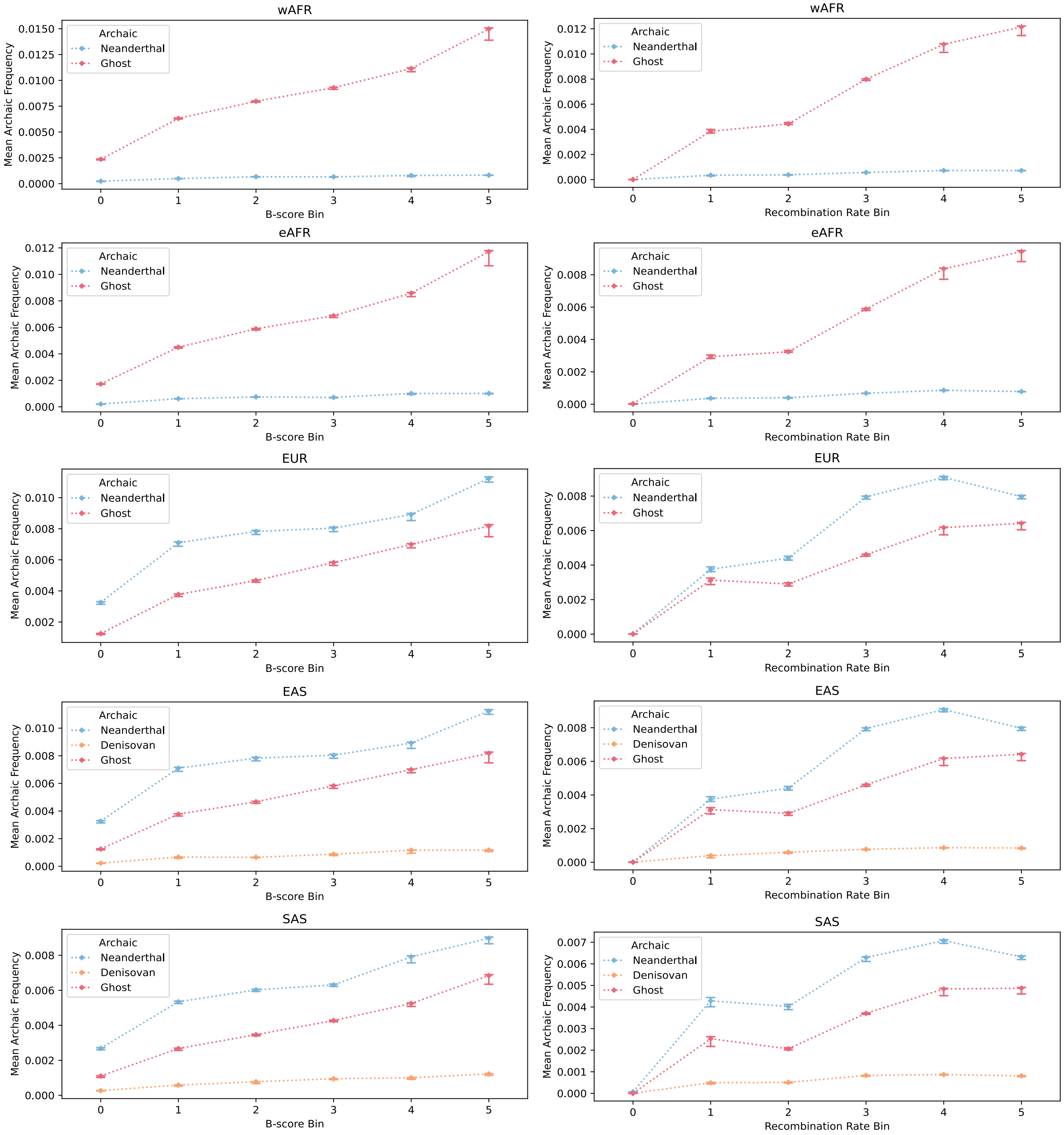

**Fig S31.** Correlation of archaic ancestry frequency with genomic features. Mean archaic ancestry frequency in 10kbp windows is plotted against binned recombination rate (left) and B-score (right) for populations wAFR, eAFR, EUR, EAS, and SAS (rows 1-5). Error bars indicate standard error from 290 jackknife replicates. Bin definitions are provided in Supplementary Section S6.1.

#### S6.2 Regions with enriched ghost ancestry

To identify regions enriched for ghost ancestry, we use a criterion defined previously for quantifying heterogeneity in archaic ancestry (*53*). We defined peaks or enriched regions for ghost ancestry as non-overlapping one kbp windows of the genome where the frequency of ghost ancestry (estimated based on the number of ghost haplotypes overlapping the region within a sample) is greater than two standard deviations above the genome-wide mean, after removing any windows that do not contain assigned ancestry across samples. We applied this enrichment criteria separately for each population to identify regions exhibiting population-specific enrichment. We detect 1932 regions enriched for ghost ancestry, with substantial heterogeneity across populations: 825 wAFR, 550 eAFR, 176 EUR, 188 SAS and 193 EAS (**Table S12; fig. S32-S33**).

To investigate the functional impact of ghost-enriched regions, we used the tool GREAT to test for enrichment of annotations and molecular pathways (**fig. S34**) (*87*). Gene assignments for peaks were performed using `bedtools intersect`, based on protein-coding gene annotations defined in Gencode v49 (**table S1**, (*88*)). We find that several genes intersect with peaks of ghost ancestry, such as *CSMD1* and *RBFOX1* (**fig. S35**). Although this metric is inherently biased towards larger genes, these intersections are supported across multiple populations, indicating a stronger and robust signal of enrichment of ghost ancestry at these loci.

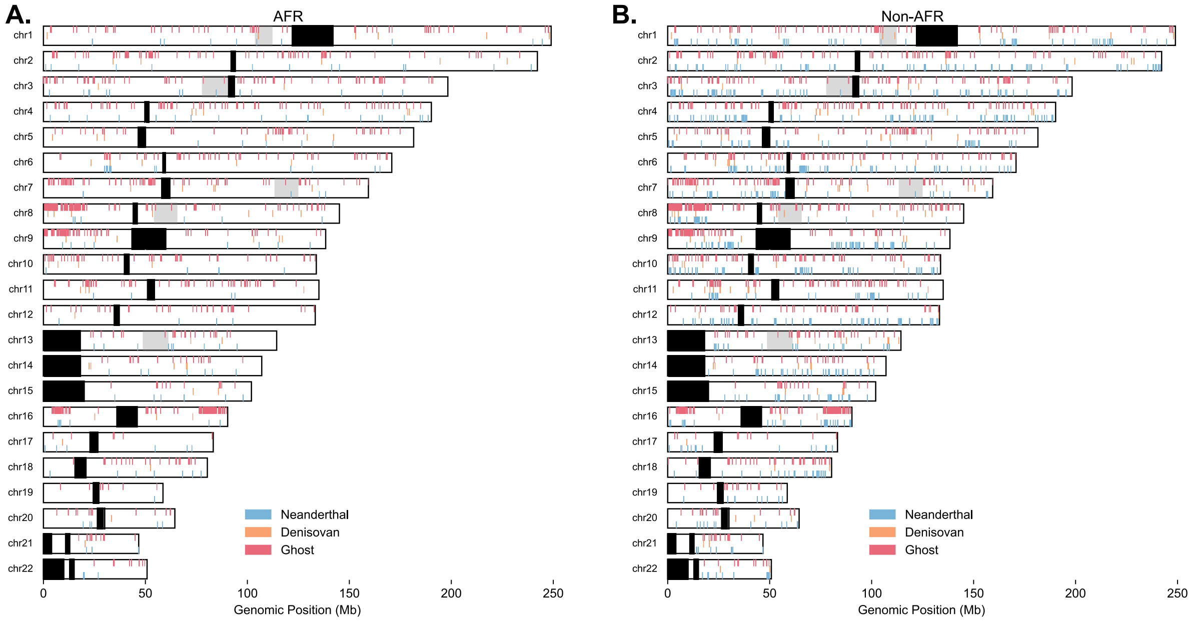

**Fig S32**. Estimated peaks of archaic ancestry, including putative Ghost ancestry, stratified by **(A)** within African (wAFR + eAFR) or **(B)** Non-African populations (EUR + SAS + EAS) across the genome. Peaks are shown across all archaic ancestries estimated in TRACE (with posterior probability > 0.9, length > 50kbp, 0.05cM). Black boxes indicate genomic regions masked as centromeric or regions without variants which are excluded from ARG inference.

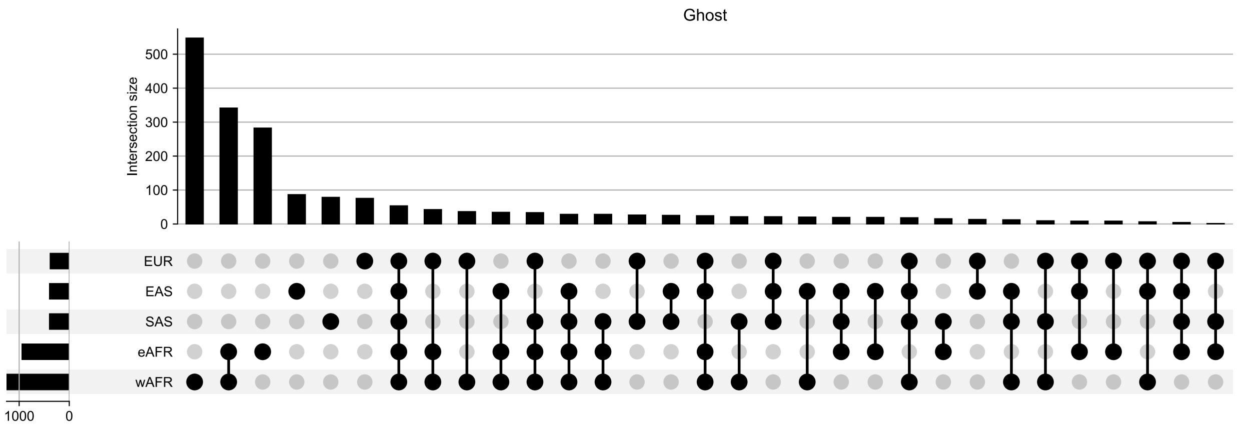

**Fig S33**. Sharing of ghost ancestry peaks across populations. Sharing between populations is defined using a minimum of 100bp overlap between called peaks.

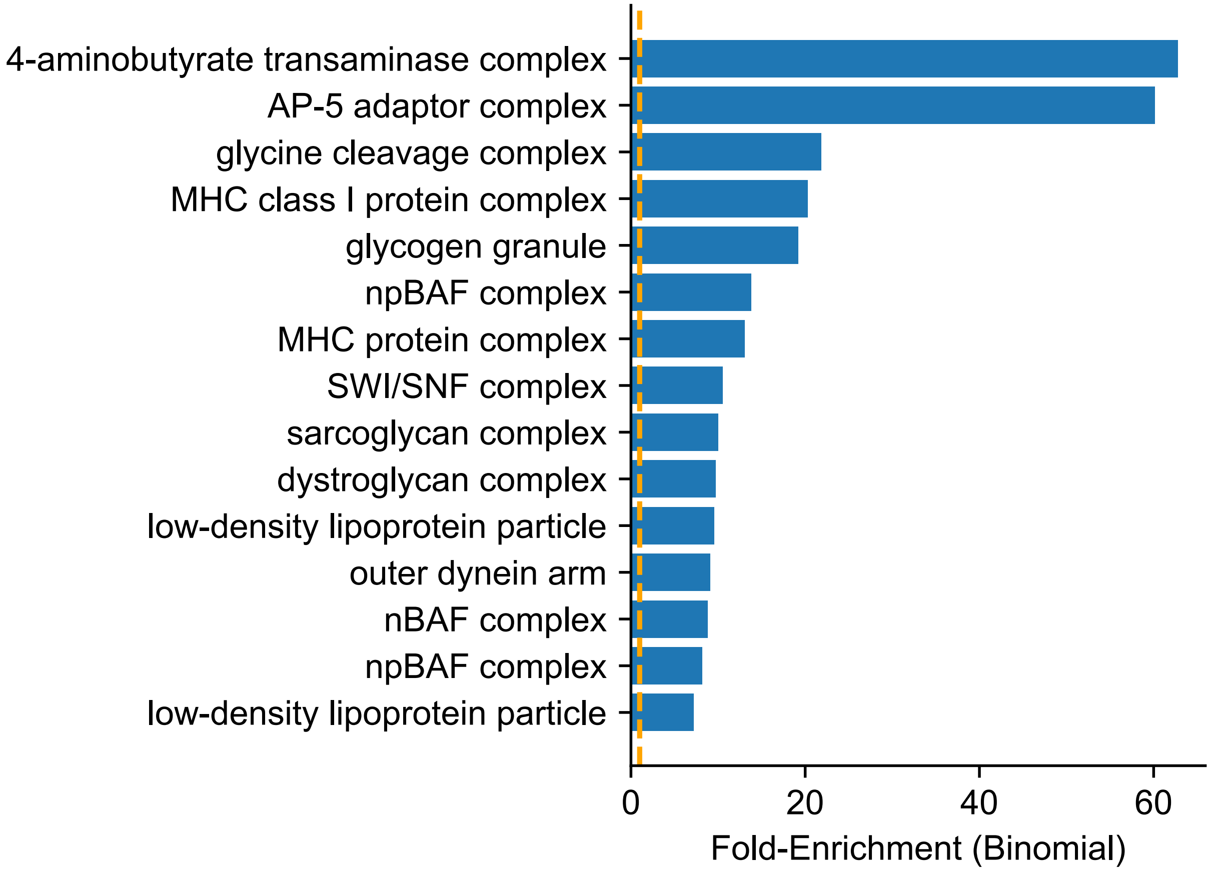

**Fig S34.** Enrichment of regional enrichments for ghost ancestry peaks estimated in population-specific analyses using GREAT. Only the top 15 categories are shown here for ease of visualization. All fold-changes for significant regions are shown for categories with $p < 0.05$ following Benjamini-Hochberg correction.

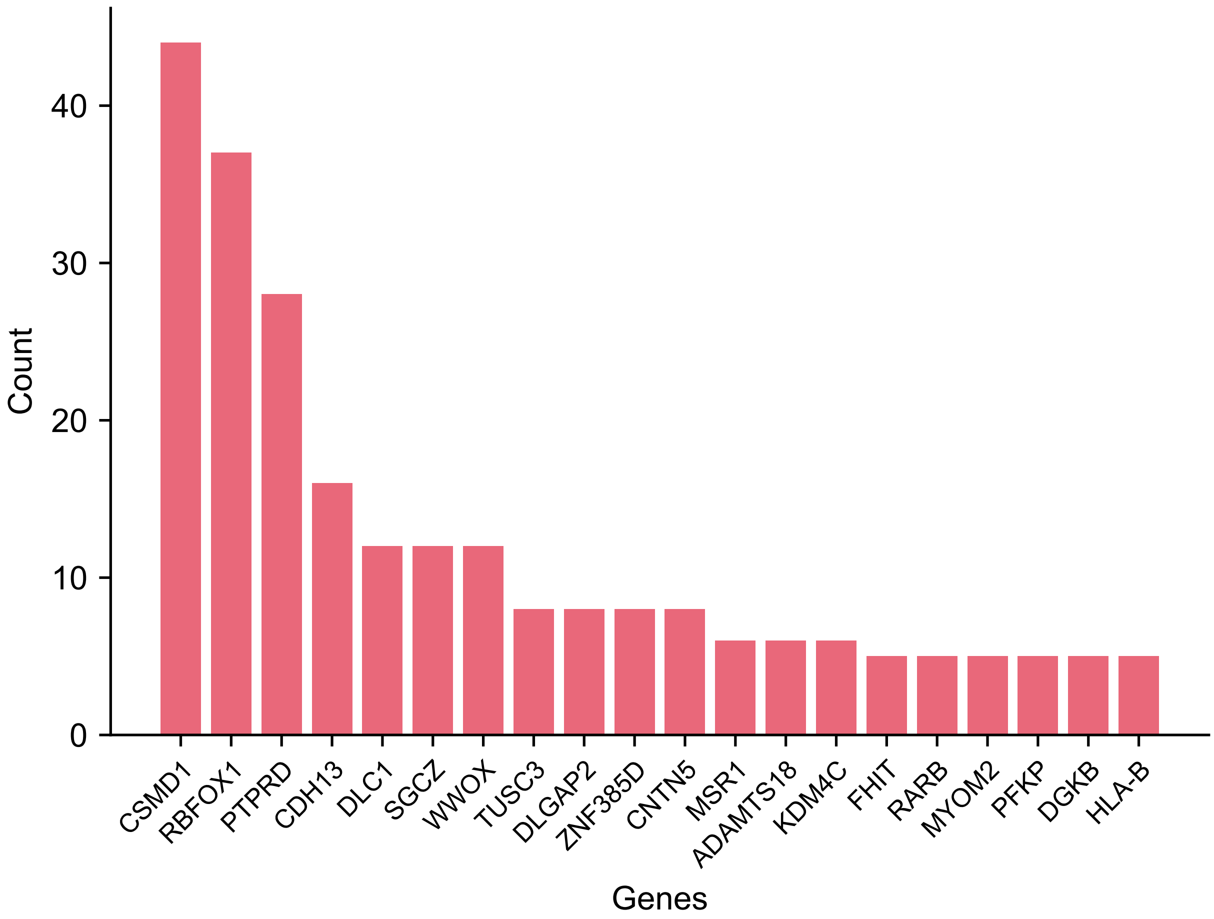

**Fig S35.** Genes intersections in high-frequency peaks of ghost ancestry. Only the top twenty genes are shown for ease of visualization. Genes are ranked by the number of ghost ancestry peaks identified in each population specifically from TRACE.

#### S6.3 Detecting deserts of ghost ancestry

Similar to previous literature defining deserts of archaic ancestry, we use two key criteria: 1) the region must be at least 10 Mbp in length and 2) have a frequency of ghost ancestry < 0.1% (*52*, *53*). The frequency is calculated as the total length of detected ghost sequence within the window normalized by the length of the window (10 Mbp) and multiplied by the number of haplotypes used for TRACE (the total haplotype length considered in the window). Given the limited recall of our method, the threshold for calling a region as a "desert" may be overly permissive such that deserts may be extended in length.

To ensure that our detection of archaic deserts is not biased by regions where ARG inference is unreliable, we considered only regions outside of the defined hg38 centromeric regions that have > 80% overlap with the strict mask from the 1000 Genomes Project (*89*). We identified deserts for each population separately. We also applied this criteria to identify Neanderthal and Denisovan ancestry deserts in non-Africans, excluding African populations from this analysis (given they have negligible amount of Neanderthal or Denisovan ancestry (*28*)).

For Neanderthal-origin deserts, we identified regions that are shared across all non-African populations (EUR, EAS, SAS), highlighting loci that consistently lack Neanderthal ancestry despite its widespread presence elsewhere in the genome. Most of these shared deserts overlap regions reported in previous studies, providing support for the robustness of our analysis (**table S11**). We did not attempt to identify deserts of Denisovan ancestry due to the overall low proportion for Denisovan ancestry across non-African populations analyzed in 1000 Genomes Project, which limits power for reliable desert detection.

We applied this framework to identify deserts of ghost ancestry per regional group in the 1000 Genomes project, and also pooling all inferred ghost segments across populations. We identified 168 ghost ancestry deserts using these criteria (**fig. S36)**. Notably, all detected ghost deserts were confined to populations outside Africa, consistent with the reduced overall levels and diversity of ghost ancestry in non-Africans (**fig. S37**). This pattern suggests that, following the Out-of-Africa bottleneck, certain genomic regions in non-African populations may have been systematically depleted of ghost ancestry, potentially due to purifying selection, drift, or a combination of both, while comparable deserts are not detectable in African populations where ghost ancestry is more abundant and diverse.

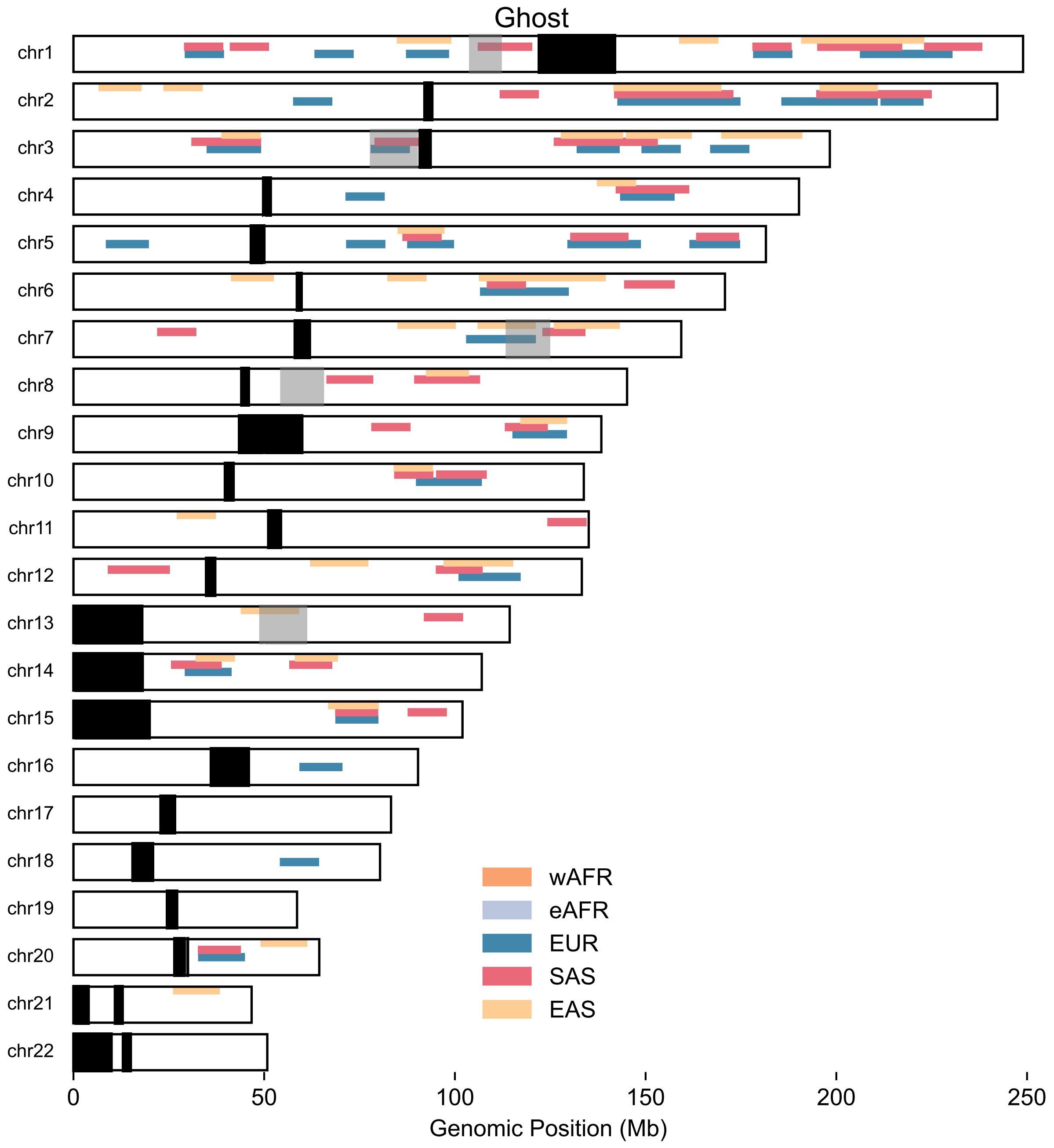

**Fig S36.** Sharing of ghost ancestry deserts across populations. Black boxes indicate genomic regions masked as centromeric or excluded from ARG inference.

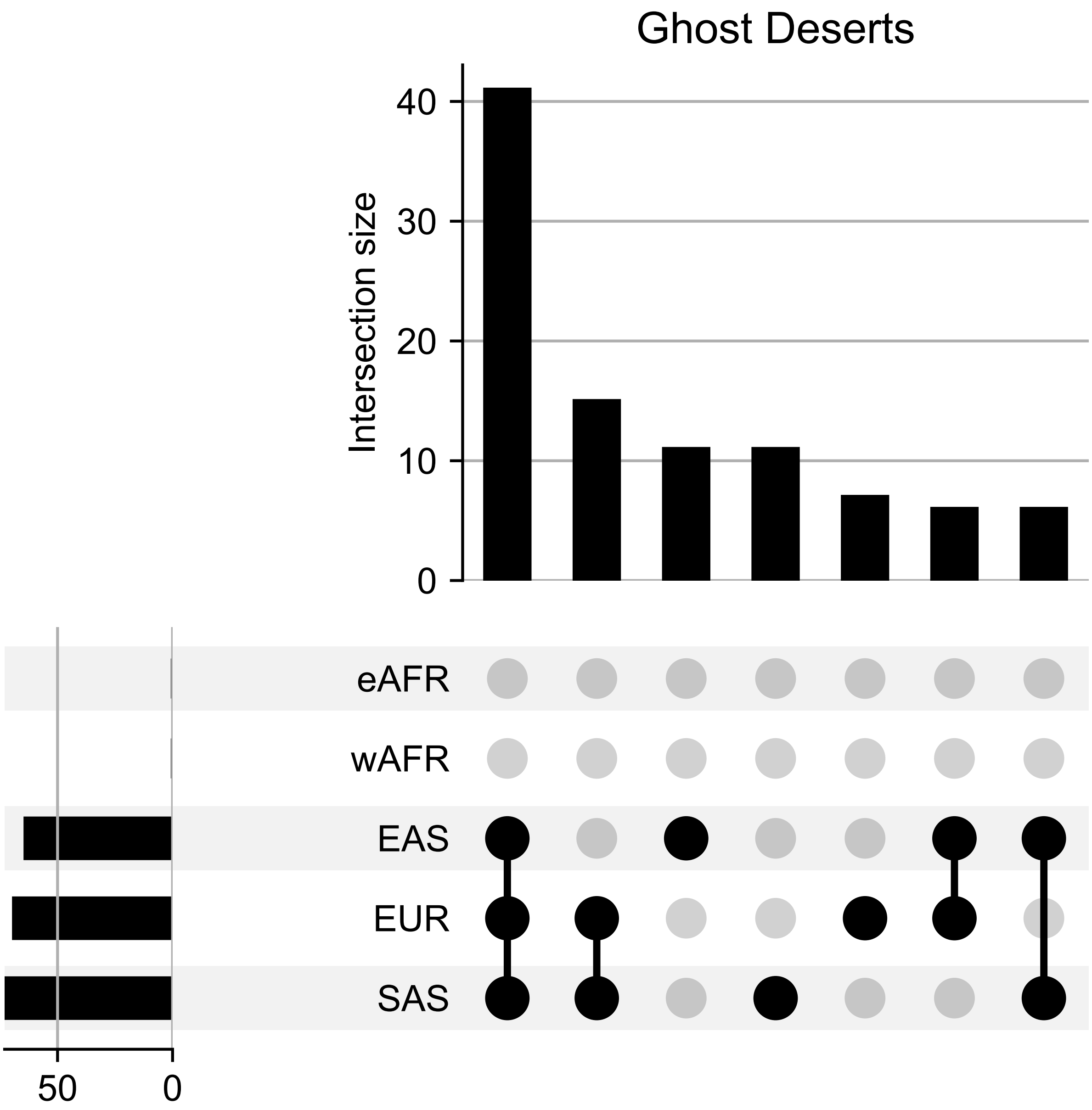

**Figure S37**. Intersection between population-specific and shared deserts of ghost ancestry. Sharing between populations is defined using a minimum of one Mbp overlap between deserts.

#### S6.4 Ghost ancestry in known Neanderthal and Denisovan deserts

Previous studies have identified five large genomic regions (> 10Mbp) as harboring limited or no Neanderthal and Denisovan ancestry ("archaic deserts"). Three out of five of these deserts have been replicated across studies (*52*, *63*). We examined whether these regions are also depleted of ghost ancestry.

To ensure high confidence in our analysis, we applied stringent filters to minimize misclassification error. We used only genomic sites that fall within the intersection of the manifesto maps for four high-coverage archaic genomes (Vindija Neanderthal, Altai Neanderthal, Chagyrskaya Neanderthal and Altai Denisovan), which spans 1,402,898,253 bp of the autosomal genome (**table S1,** (*53*)). We applied the *Rule Group 1* (**Section S4.3**) to classify inferred archaic segments longer than 50kbp and 0.05cM by their source (“Neanderthal”, “Denisovan” and “Ghost”). We analyzed both the population frequency of archaic ancestry (**Fig. 3, fig. S38-S39, table S10**) and the inferred archaic haplotypes (**fig. S40-S44**) within these five desert regions.

We first examined the frequency of inferred archaic ancestry segments in the three overlapping desert regions defined by both studies (*52*, *63*). Using previous definitions (Neanderthal deserts have < 0.1% Neanderthal frequency; Denisovan deserts have < 0.01% Denisovan frequency; (*53*)), we replicated two out of the three shared deserts, confirming them as robust Neanderthal and Denisovan deserts: chr3: 78,000,000-90,000,000 and chr7: 113,000,000-124,000,000 (**fig. S38BC, S39BC, S6.12-S6.13**). In contrast, we find non-negligible levels of Neanderthal or Denisovan ancestry in the other three deserts on chr 1 (also seen in (*53*, *64*)), chr 8 (also seen in (*53*)) and chr 13 (**fig. S38-39)**.

Interestingly, we discovered significant evidence of ghost ancestry in all of five regions (**table S10, fig. S38-S44**), and this pattern persists even when restricting to ghost segments longer than 100kbp (**fig. S39**). We infer the frequency of ghost ancestry ranges between 0–70% in these regions. The population-specific frequency patterns are consistent with expectations under a pre-OOA ghost introgression model, in which ghost segments in non-African populations are either lost or drifted to high frequencies, whereas African populations harbor lower frequency of ghost segments that are more sparsely distributed across the genome.

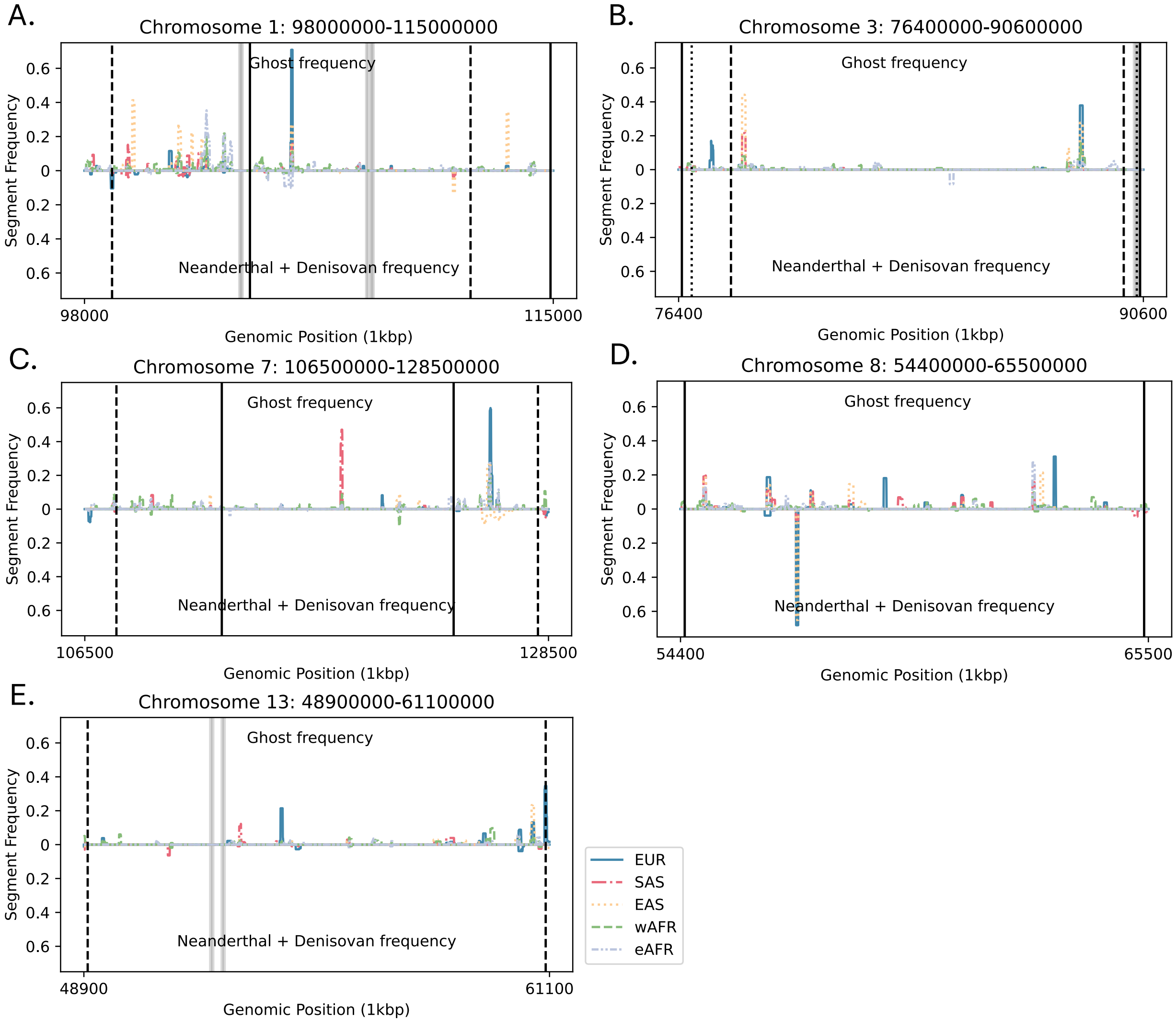

**Fig S38.** Archaic haplotype frequency (> 50kbp) in reported Neanderthal and Denisovan deserts. Population-specific frequencies of TRACE-detected segments are shown for five genomic regions previously identified as archaic ancestry deserts. Vertical lines denote desert boundaries from Vernot et al. 2016 (solid, (*63*)), Sankararaman et al. 2016 (dashed, (*52*)), and Kerdoncuff et al. 2025 (dotted, (*53*)). Grey shading indicates 50kbp windows with < 50% overlap with the 1000 Genomes strict accessibility mask.

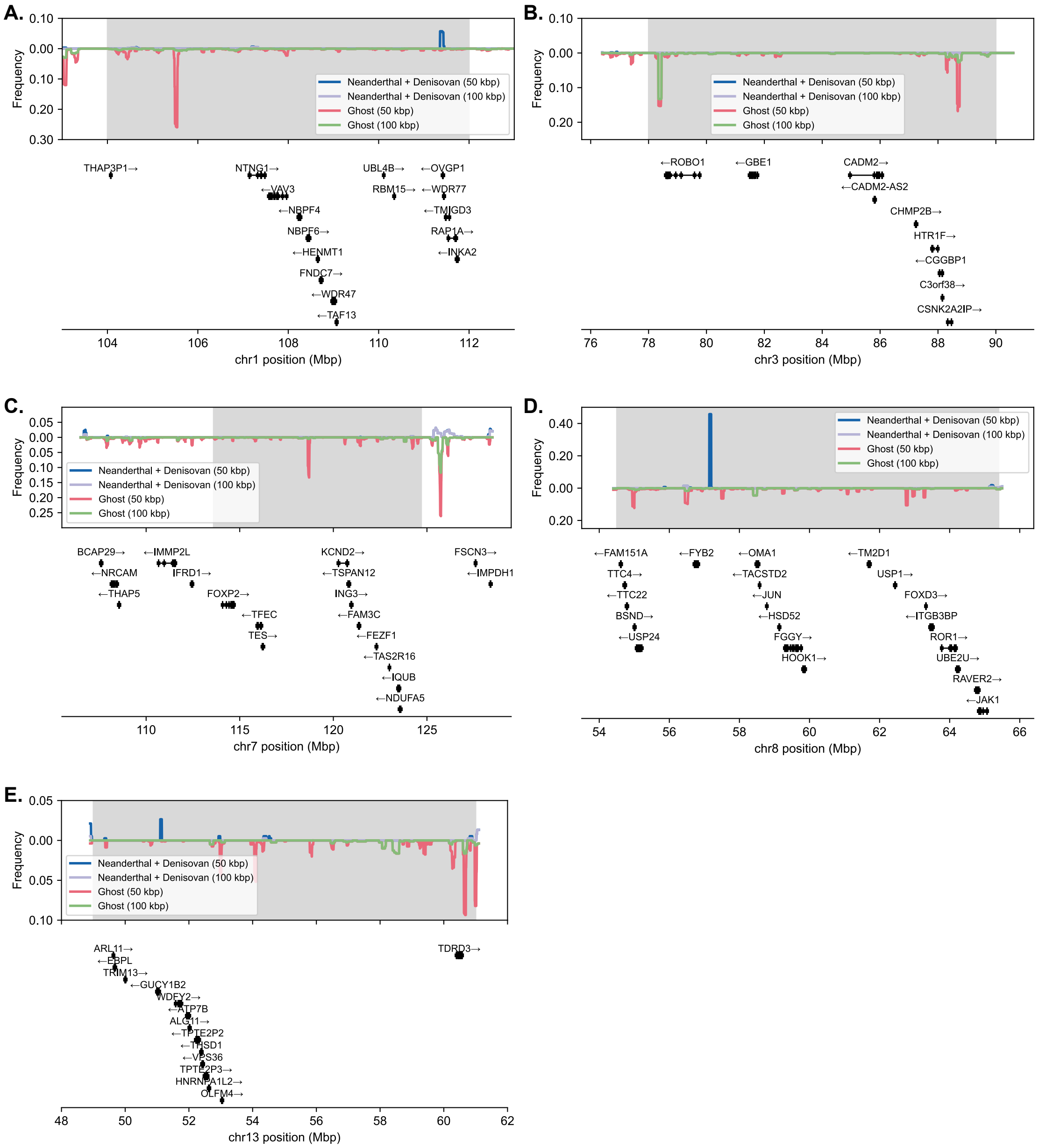

**Fig S39.** Archaic haplotype frequency in reported Neanderthal and Denisovan deserts. Frequencies of TRACE-detected segments (including ghost ancestry) are shown for five genomic regions previously identified as archaic ancestry deserts. Gray shaded boxes denote desert boundaries as an intersection from previous studies (*52*, *53*, *63*).

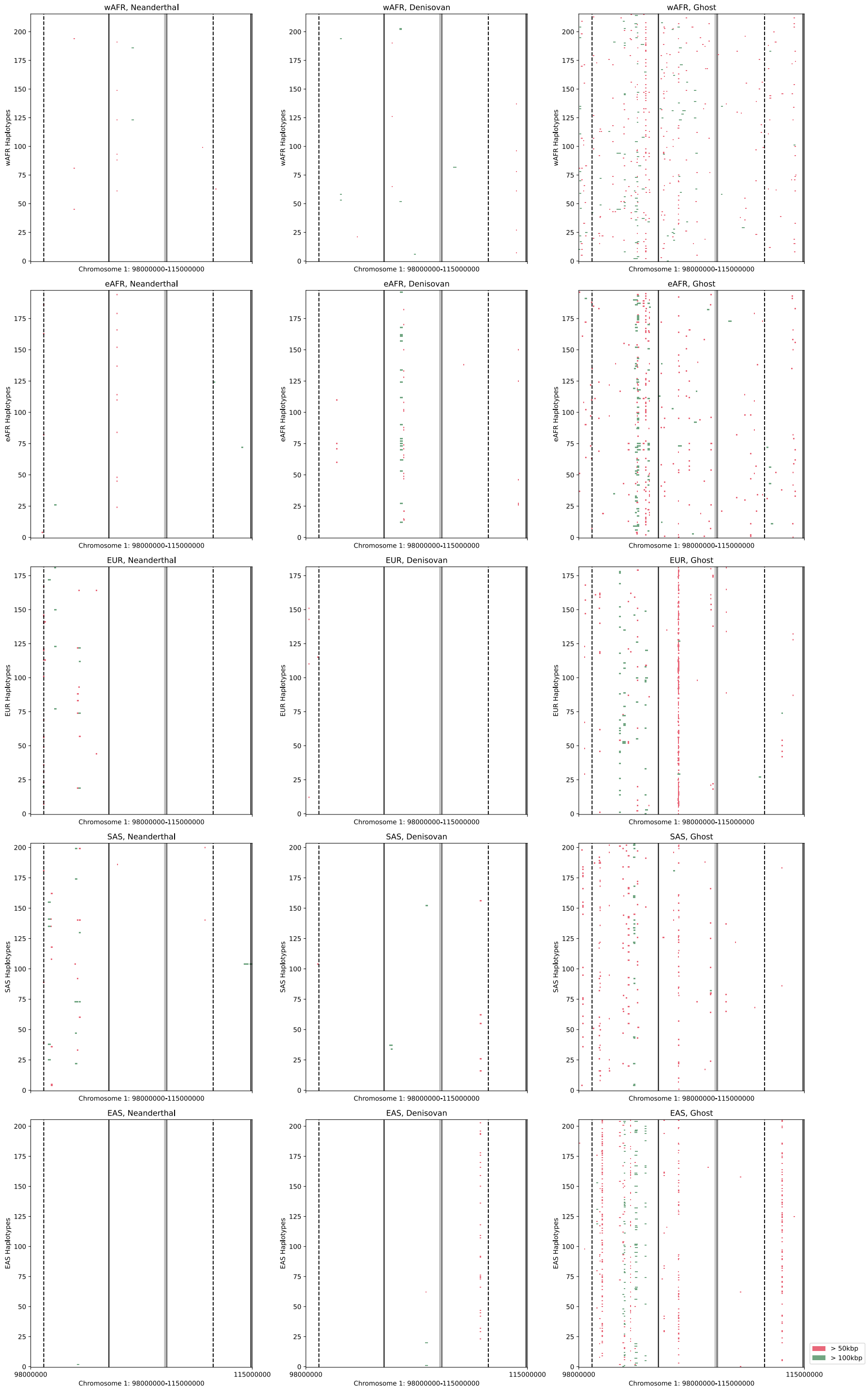

**Fig S40.** Archaic haplotypes in a desert region (chr1:98-115 Mbp). TRACE-inferred segments for different archaic ancestries are colored by length (green: > 100kbp, red: > 50kbp). Desert boundaries from Vernot et al. 2016 (solid, (*63*)), Sankararaman et al. 2016 (dashed, (*52*)), and Kerdoncuff et al. 2025 (dotted, (*53*)) are shown. Grey shading indicates low-confidence regions (< 50% overlap with the 1000 Genomes strict mask).

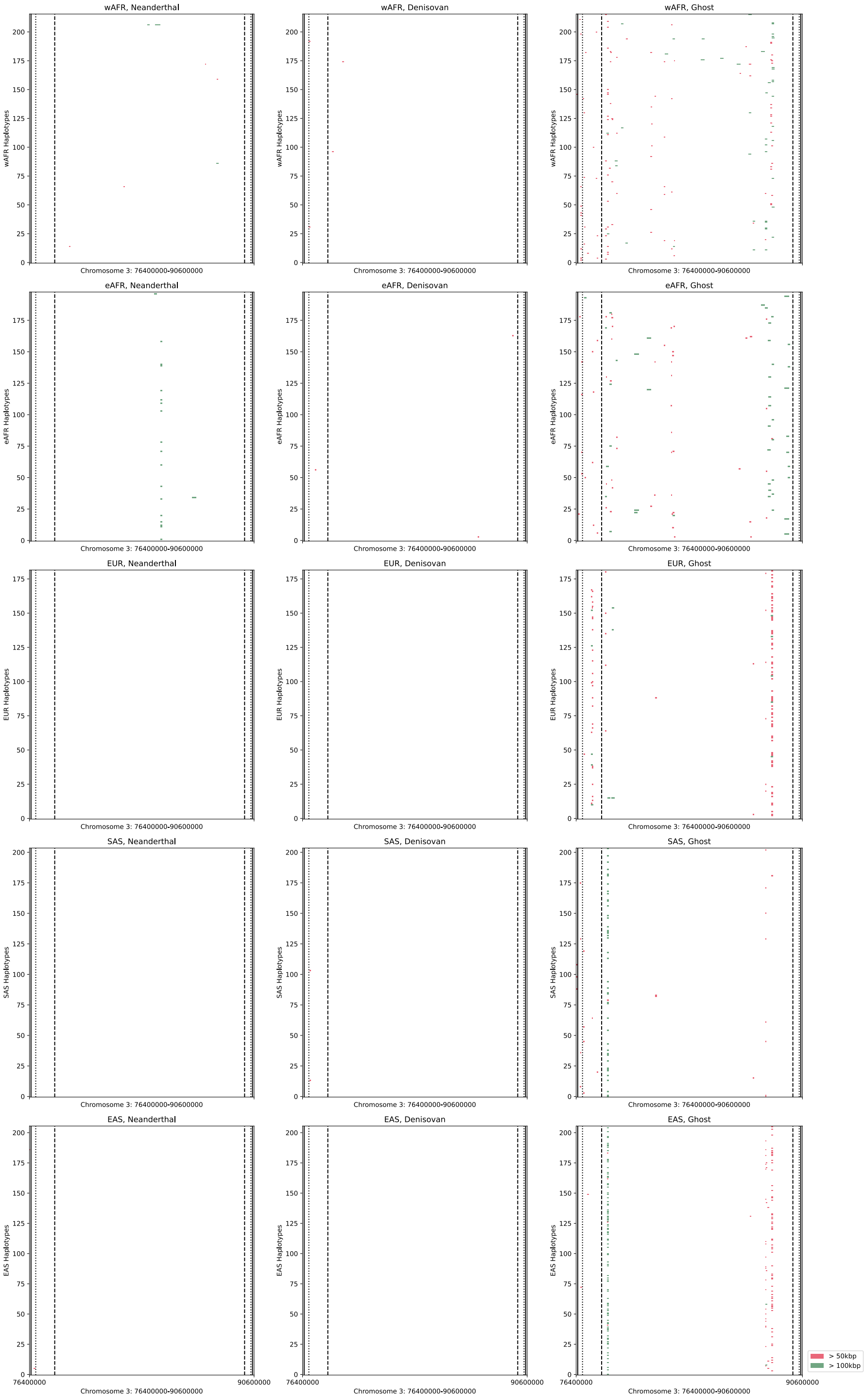

**Fig S41.** Archaic haplotypes in a desert region (chr3:76.4-90.6 Mbp). TRACE-inferred segments for different archaic ancestries are colored by length (green: > 100kbp, red: > 50kbp). Desert boundaries from Vernot et al. 2016 (solid, (*63*)), Sankararaman et al. 2016 (dashed, (*52*)), and Kerdoncuff et al. 2025 (dotted, (*53*)) are shown. Grey shading indicates low-confidence regions (< 50% overlap with the 1000 Genomes strict mask).

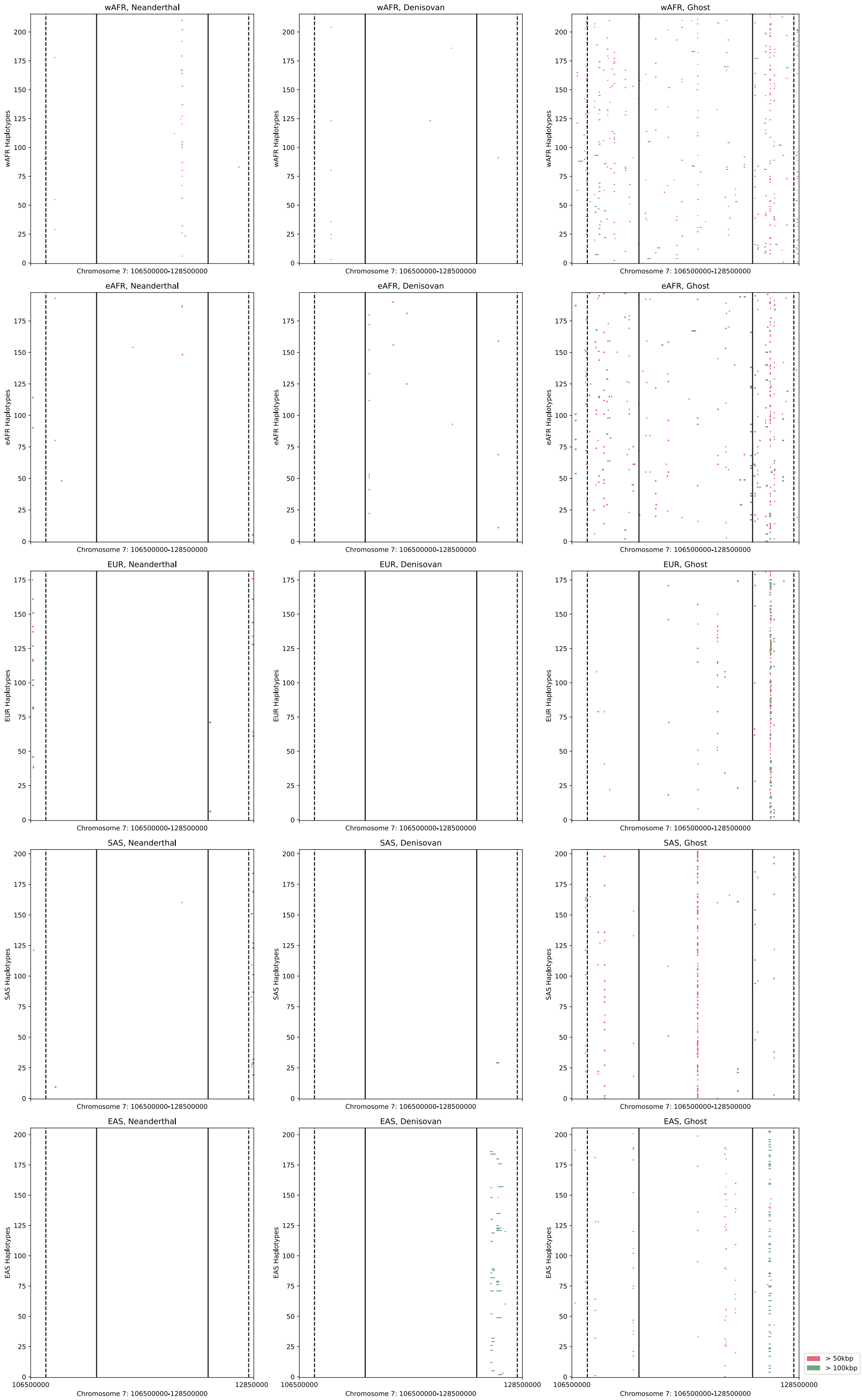

**Fig S42.** Archaic haplotypes in a desert region (chr7:106.5-128.5 Mbp). TRACE-inferred segments for different archaic ancestries are colored by length (green: > 100kbp, red: > 50kbp). Desert boundaries from Vernot et al. 2016 (solid, (*63*)), Sankararaman et al. 2016 (dashed, (*52*)), and Kerdoncuff et al. 2025 (dotted, (*53*)) are shown. Grey shading indicates low-confidence regions (< 50% overlap with the 1000 Genomes strict mask).

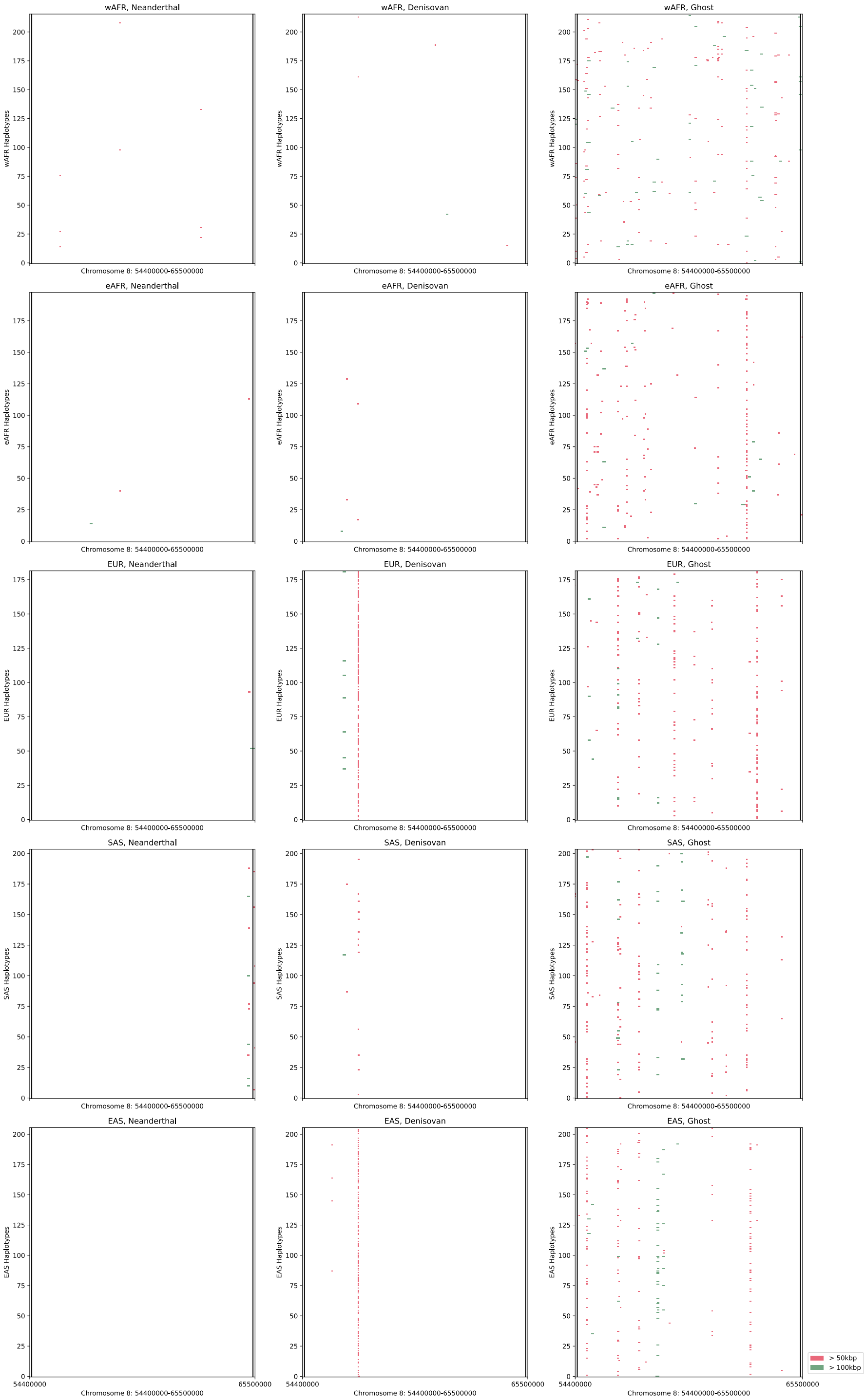

**Fig S43.** Archaic haplotypes in a desert region (chr8:54.4-65.5 Mbp). TRACE-inferred segments for different archaic ancestries are colored by length (green: > 100kbp, red: > 50kbp). Desert boundaries from Vernot et al. 2016 (solid, (*63*)), Sankararaman et al. 2016 (dashed, (*52*)), and Kerdoncuff et al. 2025 (dotted, (*53*)) are shown. Grey shading indicates low-confidence regions (< 50% overlap with the 1000 Genomes strict mask).

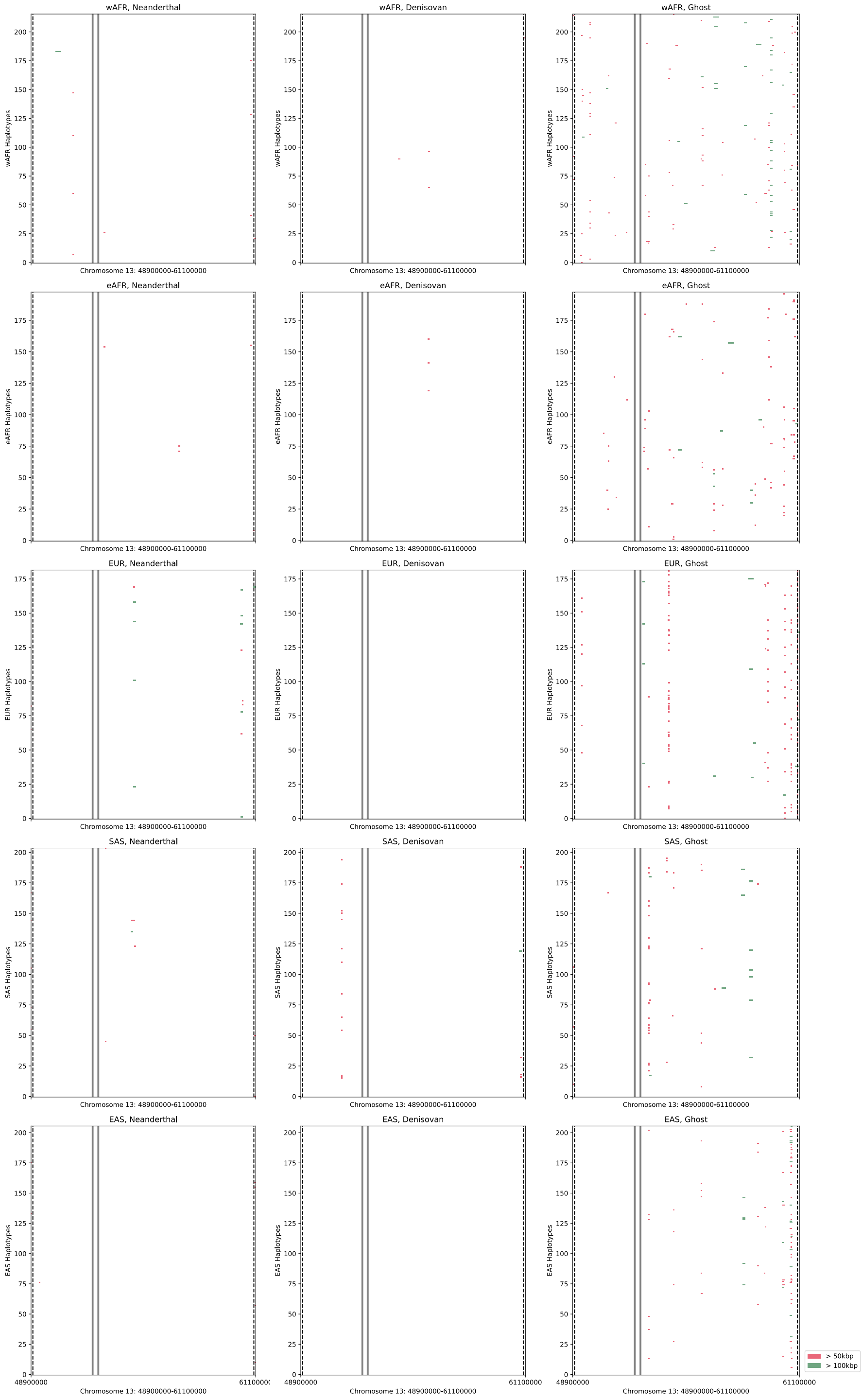

**Fig S44.** Archaic haplotypes in a desert region (chr13:48.9-61.1 Mbp). TRACE-inferred segments for different archaic ancestries are colored by length (green: > 100kbp, red: > 50kbp). Desert boundaries from Vernot et al. 2016 (solid, (*63*)), Sankararaman et al. 2016 (dashed, (*52*)), and Kerdoncuff et al. 2025 (dotted, (*53*)) are shown. Grey shading indicates low-confidence regions (< 50% overlap with the 1000 Genomes strict mask).

### Section S7. Detecting super archaic introgression in Oceanians

#### S7.1 EVOCEANIA dataset processing

To study archaic and super-archaic ancestry in Oceanians, we used whole genome sequences from the EVOCEANIA dataset along with 25 Papuan New Guinea samples from Malspinas et al (*51*, *65*). We jointly genotyped all individuals from gVCF files. Joint genotyping was conducted with *GATK3.8* (*90*) using the GenotypeGVCFs command and using the human reference genome hs37d (**table S1**). We then used *bcftools* to remove indels. Following (*51*), we applied a series of quality control filters. We first applied *GATK4.5* VQSR using the best practices and resource bundles provided by *GATK* and a truth sensitivity filter level of 99.0. (https://gatk.broadinstitute.org/hc/en-us/articles/360035890811-Resource-bundle)

Our command for running VQSR is:

```
gatk VariantRecalibrator \\
-an QC \\
-an MQRankSum \\
-an ReadPosRankSum \\
-an FS \\
-an MQ \\
-an SQR \\
-an DP \\
-mode SNP \\
--resource:hapmap,known=false,training=true,truth=true,prior=15.0
b37_hapmap_3.3.b37.vcf \\
--resource:omni,known=false,training=true,truth=true,prior=12.0
b37_1000G_omni2.5.b37.vcf \\
--resource:1000G,known=false,training=true,truth=false,prior=10.0
b37_1000G_phase3_v4_20130502.sites.vcf \\
--resource:dbsnp,known=true,training=false,truth=false,prior=2.0
b37_dbsnp_138.b37.excluding_sites_after_129.vcf
```

Our command for applying the VQSR filter is:

```
gatk ApplyVQSR -R hd27d5.fa --truth-sensitivity-filter-level 99.0
```

We used *bcftools* to filter any genotypes (by setting them to 'missing') if 1) Genotype Quality (GQ) < 30, 2) Depth (DP) < 10, or 3) DP is more than double the genome-wide depth. We removed any SNPs from the joint dataset that had missing genotypes for > 5% of individuals or had excess heterozygosity (ExcessHet p-value < 0.0001). After applying all filters, we retained 18,090,587 SNPs.

We lifted over the genotyped VCF files from hg19 to hg38 using *CrossMap*. The dataset was then phased with *SHAPEIT5* (*91*) using the 1000 Genomes and HGDP datasets as reference panels (*92*), followed by imputation with *IMPUTE5* (*93*) using the same references. We note that this standard phasing-and-imputation approach would remove variants unique to the Oceanian dataset that are absent from the reference panels. This could significantly impact archaic ancestry inference, particularly for Denisovan segments, as the 1000 Genomes and HGDP populations capture only a limited fraction of the diversity of Denisovan introgression signals in Oceanians. To mitigate this, we added the unique Oceanian variants back into the dataset after imputation and performed a second round of phasing with *SHAPEIT5*, using the already-phased and imputed dataset (merged dataset of shared variants of Oceanians and 1000 Genomes & HGDP references) as a scaffold to guide the phasing and imputation of these unique variants.

From this final dataset, we selected 92 Oceanian individuals including 25 Papuan New Guinea (PNG, (*65*)) and 67 individuals from Santa Cruz Islands (SCI) and Vanuatu Island (VAN) from the EVOCEANIA dataset, along with 107 YRI individuals from the 1000 Genomes Project for analysis using TRACE (**table S3**). We then applied *SINGER*, TRACE, *hmmix*, and *IBDmix* to these samples using the parameters and filters described in Section S4.1. We applied classification *Rule Group 1* (**Section S4.2**) to TRACE-inferred archaic segments for all analysis except inferring super-archaic segments (where we applied *Rule Group 2*), as described in Section S7.4.

#### S7.2 Comparing TRACE with other methods in Oceanian individuals

Using TRACE with $t = 15{,}000$ generations, we inferred an average of 0.73% Neanderthal, 0.66% Denisovan and 0.33% ghost ancestry in Oceanian individuals. Our estimates are lower than *hmmix*, and *IBDmix* in Oceanian individuals, as well as previous studies (*51*) (**fig. S45, table S13**). Specifically, *hmmix* inferred approximately 2% Neanderthal and 2% Denisovan ancestry in these individuals, consistent with initial f4-statistic estimates of 2-3% for both Neanderthal and Denisovan ancestry in a previous analysis of these same individuals (*51*). *IBDmix* inferred about 1.8% Neanderthal ancestry and 1% Denisovan ancestry. Despite this lower total recovery, over 85% of TRACE-detected Neanderthal segments and over 95% of Denisovan segments overlapped with calls from *hmmix* or *IBDmix*.

As Neanderthals and Denisovans coalesce with each other ($t = 13{,}600 - 16{,}900$ generations, (*2*)) before they coalesce with modern humans, many of the long branches characteristic of the archaic introgression are not detected by TRACE when using $t = 15{,}000$ generations as the cutoff. Moreover, the higher archaic introgression proportion further exacerbates this effect, reducing the recall. We replicate these patterns in simulations and show that coalescence between archaic lineages impacts around 10% of Neanderthal and Denisovan segments (**Section S8.2, table S14**). Importantly, while this effect lowers recall for both Neanderthal and Denisovan segments, it does not bias the segments that are successfully identified as evidenced by the consistent overlap between TRACE and other methods (**table S15 vs. table S8, table S13 vs. table S4**). We further validated this intuition by showing that the use of $t = 10{,}000$ generations allows TRACE to recover larger amounts of both Neanderthal and Denisovan ancestries in Oceanians (**fig. S46 vs. Fig. 4A**), recovering similar amount of Neanderthal ancestry as in other non-African populations in 1000 Genomes (**fig. S46 vs. Fig. 3A)**. However, the lower threshold can also increase the false discovery rates, and in turn, detection of ghost ancestry and hence we continue our following analysis with $t = 15{,}000$ generations.

Despite the lower recall, the segments detected by TRACE in Oceanians are robust. They show archaic affinity patterns (**fig. S47**) and site frequency spectra (SFS/cSFS, **fig. S48**) consistent with those observed in 1000 Genomes populations (see **Section S4.4** for how these figures are generated). Ghost ancestry segments detected in Oceanians also overlap with 1000 Genomes populations (**fig. S49**). These empirical patterns are accurately replicated in our simulations (**fig. S55-S56**), and the concordance between TRACE and other methods aligns with simulated expectations (**table S13 vs. table S15**).

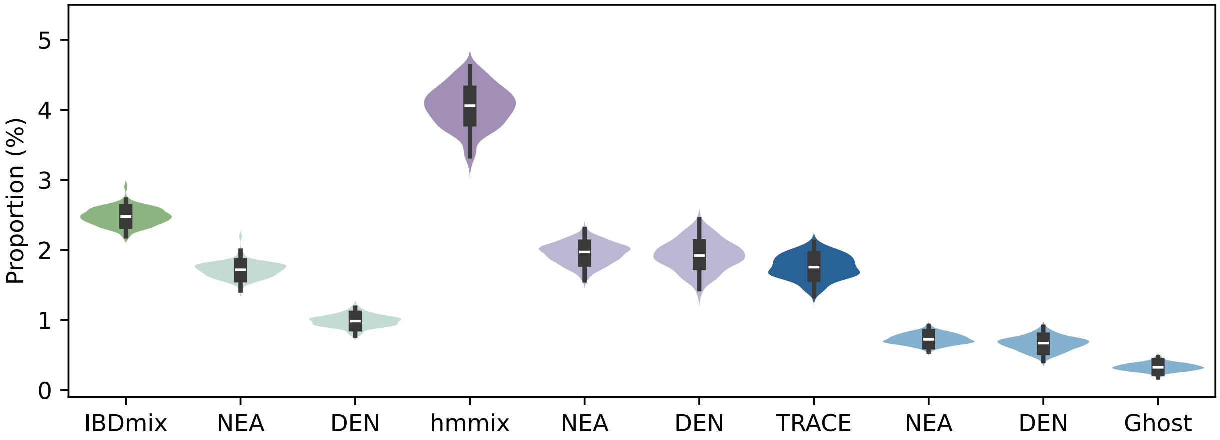

**Fig S45.** Per-individual archaic ancestry across 92 Oceanian samples. Total archaic ancestry (dark colors) and its composition (lighter colors: Neanderthal-NEA, Denisovan-DEN, Ghost) estimated by TRACE (blue), IBDmix (green), and hmmix (purple). All methods used a consistent length filter (> 50kbp, > 0.05cM). TRACE was run with $t$ = 15,000 and posterior probability > 0.9; hmmix with posterior > 0.8; and IBDmix with `slod` > 4.

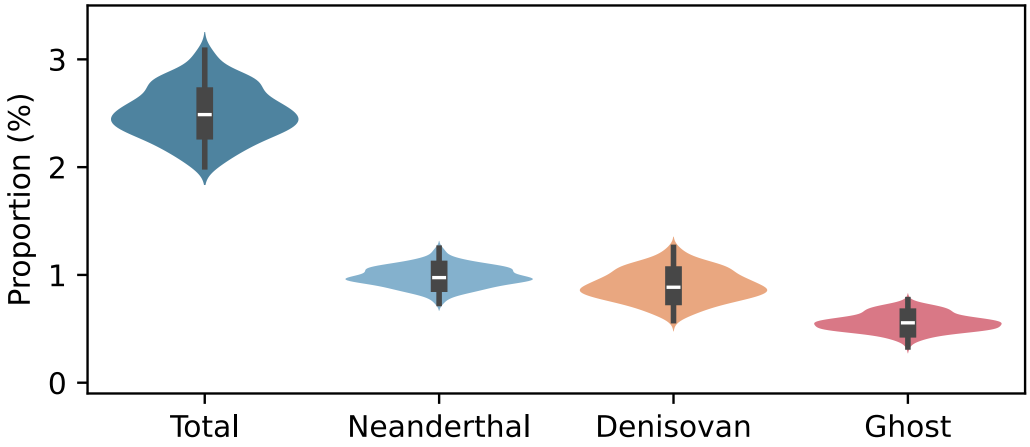

**Fig S46.** Archaic ancestry per individual recovered by TRACE in empirical Oceanian data with t=10000.

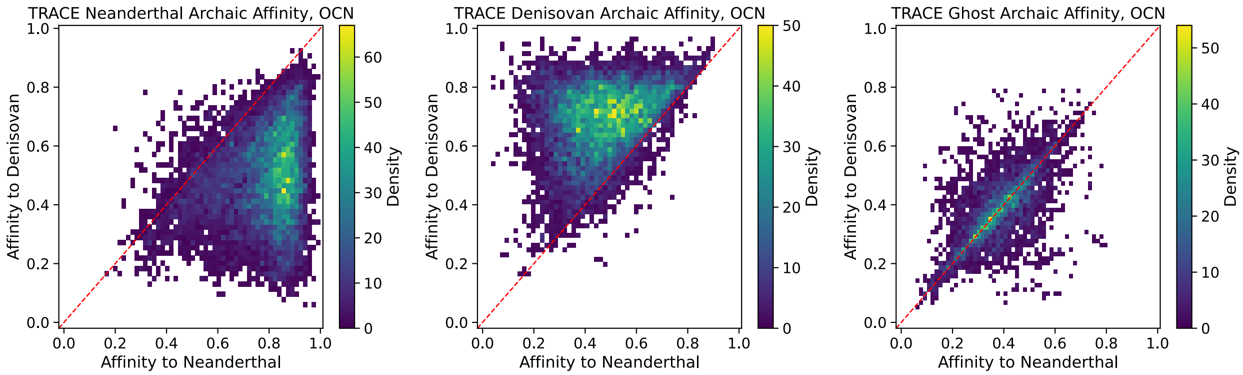

**Fig S47.** Neanderthal versus Denisovan genetic affinity for archaic segments in Oceanians. TRACE-detected segments are classified as Neanderthal (left column), Denisovan (middle), and Ghost (right) ancestry.

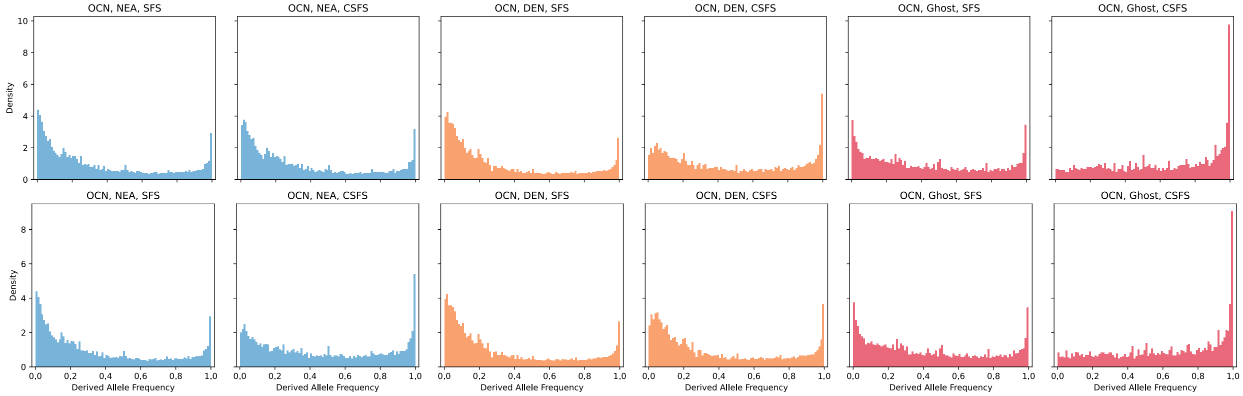

**Fig S48.** Derived allele frequency spectra of archaic segments in Oceanians. (Left) SFS and (Right) conditional SFS (cSFS) conditioned on the derived allele being present in Neanderthals (top) or Denisovan (bottom). Colors indicate TRACE-assigned ancestry: Neanderthal (blue), Denisovan (orange), Ghost (red).

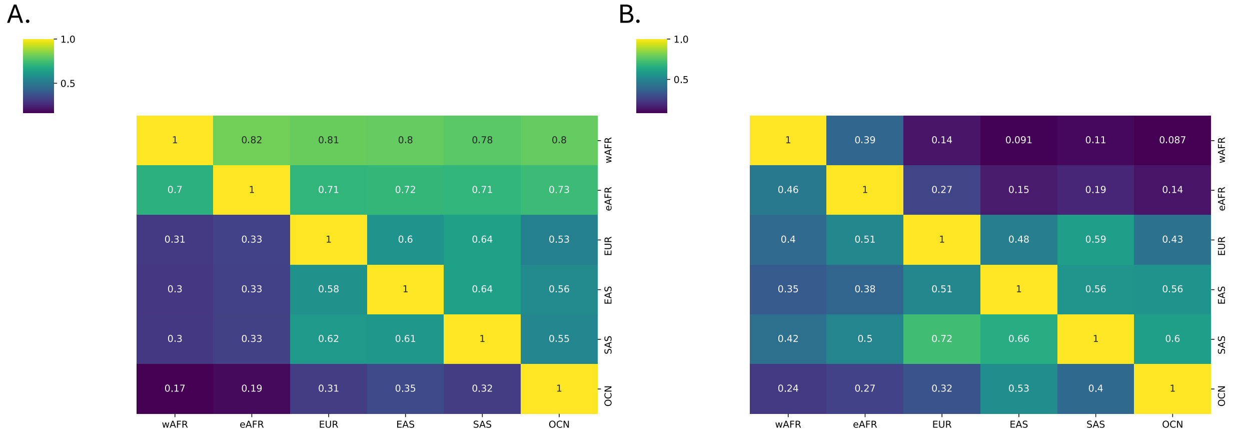

**Fig S49.** Population sharing of archaic ancestry. Heatmap shows the mean proportion of ghost ancestry (**A**) and Neanderthal ancestry (**B**) in each population (x-axis) that is shared with every other population (y-axis). Values represent the average across all individuals within a population.

#### S7.3 Detecting super deep lineages in Neanderthal and Denisovan segments

Comparison of allele sharing patterns between Neanderthals, Denisovans and modern humans has revealed potential signatures of "super-archaic" introgression from an unknown archaic lineage into Denisovans (*1*, *2*). Previous analyses suggest that this lineage diverged from modern humans around 0.7–1.3 million years ago and Denisovans might carry 2.5–6% of ancestry from this population (*2*, *27*). As Oceanians harbor significantly more Denisovan ancestry than other non-African populations, we reasoned that some fraction of super-archaic ancestry may persist in introgressed Denisovan segments in present-day Oceanians.

To test for super-archaic ancestry within Denisovan-derived segments, we analyzed the marginal trees in ARGs within the recovered Denisovan segments in Oceanians. We classified introgressed "long branches" with upper-end coalescence times greater than 31,500 generations (913,500 years ago; following (*27*)) as "super-deep lineages". For comparison, we also characterize the proportions of super-deep lineages in Neanderthal introgressed segments. We focused on genomic regions longer than 10kbp and 0.01cM within Neanderthal and Denisovan segments containing these super-deep lineages. By aggregating these super-deep segments across all individuals, we constructed a "synthetic" super-archaic genome (defined as the union of all super-archaic segments) in the "synthetic" Neanderthal and Denisovan genomes (defined as the union of all Neanderthal / Denisovan segments). We calculated the proportion of the synthetic Neanderthal and Denisovan genomes composed of super-archaic segments (**Fig. 4**). Uncertainty was quantified by using a chromosome-weighted jackknife approach, which involved iteratively removing one chromosome, recalculating the super-archaic ancestry proportion and weighting the results by chromosome size (*94*).

We find the proportion of super-archaic ancestry is significantly higher in Denisovan segments than in Neanderthal segments ($p < 2.2 \times 10^{-308}$, binomial test; **Fig. 4C**). This statistical test was applied to all pairwise comparisons between chromosome-weighted jackknife samples, with the null hypothesis that a randomly selected Denisovan background jackknife sample would have more super-archaic segments than a Neanderthal background jackknife sample with a probability of 0.5.

Based on simulations, this result is unexpected under a demographic model without super-archaic introgression and suggests introgression from a very deep lineage into the Denisovans (**Section S8**).

#### S7.4 Inferring super-archaic segments embedded in Denisovan segments

To identify a greater number of super-archaic segments within Denisovan ancestry in modern humans, we applied a modified classification scheme (*Rule Group 2*, **Section S4.2**) designed to correctly classify Denisovan segments containing embedded super-archaic ancestry. Unlike the stringent *Rule Group 1*, which requires high affinity to the sequenced Denisovan genome for classification, *Rule Group 2* leverages our prior findings that Africans carry minimal Denisovan ancestry and that most ghost ancestry is shared between Africans and non-Africans. This approach helps identify those segments that contain a high proportion of super-archaic ancestry as

"Denisovan", which would otherwise exhibit reduced affinity to the sequenced Denisovan genome and be misclassified as “ghost”. In simulations, this strategy increased the recall of super-archaic segment detection from 40% to 72%. (**table S16, Section S8.4**).

Our simulations showed that super-archaic fragments (that are not present on the sequenced Denisovan genome) exhibit characteristics similar to ghost segments: deep coalescence times with modern humans, long genomic lengths, and low affinity to both Neanderthal and Denisovan sequenced genomes (**Section S8.3-S8.4, fig. S58**). When applied to simulated data, this classification scheme detected super-archaic segments with 70.2% accuracy (95% CI: 66.4–74.1%, **Section S8.5**).

Applying the classification scheme described above to Oceanians, we inferred 0.25% super-archaic ancestry within the total detected Denisovan ancestry (**table S17**). We note that this value is a conservative lower bound, because our analysis is limited to Denisovan-introgressed segments in modern humans, and TRACE recovers only a subset of such segments under the chosen parameters (**fig. S59**).

The super-archaic segments identified from Denisovan introgression in Oceanian genomes have a mean length of 37,559 bp (**fig. S50**). However, this reflects a length-biased subset, since TRACE only recovers segments longer than ~20kbp—the mean length of ILS segments estimated from over 10kbp, 0.01cM segments subtended by “super-deep” lineages in Neanderthal introgressed regions (**Section S8.5**).

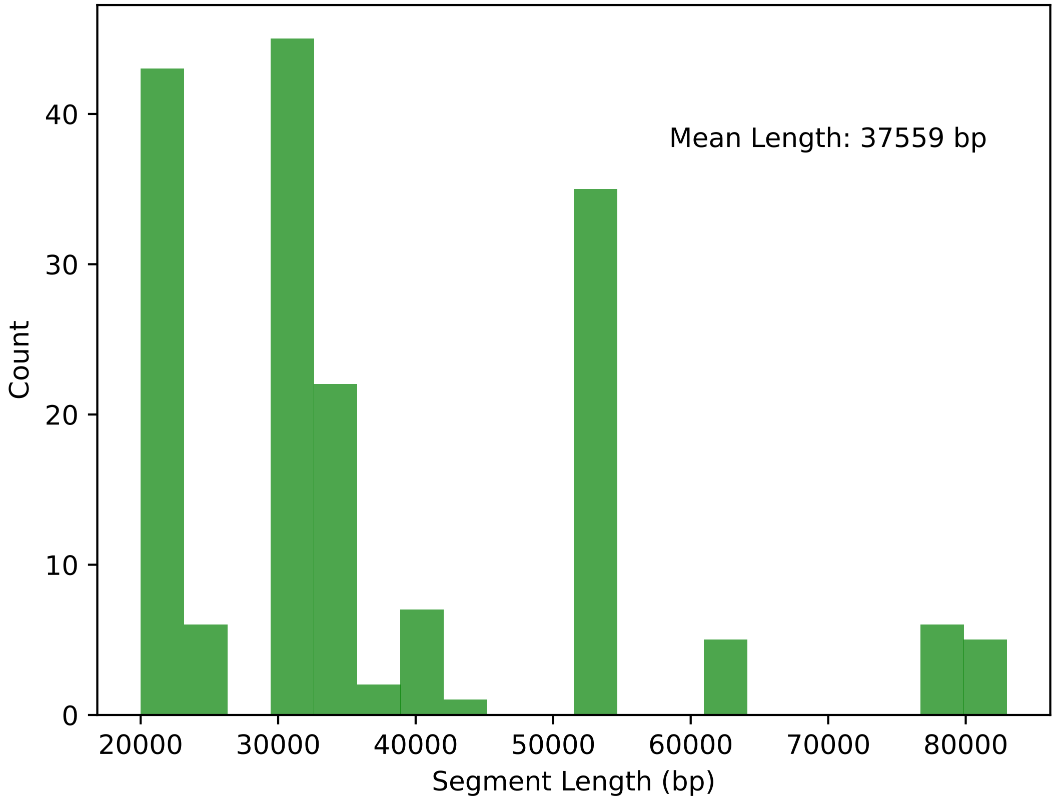

**Fig S50.** Haplotype length distributions of detected super-archaic segments in all Oceanian individuals.

#### S7.5 Functional annotation of detected super-archaic segments

We sought to characterize the functional impact of super-archaic segments detected by TRACE using two analyses: 1) identifying genes that are enriched within super-archaic segments, and 2) gene-ontology (GO) enrichment analyses. We used the protein coding gene annotations in Gencode v49 (*88*).

We find super-archaic ancestry is enriched in the broader HLA region, encompassing *RNF39, PPP1R11,* and *POLR1H* (**fig. S51-S52**). Denisovan ancestry has previously been shown to be enriched at the HLA locus (*53*), the long genealogical branches resulting from long-term balancing selection (*36*) within the region may also lead to the "super-archaic" like signal we find. We also find a high-frequency super-archaic ancestry within *CYP24A1*, a part of the cytochrome P450 family and a critical regulator of vitamin D degradation in humans (*65*, *66*) (**Fig. 4B**). Variants in *CYP24A1* are consequently associated with serum calcium measurements (*95*) and with a recessive form of severe hypercalcemia in children (*96*).

We evaluated the pathway enrichment of super-archaic segments using GREAT (87). This analysis identified 72 significantly enriched pathways at the region level (Binomial test), and 0 at the gene level (Hyper-geometric test) after multiple testing correction (**fig. S53**). The primarily enriched targets are components related to the MHC Class 1 as well as Activity-related cytoskeleton (ARC) associated regions (**fig. S53**).

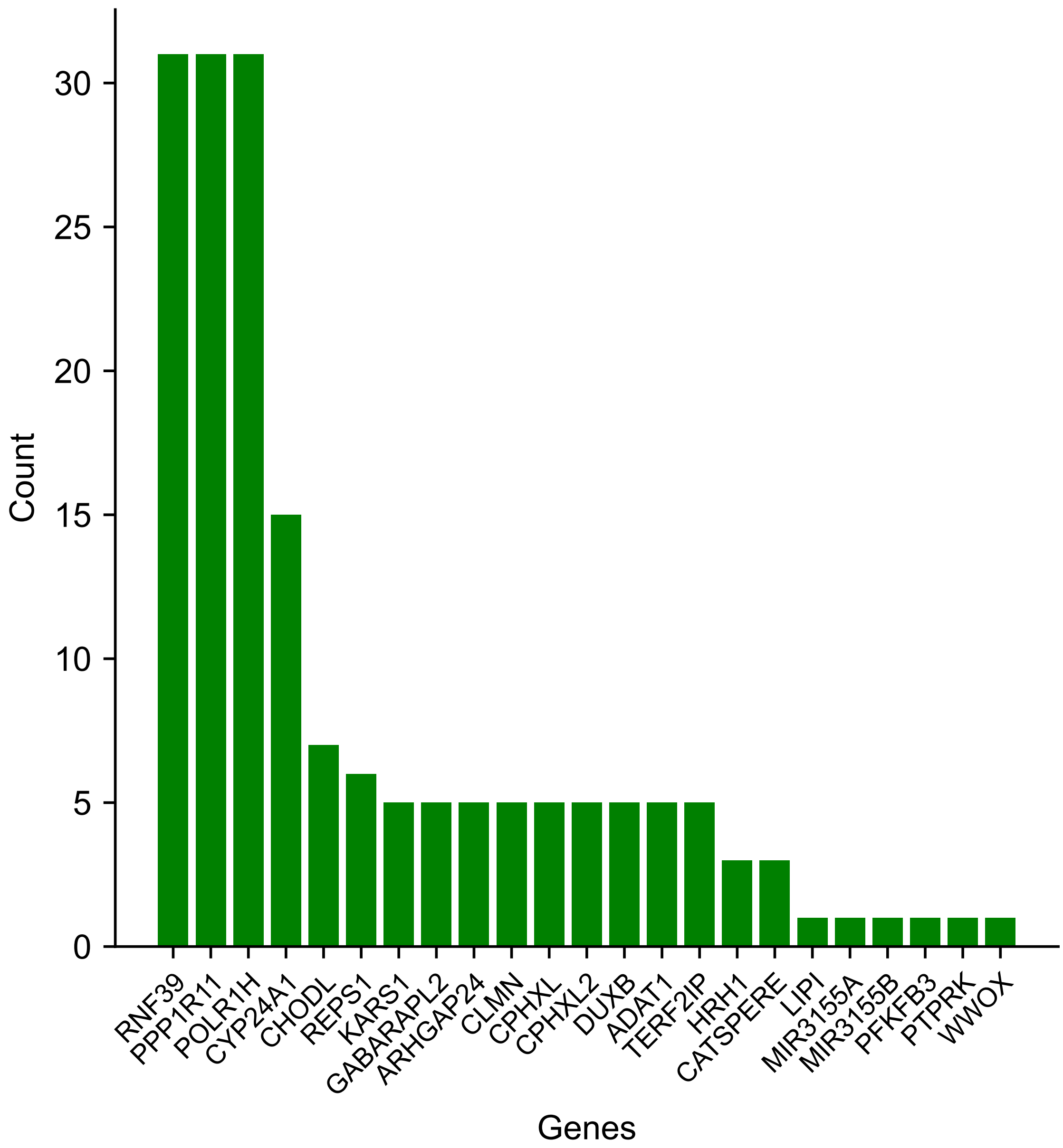

**Fig S51.** Common gene intersections with calls of super-archaic ancestry in TRACE. Each count reflects an intersection within a single haplotype (e.g., a superarchaic call split over a long gene such that two tracts overlap with the gene will only count as a single occurrence). Note that some genes are not plotted for visualization purposes.

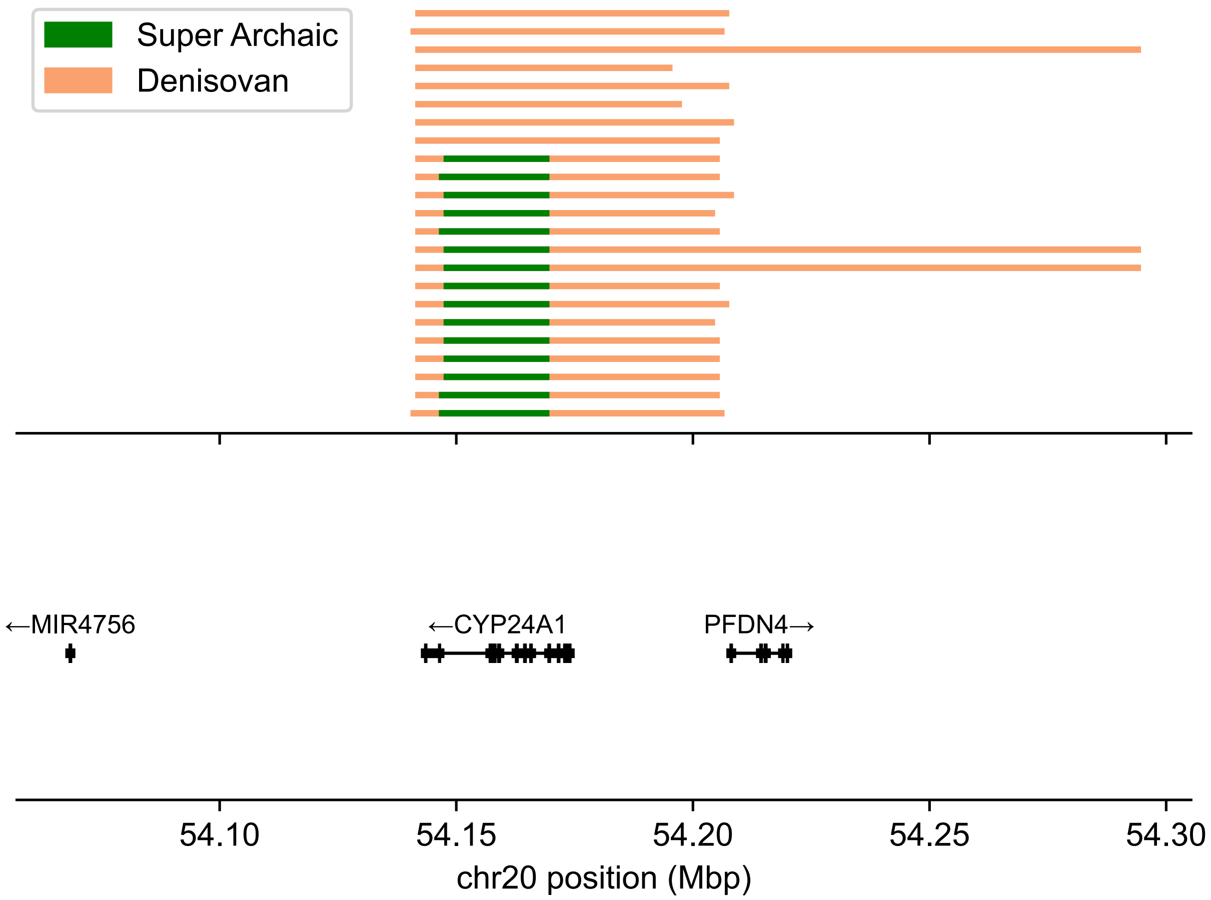

**Fig S52.** Zoomed intersection into genes overlapping a high-frequency cross-sample call of superarchaic ancestry using TRACE on chromosome 20.

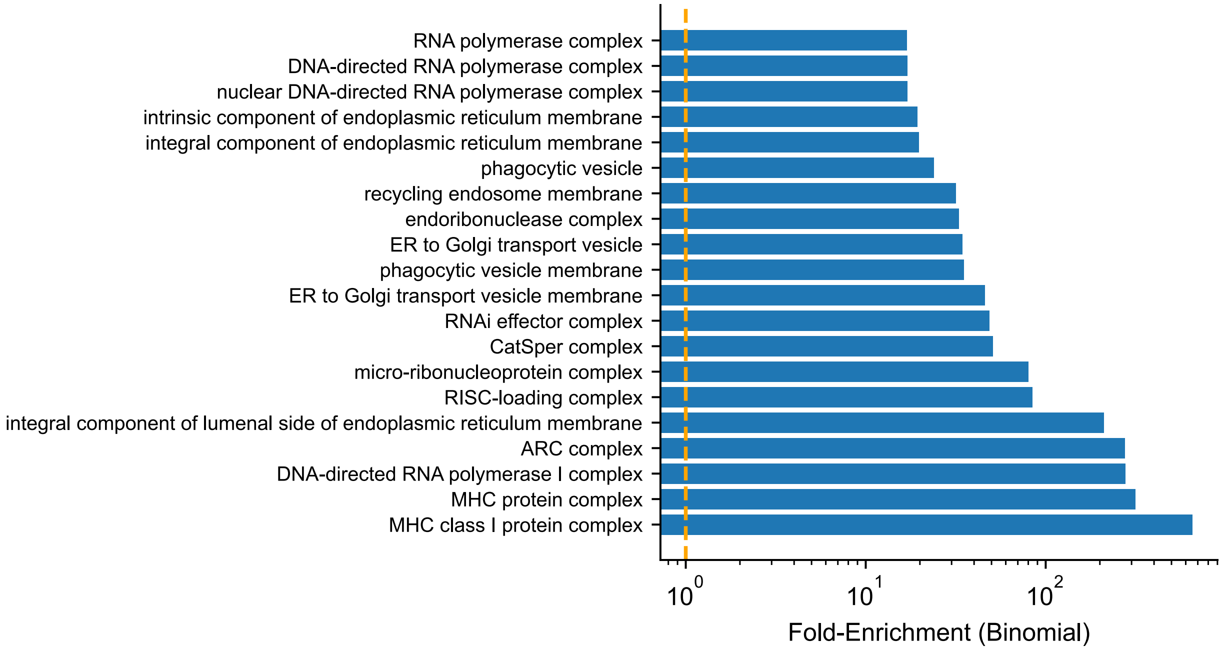

**Fig S53.** Enrichment of gene ontologies in superarchaic segments inferred by TRACE. Top twenty enriched categories are shown for categories exhibiting significant region-specific enrichment.

### Section S8. Validating patterns of super-archaic ancestry using simulations

#### S8.1 Simulation models

To validate the signal of super-archaic introgression, we developed two demographic models based on the OOA model (**fig. S26; Section S5.1**), tailored to reflect Oceanian (OCN) and African population history.

1. "No Super-Archaic" Model (**fig. S54A**): This baseline model includes:
    - A single pulse of 3% Denisovan introgression into the non-African ancestral population 45,000 years ago.
    - No subsequent back-migration from non-Africans into Africa.
2. "Super-Archaic" Model (**fig. S54B**): This model extends the "No Super-Archaic" model by adding a super-archaic lineage, with parameters informed by (*27*):
    - A divergence from the modern human lineage 910,000 years ago.
    - A single pulse of 6% introgression from the super-archaic population into the ancestral Denisovan population 300,000 years ago.

For each model, we generated 10 simulation replicates of 50 Mbp genomes. From each replicate, we sampled 100 non-African and 100 African individuals, alongside one diploid individual each from the sequenced Neanderthal and Denisovan populations. We then applied *SINGER*, TRACE, *hmmix*, and *IBDmix* using the standard settings detailed in Section S4.1.

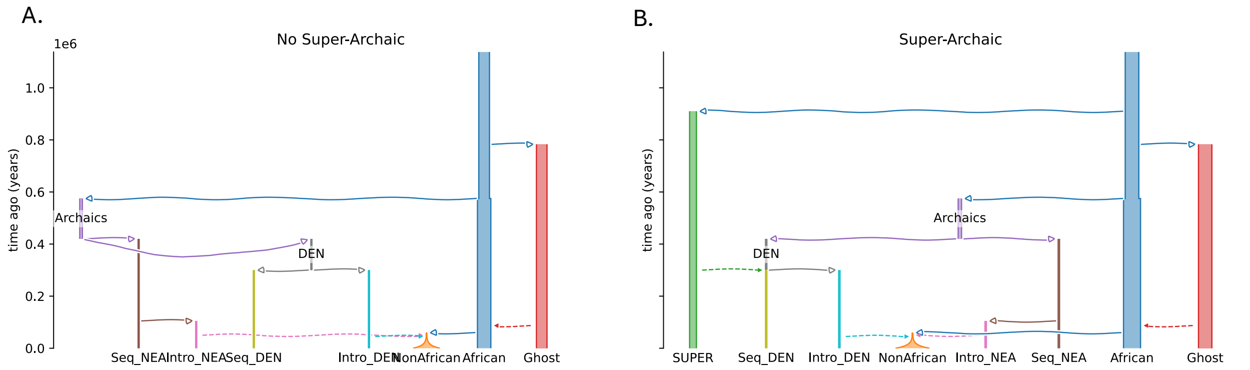

**Fig S54.** Demographic models for super-archaic introgression validation. **(A)** No SuperArchaic model: a model recapitulating known demographic history for Oceanians. **(B)** SuperArchaic model: model in A incorporating introgression from a super-archaic lineage into Denisovans alone. Parameters following (*27*).

#### S8.2 Concordance between simulation and real data

We assessed the precision and recall of TRACE, *hmmix* and *IBDmix* and checked concordance of archaic segment calls among these methods using the same approach as described in Section S5.2. We found high concordance between the empirical data and simulation outputs. As shown in table S14, the relative performance of the methods in simulations mirrors the patterns in real data (**fig. S45**): Among the three methods, *hmmix* recovers the highest amount of archaic ancestry. Both TRACE and *hmmix* show comparable recall for Neanderthal and Denisovan ancestry, whereas *IBDmix* shows lower recall for Denisovan ancestry than Neanderthal ancestry as the reference Denisovan sampled from simulation is more divergent from the introgressing Denisovan than the reference Neanderthal. Interestingly, the precision for identifying Neanderthal ancestry is lower than for Denisovan across all methods. This is due to the misclassification of Denisovan segments as Neanderthal, which stems from the higher divergence between the reference Denisovan and introgressing Denisovan population—this leads to misclassification of some ND11 sites as ND10 and hence these segments are assigned to the Neanderthal lineage, instead of Denisovan.

Among the three methods, TRACE exhibits lower recall for both Neanderthal and Denisovan ancestry. Notably, TRACE recovered less Neanderthal ancestry in these Oceanian simulations than in the European simulations from Section S5 (**table S7 vs. S14**). In these simulations, as Neanderthals and Denisovans coalesce with each other 420,000 years ago (14482 generations), many of the long branches of the archaic introgression are not detected by TRACE when using $t = 15{,}000$ generations as the cutoff. Moreover, the higher archaic introgression proportion further exacerbates this effect, as a larger fraction of coalescent events occur among archaic lineages (**fig. S57**). These events break the long, distinct branches that TRACE relies on for detection of archaic segments and thereby reduces recall by ~10% for both Nenaderthal and Denisovan detection (**table S14**). However, this does not bias the segments that are successfully identified, as evidenced by the consistent overlap patterns between TRACE and other methods (**table S15**).

Finally, we generated archaic affinity plots, SFS, and conditional SFS (cSFS) as previously described (**Section S4.4, fig. S55-S56**). The results from the "Super-Archaic" model simulations showed high consistency with the patterns observed in the real Oceanian data (**fig. S47-S48**), as well as with empirical patterns in 1000 Genomes non-African populations (**fig. S17-S21**).

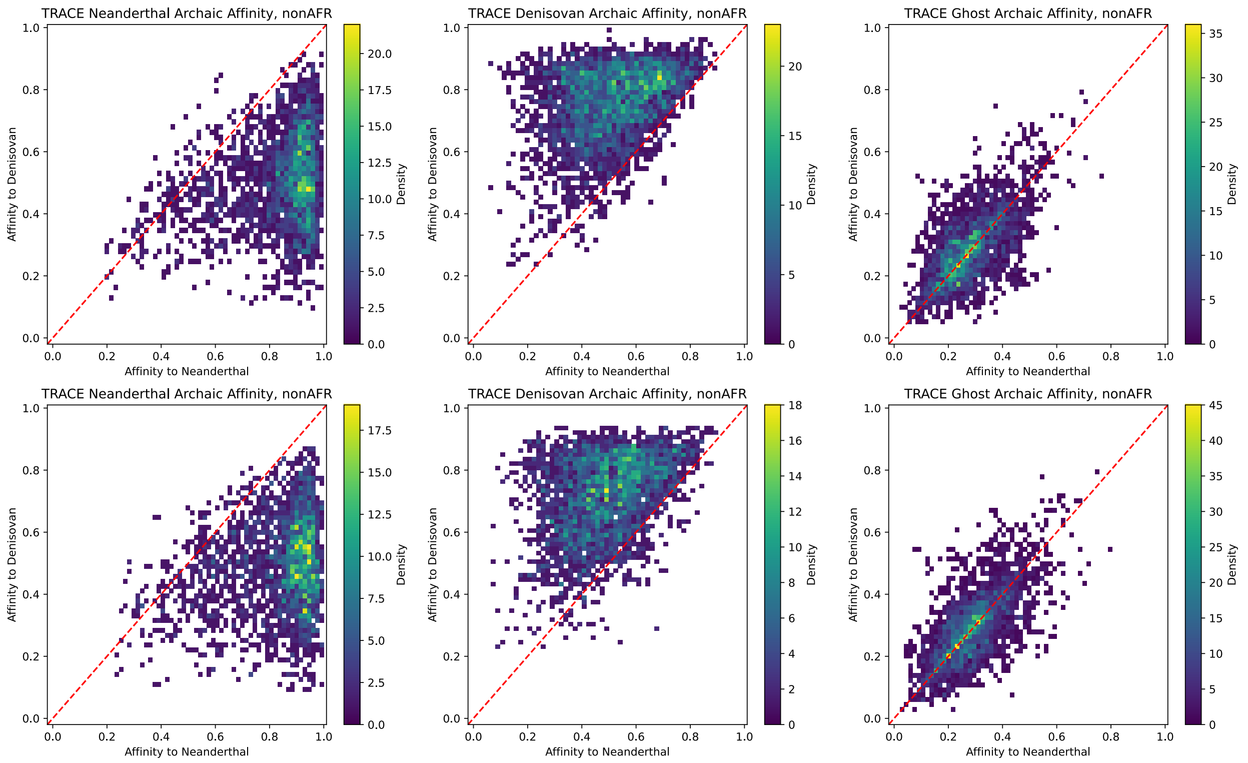

**Fig S55.** Archaic segment affinity in simulated models. Neanderthal vs. Denisovan affinity for TRACE-classified segments (columns: Neanderthal, Denisovan, Ghost) under No Super-Archaic (top row) and Super-Archaic (bottom row) models.

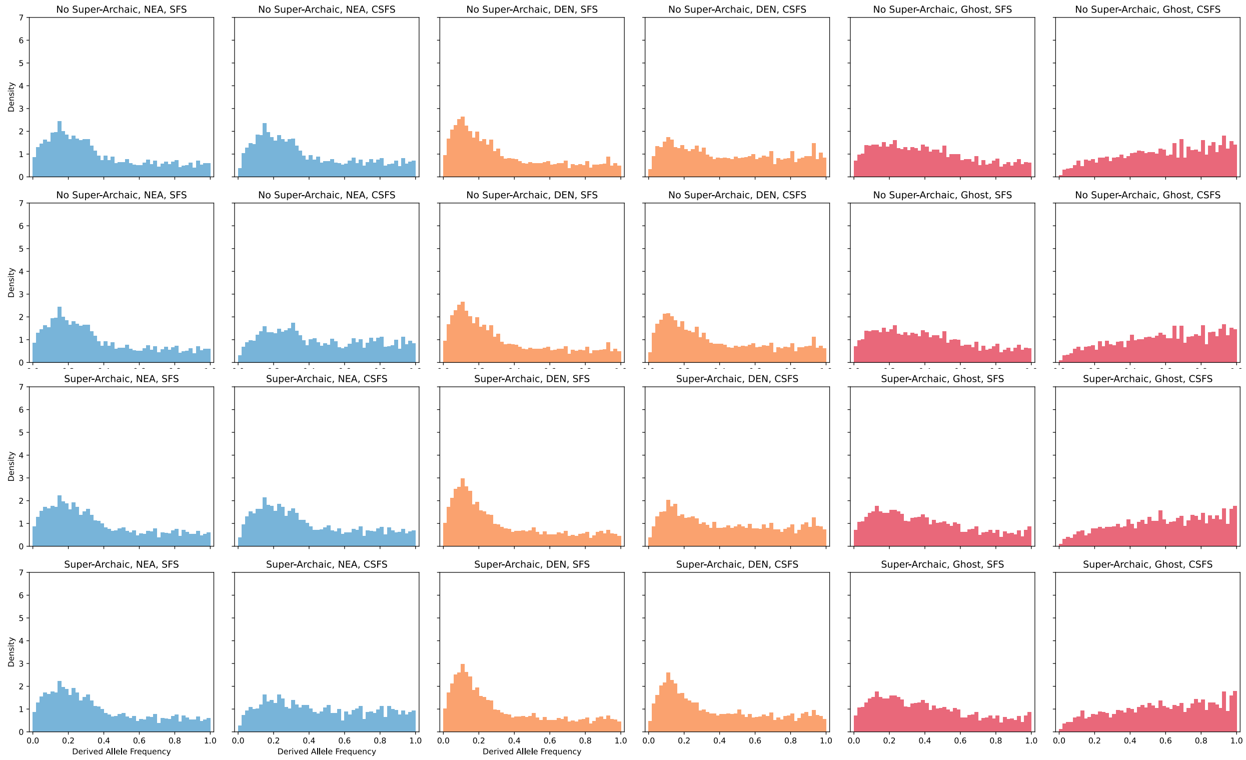

**Fig S56.** SFS and cSFS of archaic segments in simulated models. Results are shown for No Super-Archaic (top two rows) and Super-Archaic (bottom two rows) models. Left: SFS; Right: cSFS conditioned on Neanderthal (rows 1,3) or Denisovan (rows 2,4). Colors indicate ancestry type (Neanderthal: blue, Denisovan: orange, Ghost: red).

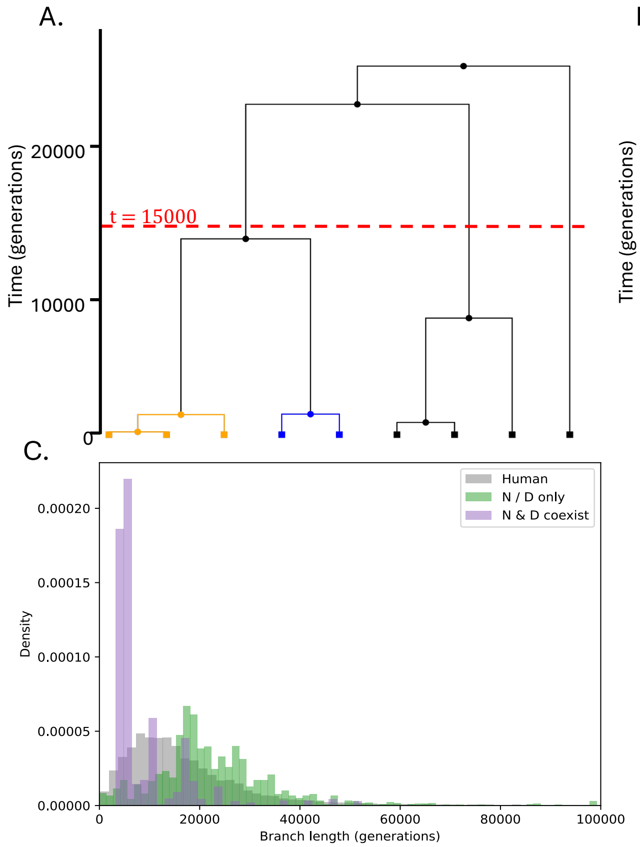

**Fig S57.** Coalescence between archaic lineages. Examples of marginal trees under the No Super-Archaic model with **(A)** both Neanderthal (blue) and Denisovan (orange) introgression and (**B**) Denisovan introgression only. **(C)** Branch length distributions for branches spanning t=15,000 generations and subtending the target haplotype. Colors indicate the simulation ground truth: gray (modern human ancestry only), green (one archaic lineage present), purple (multiple archaic lineages present).

#### S8.3 Detecting super-deep lineages in simulations

We screened for super-deep lineages within the detected Neanderthal and Denisovan segments from both the "No Super-Archaic" and "Super-Archaic" simulation models, following the method in Section S7.3. We quantified uncertainty using a jackknife procedure across the 10 simulation replicates. In the "No Super-Archaic" model, there is no significant difference between the proportion of super-deep lineages within Denisovan and Neanderthal segments ($p = 1.00$; Binomial Test). In contrast, the "Super-Archaic" model successfully recapitulated this key pattern, providing strong simulation-based support for introgression from a super-archaic lineage into the Denisovans ($p < 10^{-36}$; Binomial Test) (**Fig. 4**).

#### S8.4 Signals differentiating super-archaic tracts in Denisovan segments

Leveraging simulation ground truth, we characterized the lineages identified as super-archaic. First, we confirmed that the upper-end coalescence times of introgression branches in super-archaic regions are significantly older than those in Neanderthal or Denisovan segments (without super-archaic ancestry), consistent with the deep divergence of the super-archaic lineage (**fig. S58A**).

We further analyzed the mutation profiles of these segments—specifically the $P_{ND00}$, $P_{ND01}$ and $P_{sYRI}$ statistics (see **Section S4.2** for definitions)—for super-archaic segments that were embedded within detected Denisovan segments but not observed in the sequenced Denisovan genome (**fig. S58B-D**). We find that the super-archaic segments found in Oceanian genomes exhibited high $P_{ND00}$ (indicating divergence from both sequenced archaic genomes), low $P_{ND01}$ (indicating low affinity to the sequenced Denisovan), and low $P_{sYRI}$ (reflecting the scarcity of Denisovan ancestry in Africans). Notably, when the sequenced Denisovan genome itself contains super-archaic ancestry at these loci, introgressed super-archaic segments would display high Denisovan affinity comparable to other Denisovan segments, inhibiting detection of super-archaic signal from the Denisovan ancestry background.

Finally, to distinguish true super-archaic introgression from segments with deep coalescence arising from ILS, we compared the physical length and number of introgressed mutations for deep segments (> 31,500 generations) embedded in Neanderthal segments (ILS) versus true super-archaic segments (**fig. S58E-F**). As predicted, true super-archaic segments were longer and carried more introgressed mutations, consistent with their origin from a deep introgression event that introduced long, divergent haplotypes.

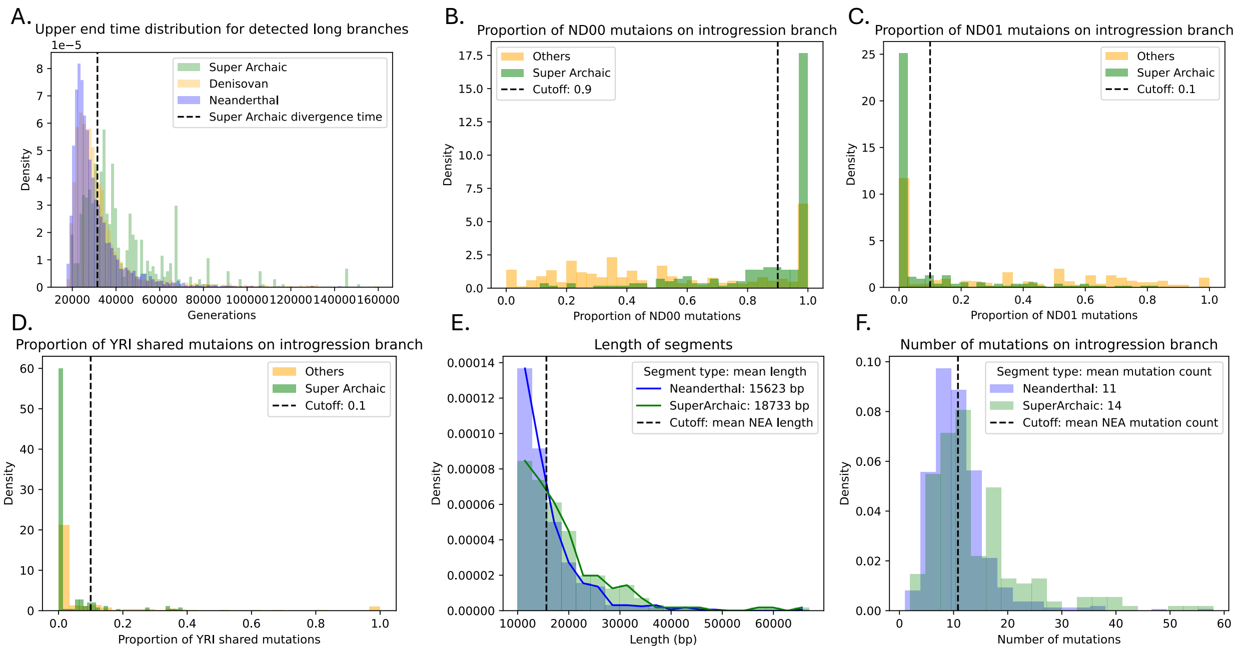

**Fig S58.** Characteristics of super-archaic segments in simulations. Features are shown for segments labeled by simulation ground truth: Neanderthal (blue), Denisovan (orange), and super-archaic (green). **(A)** Distribution of introgression branch upper-end times. (B-D) Proportion of mutations mapped to the "super-deep" branch for ND00 **(B)**, ND01 **(C)**, and YRI-shared **(D)** sites. **(E)** Length distributions of segments subtended by "super-deep" branches (> 0.01cM). **(F)** Number of mutations on the "super-deep" branch. See Section S8.2 for the definition of "super-deep" branches.

#### S8.5 Inferring super-archaic segments in simulations

Standard classification rules (*Rule Group 1*), which require high affinity to the sequenced Denisovan genome, can misclassify Denisovan segments with substantial super-archaic ancestry as "Ghost" segments due to their reduced Denisovan affinity. To mitigate this, we developed modified classification rules (*Rule Group 2*, **table S5**) that leverage the distinct population distribution of Denisovan and ghost ancestry: namely, that Denisovan ancestry is largely absent from Africans, while ghost ancestry is shared across all modern humans. In simulations, *Rule Group 2* correctly classified over 70% of segments containing super-archaic ancestry as Denisovan, while maintaining high accuracy and recall for standard Denisovan segments (**table S16**).

Guided by simulation results (**fig. S58**), we established specific criteria to identify super-archaic segments within the classified Denisovan ancestry. A candidate segment was defined as a continuous region (≥ 0.01cM, 10kbp) of consecutive marginal trees, each containing introgression branches with upper-end coalescence times > 31,500 generations. From these candidates, we classified a segment as super-archaic if it met all of the following thresholds:

- Mutation Profile: $P_{ND00} > 0.9, P_{ND10} = 0, P_{ND01} < 0.1, P_{sYRI} < 0.1$
- Segment Size: Segment length and number of introgressed mutations both exceeded the mean values observed for deep-coalescent segments within Neanderthal ancestry (ILS segments with upper-end coalescence time > 31,500 generations, ≥ 0.01cM, 10kbp).

We validated this inference framework on simulations with known super-archaic segments. The criteria identified segments that overlapped with ground-truth super-archaic regions with 70.2% accuracy (95% CI: 66.4% - 74.1%, accessed through jackknife across 10 simulation replicates; **table S17**).

Finally, we quantified the sensitivity of our approach. Our inference method recovers only 3.68% of the total simulated super-archaic ancestry that introgressed into modern humans (**fig. S59**). This indicates that our empirical estimate of super-archaic ancestry represents a substantial lower bound.

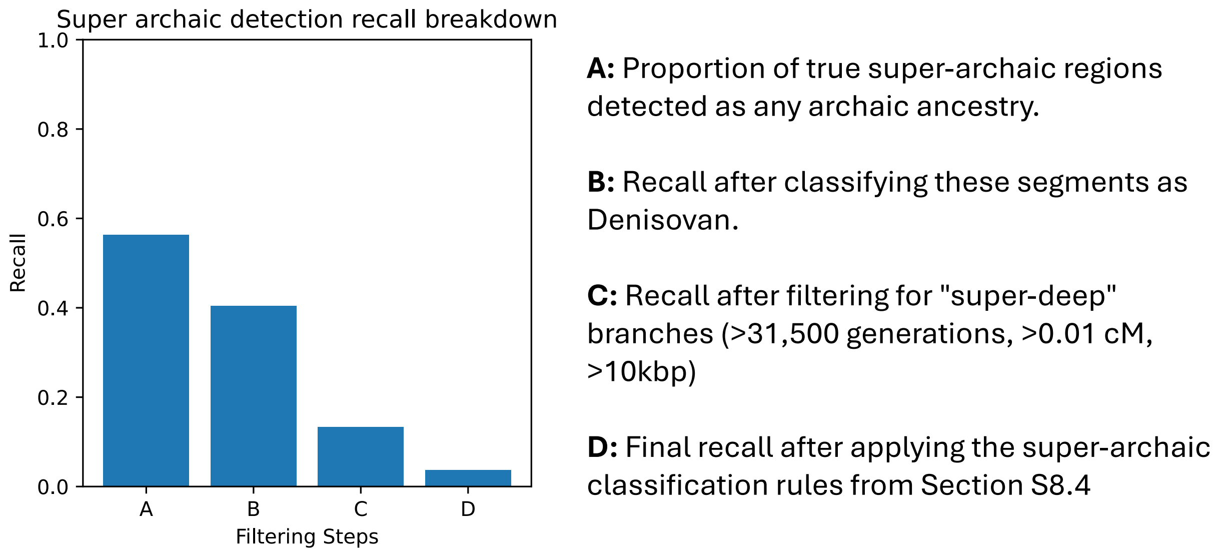

**Fig S59.** Step-wise recall in super-archaic detection. Cumulative recall of super-archaic segments at successive stages of the super-archaic inference pipeline in simulations under the Super-Archaic model. **(A)** Proportion of true super-archaic regions detected as any archaic ancestry. **(B)** Recall after classifying these segments as Denisovan. **(C)** Recall after filtering for "super-deep" branches (> 31,500 generations, > 0.01cM). **(D)** Final recall after applying the super-archaic classification rules from Section S8.4.

### Supplementary Tables

**Table S1**. Resources used in this paper

| Data | Source | Identifier |
|---|---|---|
| 1000 Genomes Whole Genome Sequence | IGSR | https://ftp.1000genomes.ebi.ac.uk/vol1/ftp/data_collections/1000G_2504_high_coverage/working/20220422_3202_phased_SNV_INDEL_SV/ |
| 1000 Genomes and HGDP combined phasing reference | Broad Institute | https://gnomad.broadinstitute.org/downloadsv3-hgdp-1kg |
| EVOCEANIA | EGA | EGAS00001004540 |
| Papuan New Guinea whole genome sequences | dbGAP | phs001085.v1.p1 |
| Altai Neanderthal | MPI | http://ftp.eva.mpg.de/neandertal/Vindija/FilterBed/Altai/ |
| Chagyrskaya Neanderthal | MPI | http://ftp.eva.mpg.de/neandertal/Chagyrskaya/FilterBed/ |
| Vindija Neanderthal | MPI | http://ftp.eva.mpg.de/neandertal/Vindija/FilterBed/Vindija33.19/ |
| Altai Denisovan | MPI | http://ftp.eva.mpg.de/neandertal/Vindija/FilterBed/Denisova/ |
| Human ancestral genome | Ensembl | https://ftp.ensembl.org/pub/release-86/fasta/ancestral_alleles/homo_sapiens_ancestor_GRCh38_e86.tar.gz |
| Genetic Map | Hapmap Project | https://alkesgroup.broadinstitute.org/Eagle/downloads/tables/genetic_map_hg38_withX.txt.gz |
| 1000 Genomes Strict Mask | IGSR | https://www.internationalgenome.org/announcements/genome-accessibility-masks/ |
| Reference genome hs37d | GATK | https://ftp.1000genomes.ebi.ac.uk/vol1/ftp/technical/reference/phase2_reference_assembly_sequence/ |
| Gencode | Gencode | https://www.gencodegenes.org/human/release_49.html |

**Table S2.** Software and algorithms

| Name | Reference | Version |
|---|---|---|
| msprime | https://tskit.dev/msprime/docs/stable/installation.html | 1.2.0 |
| tskit | https://tskit.dev/tskit/docs/stable/installation.html | 0.5.5 |
| hmmix | https://github.com/LauritsSkov/Introgression-detection | 0.8.0 |
| Sprime | https://github.com/browning-lab/sprime | |
| IBDmix | https://github.com/PrincetonUniversity/IBDmix | |
| Relate | https://myersgroup.github.io/relate/ | v1.1.5_x86_64_static |
| SINGER | https://github.com/popgenmethods/SINGER | 0.1.8 |
| bcftools | https://samtools.github.io/bcftools/bcftools.html | 1.6 |
| pybedtools | https://daler.github.io/pybedtools/ | 0.10.0 |
| bedtools | https://bedtools.readthedocs.io/en/latest/ | v2.31.1 |
| GATK | https://github.com/broadinstitute/gatk/releases | v3.8.1, v4.5.0 |
| CrossMap | https://crossmap.readthedocs.io/en/latest/ | v0.7.0 |
| SHAPEIT5 | https://odelaneau.github.io/shapeit5/ | 5.1.1 |
| IMPUTE5 | https://www.dropbox.com/scl/fo/ukwimchnvp3utikrc3hdo/AKqYvE6-9C5kLpKDSfhR8xQ?rlkey=n2zty39bdst5j5tycd0sf89ee&e=1&dl=0 | v1.2.0 |
| GREAT | https://bioconductor.org/packages/release/bioc/html/rGREAT.html | v.2.13.1 |
| Snakemake (*97*) | https://snakemake.readthedocs.io/en/stable/ | v7.31.0 |

**Table S4.** Percentage of TRACE detected segments overlap with hmmix and IBDmix outcomes.

| | | hmmix NEA | hmmix DEN | hmmix | IBDmix NEA | IBDmix DEN | IBDmix |
|---|---|---|---|---|---|---|---|
| wAFR | TRACE NEA | -- | -- | -- | 44.9% | 16.0% | 46.4% |
| | TRACE Ghost | -- | -- | -- | 5.1% | 3.6% | 7.6% |
| | Total | -- | -- | -- | 8.1% | 5.4% | 11.1% |
| eAFR | TRACE NEA | -- | -- | -- | 57.7% | 21.3% | 58.5% |
| | TRACE Ghost | -- | -- | -- | 5.5% | 3.7% | 27.9% |
| | Total | -- | -- | -- | 10.6% | 6.5% | 13.7% |
| EUR | TRACE NEA | 83.0% | 0.02% | 83.3% | 71.2% | 14.8% | 81.3% |
| | TRACE Ghost | 2.6% | 1.5% | 18.9% | 4.8% | 1.8% | 7.4% |
| | Total | 50.3% | 1.4% | 57.6% | 44.4% | 10.4% | 52.1% |
| EAS | TRACE NEA | 91.5% | 0.2% | 91.8% | 74.4% | 13.6% | 84.1% |
| | TRACE DEN | 1.5% | 68.1% | 71.6% | 15.7% | 44.6% | 59.7% |
| | TRACE Ghost | 2.2% | 2.1% | 18.5% | 5.0% | 2.5% | 7.9% |
| | Total | 51.8% | 5.0% | 62.4% | 44.1% | 11.4% | 53.5% |
| SAS | TRACE NEA | 90.3% | 0.1% | 90.7% | 75.4% | 18.0% | 84.3% |
| | TRACE DEN | 2.2% | 75.5% | 80.0% | 15.3% | 49.4% | 64.5% |
| | TRACE Ghost | 2.6% | 3.1% | 20.7% | 5.6% | 2.4% | 8.7% |
| | Total | 51.3% | 6.9% | 64.2% | 45.2% | 14.8% | 55.1% |

**Table S5.** Performance of classification rules in simulations. See Section S4.3 for more details.

| | | Neanderthal | | Denisovan | | Ghost | |
|---|---|---|---|---|---|---|---|
| | | Precision | Recall | Precision | Recall | Precision | Recall |
| *Rule Group 1* | $P_{ND00} > 0.8$ | 0.7% | 1.4% | 4.4% | 5.2% | **91.9%** | **97.2%** |
| | $P_{ND10} > 0.2$ & $P_{ND10} > P_{ND01}$ | **93.6%*** | **97.4%** | 6.0% | 3.4% | 0.5% | 0.2% |
| | $P_{ND01} > 0.2$ & $P_{ND01} > P_{ND10}$ | 0.1% | 0.1% | **99.6%** | **79.0%** | 0.3% | 0.2% |
| *Rule Group 2* | $P_{ND00} > 0.8$ & $P_{sYRI} > 0.1$ | 0.7% | 1.3% | 3.1% | 3.1% | **93.0%** | **85.0%** |
| | $(P_{ND00} > 0.8 \vert P_{sYRI} > 0.1)$ & $P_{ND10} > P_{ND01}$ | **84.3%** | **98.4%** | 8.7% | 5.5% | 6.9% | 3.9% |
| | $(P_{ND00} > 0.8 \vert P_{sYRI} > 0.1)$ & $P_{ND01} > P_{ND10}$ | 0.1% | 0.2% | **96.1%** | **89.0%** | 3.8% | 3.2% |

*Bold records reflect the performance statistics for the archaic ancestry that the rule is aimed to detect.

**Table S6.** Inferred divergence times between archaics and modern human using upper-end coalescence times of introgression branches.

| | Neanderthal | Denisovan | Ghost |
|---|---|---|---|
| wAFR | 32444 (22372, 71747)* | -- | 29025 (21416, 43946) |
| eAFR | 31434 (22113, 63665) | -- | 28888 (21410, 43489) |
| EUR | 29547 (22215, 42161) | -- | 30159 (22006, 46220) |
| EAS | 29570 (22686, 42886) | 34066 (22750, 122768) | 30668 (22066, 46959) |
| SAS | 29615 (22272, 43137) | 30492 (21880, 48004) | 30221 (22033, 47795) |
| All | 29749 (22373, 43885) | 32289 (22299, 53976) | 29626 (21656, 45288) |

* Showing mean and 2.5 and 97.5 percentiles (in brackets) across archaic segments

**Table S7**. Performance of methods in simulations under OOA model and OOA + Ghost model.

| | | OOA model | | | OOA + Ghost model | | |
|---|---|---|---|---|---|---|---|
| | | TRACE | hmmix | IBDmix | TRACE | hmmix | IBDmix |
| NEA (2%)[†] | precision | 96.5% | 91.5% | 96.8% | 93.4% | 81.3% | 96.1% |
| | recall | 52.9% | 54.8% | 78.1% | 50.4% | 55.0% | 78.8% |
| DEN (0.1%)[†] | precision | 72.8% | 44.6% | 17.6% | 62.5% | 17.0% | 20.3% |
| | recall | 31.1% | 34.3% | 35.9% | 25.5% | 33.5% | 41.7% |
| Ghost (0 / 9%)[+] | precision | (0.07%)* | – | – | 94.1% | – | – |
| | recall | | | | 16.4% (1.5%)* | | |
| Total | precision | 84.6% | 83.6% | 97.1% | 94.8% | 84.3% | 97.2% |
| | recall | 53.8% | 54.8% | 76.3% | 21.7% | 9.9% | 12.5% |
| | Archaic per genome | 0.9% | 1.0% | 1.2% | 2.4% | 1.2% | 1.2% |

*Identified ghost ancestry per genome.
[†]Simulated introgression proportion.

**Table S8**. Percentage of TRACE detected segments overlap with hmmix and IBDmix outcomes in simulations: OOA model & OOA + Ghost model.

| | | hmmix NEA | hmmix DEN | hmmix | IBDmix NEA | IBDmix DEN | IBDmix |
|---|---|---|---|---|---|---|---|
| OOA model | TRACE NEA | 83.0% | 0.1% | 83.2% | 96.1% | 16.5% | 99.6% |
| | TRACE DEN | 0.8% | 1.0% | 20.4% | 0.8% | 0.2% | 18.1% |
| | TRACE | 71.8% | 3.8% | 77.4% | 83.3% | 16.4% | 92.4% |
| OOA + Ghost model | TRACE NEA | 83.1% | 0.2% | 84.2% | 95.6% | 17.2% | 99.8% |
| | TRACE DEN | 4.0% | 4.2% | 39.7% | 1.6% | 0.2% | 26.7% |
| | TRACE | 30.4% | 4.1% | 55.3% | 33.0% | 7.0% | 52.6% |

**Table S9**. Correlation between ghost ancestry and B-score / recombination rate.

| | | B-score | | Recombination rate | |
|---|---|---|---|---|---|
| | | Spearman's rho | P-value | Spearman's rho | P-value |
| wAFR | Neanderthal | 0.1108 | < 2.2e-308 | 0.0862 | < 2.2e-308 |
| | Ghost | 0.3810 | < 2.2e-308 | 0.2863 | < 2.2e-308 |
| eAFR | Neanderthal | 0.1246 | < 2.2e-308 | 0.0899 | < 2.2e-308 |
| | Ghost | 0.3416 | < 2.2e-308 | 0.2529 | < 2.2e-308 |
| EUR | Neanderthal | 0.1856 | < 2.2e-308 | 0.1405 | < 2.2e-308 |
| | Ghost | 0.2094 | < 2.2e-308 | 0.1561 | < 2.2e-308 |
| EAS | Neanderthal | 0.1747 | < 2.2e-308 | 0.1314 | < 2.2e-308 |
| | Denisovan | 0.0795 | < 2.2e-308 | 0.0599 | < 5.0e-299 |
| | Ghost | 0.2097 | < 2.2e-308 | 0.1586 | < 2.2e-308 |
| SAS | Neanderthal | 0.2113 | < 2.2e-308 | 0.1492 | < 2.2e-308 |
| | Denisovan | 0.1019 | < 2.2e-308 | 0.0630 | < 2.1e-252 |
| | Ghost | 0.2074 | < 2.2e-308 | 0.1553 | < 2.2e-308 |
| All | Neanderthal | 0.2756 | < 2.2e-308 | 0.2008 | < 2.2e-308 |
| | Denisovan | 0.1252 | < 2.2e-308 | 0.0837 | < 2.2e-308 |
| | Ghost | 0.4468 | < 2.2e-308 | 0.3395 | < 2.2e-308 |

**Table S10.** Frequencies of ancestries in previously reported Neanderthal and Denisovan shared deserts.

| chromosome | start | end | NEA frequency* (%) | DEN frequency* (%) | Ghost frequency† (%) |
|---|---|---|---|---|---|
| 1 | 99000000 | 112000000 | 0.068 | 0.049 | 0.757 |
| 3 | 78000000 | 90000000 | 0.0006 | 0.002 | 0.414 |
| 7 | 113000000 | 124000000 | 0.004 | 0.002 | 0.227 |
| 8 | 54500000 | 65400000 | 0.023 | 0.185 | 0.622 |
| 13 | 49000000 | 61000000 | 0.026 | 0.016 | 0.281 |

* Frequencies in non-African populations (EUR, EAS, SAS).
† Frequencies in all populations (wAFR, eAFR, EUR, EAS, SAS).

**Table S11**. Neanderthal ancestry deserts identified using TRACE which are shared across *all* non-African populations. Previous publications referring to observed deserts are reported as references.

| Chromosome | Start | End | References |
|---|---|---|---|
| chr1 | 61436000 | 78436000 | |
| chr1 | 87036000 | 97136000 | |
| chr1 | 100436000 | 114436000 | (*52*, *63*) |
| chr2 | 196434000 | 210934000 | |
| chr3 | 76117000 | 90217000 | (*28*, *52–54*, *63*) |
| chr3 | 176017000 | 186217000 | |
| chr4 | 65999000 | 75199000 | |
| chr5 | 80392000 | 95192000 | (*53*, *54*) |
| chr5 | 130392000 | 140392000 | |
| chr7 | 105509000 | 128109000 | (*28*, *52–54*, *63*) |
| chr8 | 52714000 | 64814000 | (*28*, *54*, *63*) |
| chr8 | 89714000 | 97214000 | |
| chr8 | 104714000 | 114814000 | (*53*) |
| chr10 | 95617000 | 107617000 | |
| chr13 | 48774000 | 61874000 | (*52*) |
| chr18 | 27052000 | 45152000 | (*53*, *54*, *63*) |

**Table S13**. Percentage of TRACE detected segments overlap with hmmix and IBDmix outcomes in Oceanians.

| | hmmix NEA | hmmix DEN | hmmix | IBDmix NEA | IBDmix DEN | IBDmix |
|---|---|---|---|---|---|---|
| TRACE NEA | 86.9% | 3.0% | 90.4% | 79.1% | 22.2% | 80.4% |
| TRACE DEN | 2.0% | 94.1% | 96.7% | 23.2% | 65.6% | 68.5% |
| TRACE Ghost | 4.0% | 15.0% | 32.0% | 8.2% | 6.6% | 12.1% |
| Total | 37.7% | 40.4% | 81.4% | 43.8% | 35.9% | 62.4% |

**Table S14**. Performance of methods in simulations under No Super-Archaic model and Super-Archaic model.

| | | No Super-Archaic model | | | | Super-Archaic model | | | |
|---|---|---|---|---|---|---|---|---|---|
| | | TRACE (t=15000) | TRACE (t=10000) | hmmix | IBDmix | TRACE (t=15000) | TRACE (t=10000) | hmmix | IBDmix |
| NEA (2%)[†] | precision | 73.8% | 71.0% | 62.5% | 72.6% | 69.3% | 68.4% | 58.9% | 72.3% |
| | recall | 38.9% | 50.1% | 69.3% | 78.6% | 38.9% | 50.7% | 70.9% | 77.0% |
| DEN (3%)[+] | precision | 92.9% | 90.8% | 81.0% | 90.3% | 90.4% | 89.1% | 80.8% | 88.9% |
| | recall | 40.7% | 50.3% | 69.9% | 48.9% | 37.7% | 46.8% | 65.3% | 38.8% |
| Ghost (9%)[+] | precision | 91.7% | 86.3% | – | – | 91.1% | 86.3% | – | – |
| | recall | 11.0% (1.1%)* | 12.6% (1.3%)* | | | 10.6% (1.0%)* | 12.3% (1.3%)* | | |
| Total | precision | 96.2% | 93.7% | 91.7% | 98.1% | 96.4% | 94.2% | 92.5% | 98.0% |
| | recall | 23.1% | 27.4% | 29.6% | 23.0% | 22.9% | 27.1% | 29.5% | 20.7% |
| | Archaic per genome | 3.3% | 4.0% | 4.6% | 3.0% | 3.3% | 4.0% | 4.6% | 2.8% |

*Identified ghost ancestry per genome.
[†]Simulated introgression proportions.

**Table S15.** Percentage of TRACE detected segments overlap with hmmix and IBDmix outcomes in simulations: No Super-Archaic model & Super-Archaic model.

| | | hmmix NEA | hmmix DEN | hmmix | IBDmix NEA | IBDmix DEN | IBDmix |
|---|---|---|---|---|---|---|---|
| No Super-Archaic model | TRACE NEA | 94.7% | 3.7% | 98.3% | 87.7% | 19.3% | 99.2% |
| | TRACE DEN | 2.0% | 97.2% | 99.4% | 18.1% | 78.0% | 99.2% |
| | TRACE Ghost | 9.7% | 16.2% | 72.0% | 1.6% | 2.7% | 31.4% |
| | TRACE | 27.6% | 44.0% | 88.9% | 28.9% | 35.3% | 73.9% |
| Super-Archaic model | TRACE NEA | 94.1% | 4.2% | 98.7% | 88.1% | 20.7% | 98.8% |
| | TRACE DEN | 2.7% | 96.7% | 99.2% | 19.0% | 72.4% | 99.6% |
| | TRACE Ghost | 12.2% | 21.0% | 76.0% | 2.9% | 3.6% | 34.1% |
| | TRACE | 30.1% | 42.8% | 89.8% | 30.5% | 31.8% | 73.8% |

**Table S16.** Proportion of Denisovan segments with super-archaic ancestry classified as "Denisovan" under different classification rules in simulations under Super-Archaic model.

| | Super-Archaic classified as DEN | DEN precision | DEN recall |
|---|---|---|---|
| $P_{ND10} > 0.2$ & $P_{ND10} < P_{ND01}$ | 39.8% | 99.6% | 66.0% |
| $P_{ND00} \leq 0.8$ & $P_{ND10} < P_{ND01}$ | 53.9% | 99.0% | 77.5% |
| ($P_{ND00} \leq 0.8 \| P_{sYRI} \leq 0.1$) & $P_{ND10} < P_{ND01}$ | 71.7% | 93.9% | 82.7% |

**Table S17.** Precision of super-archaic segments detection in simulations and proportions of Denisovan ancestry identified as super-archaic in simulations and real data.

| | Precision | Proportion of genomewide Denisovan ancestry called as super-archaic | Proportion of Denisovan ancestry per individual called as super archaic |
|---|---|---|---|
| Super-Archaic simulation | 70.2% (95% CI: 66.4%-74.4%) | 1.73% | 1.4% |
| Real data OCN | – | 0.3% | 0.3% |
